# Convergent and non-additive impact of schizophrenia risk genes in human neurons

**DOI:** 10.1101/2022.03.29.486286

**Authors:** PJ Michael Deans, Kayla G. Townsley, Aiqun Li, Carina Seah, Jessica Johnson, Judit Garcia Gonzalez, Evan Cao, Nadine Schrode, Alex Yu, Sam Cartwright, Georgios Voloudakis, Wen Zhang, Minghui Wang, John F. Fullard, Kiran Girdhar, Eli Stahl, Schahram Akbarian, Bin Zhang, Panos Roussos, Paul O’Reilly, Laura M. Huckins, Kristen J. Brennand

## Abstract

Genetic studies of schizophrenia reveal a complex polygenic risk architecture comprised of hundreds of risk variants; most are common in the population at-large, non-coding, and act by genetically regulating the expression of one or more gene targets (“eGenes”). It remains unclear how genetic variants predicted to confer individually small effects combine to yield substantial clinical impacts in aggregate. Here, we demonstrate that eGenes have shared downstream transcriptomic effects (“convergence”) that may underlie unexpected interactions (“non-additive effects”) observed when eGenes are manipulated in combination. We apply a pooled CRISPR approach to perturb schizophrenia eGenes in human induced pluripotent stem cell-derived glutamatergic neurons. The strength and specificity of convergence increased between functionally similar eGenes. Predicting that convergence might impact additive relationships between risk loci when inherited together, we use an arrayed approach to explore bidirectional combinatorial perturbations of a partially overlapping set of fifteen schizophrenia eGenes. When specifically considering groups of synaptic or epigenetic eGenes, combinatorial eGene perturbations yield changes that are smaller than predicted by summing individual eGene effects (“sub-additive effects”). Moreover, convergent and non-additive downstream transcriptomic effects overlap, suggesting that functional redundancy of eGenes may be a major mechanism underlying non-additivity. Combinatorial perturbations result in outcomes that are not yet well-predicted by single eGene perturbations alone, indicating that the effects of polygenic risk cannot necessarily be extrapolated from experiments testing one risk gene at a time.

## INTRODUCTION

The genetic architecture of schizophrenia is complex and polygenic. Highly penetrant rare mutations underlie only a fraction of cases^1^. Rather, genome wide association studies (GWAS) indicate that schizophrenia is predominantly associated with genetic variation that is common in the population^2^. These risk loci have small effect sizes, are typically found in non-coding regions, and regulate the expression of one or more genes^3–5^. Mapping GWAS loci to their target genes (termed “eGenes”, as defined by significant genetic regulation of expression) remains challenging, but is informed by expression quantitative trait loci (eQTL)^6–9^, chromatin accessibility^10–12^, enhancers^13–16^, and 3D chromatin architecture^17–22^. The regulatory activity of risk loci can be empirically evaluated using massively parallel reporter assays^23–25^ and pooled CRISPR screens^26^, and causal gene targets and functions definitively resolved by genetic engineering in human induced pluripotent stem cells (hiPSCs)^10,11,17,27,28^.

Schizophrenia eGenes are particularly expressed during fetal cortical development^29–31^ and in glutamatergic neurons (as well as medium spiny neurons, and certain interneurons)^31–33^. They are highly co-expressed in human brain tissue^34^ and cultured neurons^17^, show high connectivity in protein-protein interaction networks^17,35–37^, and are enriched for roles in synaptic function and gene regulation^2,17,38–42^. Likewise, transcriptomic studies of schizophrenia post-mortem brains also identify aberrant expression of genes associated with synaptic function and chromatin dynamics in neurons^43–45^. The mechanism by which hundreds of distinct eGenes lead to shared molecular pathology is unknown.

We predicted that eGenes linked to schizophrenia would share substantial downstream transcriptomic changes with a common direction of effect (termed “convergence”). Although convergence has already been described in the context of loss-of-function autism spectrum disorder risk genes^46–56^, these rare mutations almost never co-occur in the same individual. The convergent impact of common variants − which are frequently inherited together, and the impacts of which are apparent only in aggregate − remain unknown. We targeted twenty schizophrenia eGenes in iGLUTs using a pooled CRISPRa approach, successfully perturbed ten, and resolved their convergent impact (*CALN1, CLCN3, FES, NAGA, PLCL1, TMEM219*, *SF3B1, UBE2Q2L, ZNF823, ZNF804A*). To test if convergence impacts additive relationships between eGenes inherited in combination (i.e. if eGene effects sum linearly according to the additive model^26^), we manipulated eGenes one at a time and together in biological groups defined by known roles at the synapse (“synaptic”: *SNAP91, CLCN3, PLCL1, DOC2A, SNCA*), or regulating transcription (“regulatory”: *ZNF823, INO80E, SF3B1, THOC7, GATAD2A*), or with un-related non-synaptic, non-regulatory biology (“multi-function”: *CALN1, CUL9, TMEM219, PCCB, FURIN),* and random combinations thereof. Combinatorial eGene perturbations within biological functions resulted in transcriptomic changes that were predominantly smaller than predicted by the additive model (“sub-additive”); sub-additive genes overlapped with convergent genes of the same eGenes.

Our work begins to address the long-standing question of how risk variants with individually minuscule effects combine to yield substantial impacts on neuronal biology. Overall, convergent signatures were robust, observable across three partially overlapping lists of schizophrenia eGenes, whether manipulated in arrayed or pooled approaches. Altogether, our work suggests that combinatorial interactions between eGenes^57,58^ should be empirically measured rather than predicted from single eGene studies.

## RESULTS

### Convergence of downstream transcriptomic impacts across schizophrenia eGene perturbations

We^27,59–61^ and others^11,62–68^ demonstrated that iGLUTs are >95% glutamatergic neurons, robustly express glutamatergic genes, release neurotransmitters, produce spontaneous synaptic activity, and recapitulate the impact of psychiatric trait associated genes. iGLUTs express most schizophrenia eGenes, including all eGenes prioritized herein^27^.

eGenes whose brain expression was predicted to be up-regulated by GWAS loci^2^ were prioritized for a pooled CRISPR activation (CRISPRa) experiment, which are currently restricted to one direction of effect. eGenes that were non-coding, located in the MHC locus, or poorly expressed in iGLUTs were excluded. First, transcriptome and epigenome imputation (EpiXcan^69^) of schizophrenia GWAS^2^ risk loci from post-mortem brain^43,70^ prioritized seven schizophrenia eGenes (**SCZ1**: *NEK4, PLCL1, UBE2Q2L, NAGA, FES, CALN1,* and *ZNF804A*) (**Table 1A**; **Fig. 1A**). Second, transcriptomic imputation (prediXcan^71–73^, p<6×10^-6^) of SCZ GWAS^2^ identified ∼250 eGenes (**SI Table 1)**, subsequently narrowed by considering colocalization (COLOC^74,75^, PP4 > 0.8) between schizophrenia GWAS^2^ and post-mortem brain expression quantitative loci (eQTL) peaks^6^, which identified 25 eGenes (**SI Table 1)**. 22 eGenes overlapped between approaches, ten of which were coding genes associated with increased expression in schizophrenia (**SCZ2**: *CALN1*, *CLCN3, CUL9, DOC2A, PLCL1, INO80E, SF3B1, SNAP91, TMEM219, ZNF823)* (**Table 1B**; **Fig. 1A**). Of note, our eGene selection, derived in bulk post-mortem brain, is largely preserved using an excitatory neuron-specific PrediXcan analysis (ExN-PrediXcan, **Tables 1A and 1B**).

**Figure 1.**
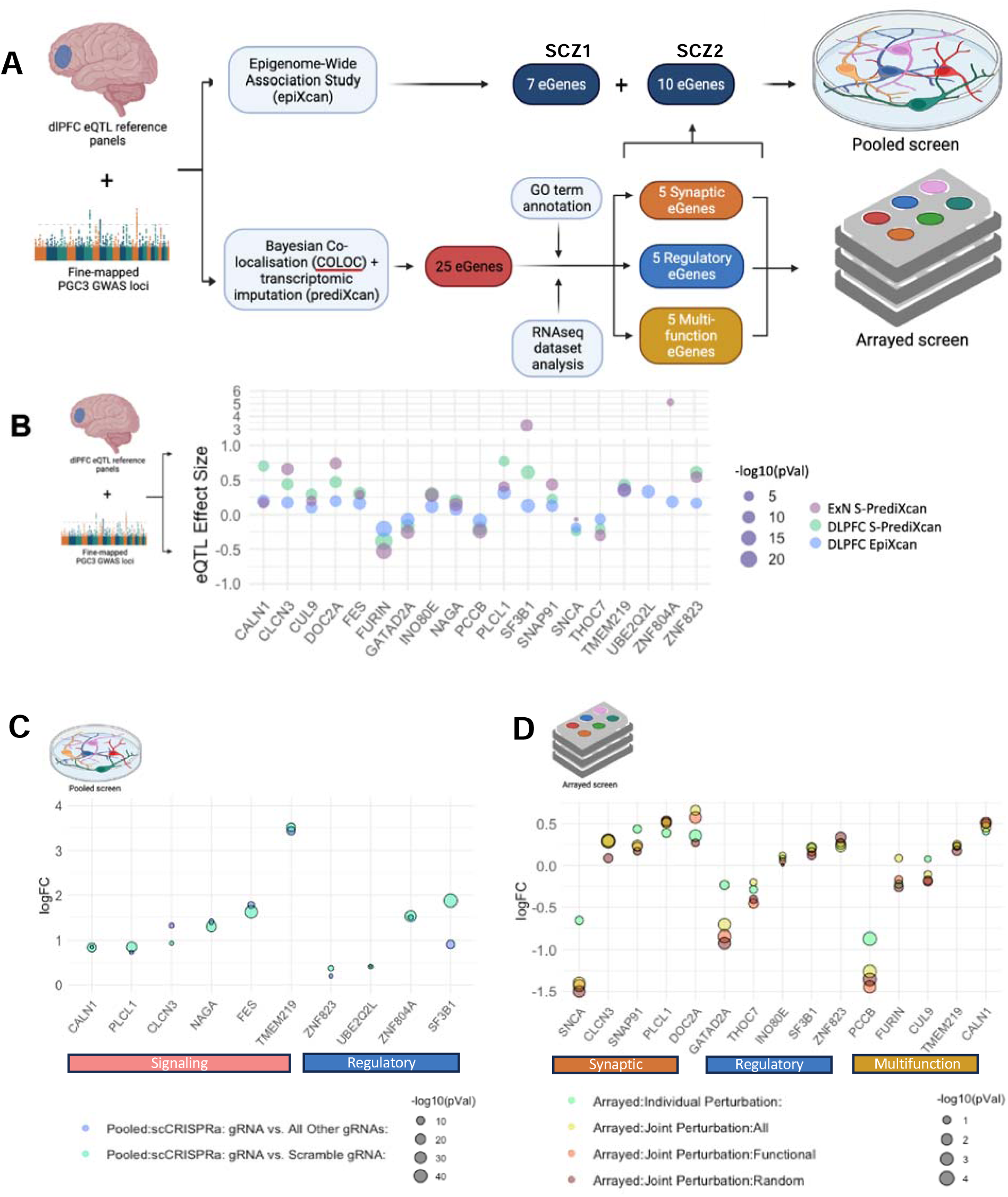
Prioritization and manipulation of synaptic, regulatory, and multi-function brain eGenes regulated by schizophrenia. **A.** Schematic of schizophrenia eGene identification and prioritization. Schizophrenia eGenes were prioritized by fine-mapping (COLOC), transcriptomic imputation (PrediXcan), and/or epigenomic imputation (EpiXcan) schizophrenia GWAS using post-mortem brain expression data. **B.** Effect sizes of significant eGenes from either dorsolateral prefrontal cortex (DLPFC) EpiXcan (blue), DLPFC S-PrediXcan (green) or excitatory neuron (ExN) S-PrediXcan (purple) transcriptomic imputation studies. Size of circles corresponds with the -log10(adjusted p-value) **C.** Log2(fold change) of all eGenes in the pooled experiments **SCZ1** and **SCZ2** comparing all perturbed cells of one target eGene identity to all other cells of different eGene identities (blue) or compared to only Scramble gRNA (teal). Size of circles corresponds with the -log(adjusted p-value) from a one-way pairwise Wilcox Rank Sum. **D.** Log2(fold change) of all eGenes in the arrayed experiment following single (teal) and joint perturbations across all 15 eGenes (yellow) or functional (orange) or random (maroon) sets of 5 eGenes in D21 hiPSC-NPC derived iGLUTs, using individual vectors. Size of circles corresponds with the - log(adjusted p-value) from a one-tailed t-test.

**Table 1A.** Top PGC3 SCZ-GWAS eGenes, prioritized by EpiXcan and epigenetic annotation, as epigenetic/regulatory (blue) and signaling (red) for pathway studies. *NEK4*, *PLCL1*, *UBE2Q2L*, *NAGA*, *FES*, *CALN1*, and *ZNF804A* are present in the pooled screen **SCZ1.**

|  |  |  |  |  |  |  |  |  |  |  |  |
| --- | --- | --- | --- | --- | --- | --- | --- | --- | --- | --- | --- |
|  |  |  | EpiXcan |  | ExN-PrediXcan |  |  | Epigenetic Annotation |  |  |  |
| Group | Chr | Gene | z | r2 | z | effect_size | P | ATACseq | H3K27ac | H3K4me3 | H3K27ac (neuron) |
| regulatory | 3 | *NEK4* | 8.64 | 0.26 | NA | NA | NA | NA | Promoter | NA | Promoter |
| signaling | 2 | *PLCL1* | 7.2 | 0.04 | 5.214354 | 0.403086 | 1.84E-07 | NA | Intron | Promoter | Intron |
| regulatory | 15 | *UBE2Q2L* | 6.89 | 0.02 | NA | NA | NA | NA | Promoter | Promoter | Promoter |
| signaling | 22 | *NAGA* | 6.67 | 0.25 | 6.983019 | 0.148533 | 2.89E-12 | NA | Promoter | Promoter | NA |
| signaling | 15 | *FES* | 6.63 | 0.05 | 3.917081 | 0.289907 | 8.96E-05 | NA | Promoter | Promoter | NA |
| signaling | 7 | *CALN1* | 6.57 | 0.07 | 5.356091 | 0.171432 | 8.50E-08 | NA | Intron | Promoter | Intron |
| regulatory | 2 | *ZNF804A* | 6.27 | 0.05 | 2.867957 | 5.119801 | 4.13E-03 | NA | Promoter | Intron | NA |

**Table 1B.**
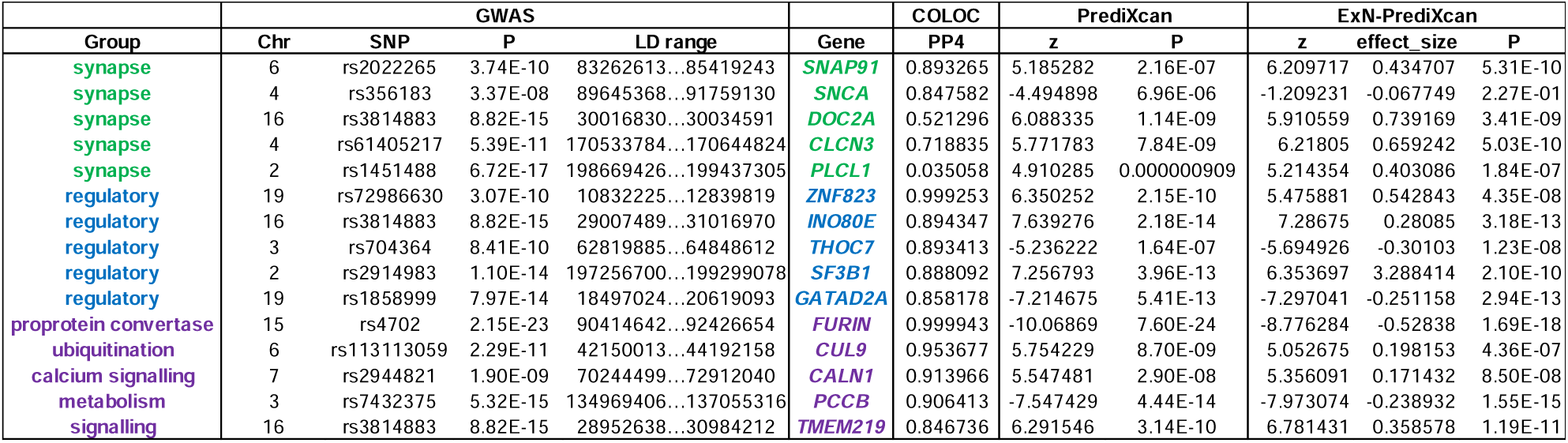
Top PGC3 SCZ-GWAS eGenes, prioritized by COLOC and PrediXcan, as synaptic (green), regulatory/epigenetic (blue), and multi-function (purple). *CALN1*, *CLCN3*, *CUL9*, *DOC2A*, *PLCL1*, *INO80E*, *SF3B1*, *SNAP91*, *TMEM219*, *ZNF823* are present in the pooled screen **SCZ2**. All 15 eGenes are present in the subsequent arrayed screen.

Pooled CRISPR screening combined single-cell RNA sequencing readouts and direct detection of sgRNAs^76^. Two independently designed, constructed, and validated pooled CRISPRa libraries (**SCZ1** and **SCZ2**) were transduced into iGLUTs from two donors in independent experiments at unique developmental time-points (DIV7 or DIV21, **SI Fig 1F**). Non-perturbed cells from both **SCZ1** and **SCZ2** demonstrated gene expression patterns which correlated with expression in the adult postmortem DLPFC in neurotypical controls (**SI Fig. 12A-C**) and cortical neurons. Specifically, these cells were most strongly correlated with fetal cells transitioning to neuronal fate, fetal excitatory neurons, and cortical adult neurons (**SI Fig. 13**). Pooled CRISPR screens may be confounded by non-cell autonomous effects, which is of particular concern in neurons, as maturation is necessarily activity-dependent and therefore requires normal functioning of nearby cells; here, this is partially mitigated by the large number of presumably wildtype neurons in the population expressing either a scramble gRNA or no detectable gRNA at all (>60% of all pooled cells, see **SI Fig. 4A**). There was no significant difference in the degree of variance in maturity of the cell population between experiments and imputed cell fractions were not correlated with perturbation status (**SI Fig. 2 and 3**). An unsupervised framework, Weighted Nearest Neighbor Analysis^77^, assigned successful perturbations; in total, we resolved perturbations of six of seven SCZ1 eGenes (ten gRNAs each) and four of ten SCZ2 eGenes (three gRNAs each). For 5401 and 6352 cells, respectively, we identified the sgRNA in each cell, the *cis* target gene with differential expression, and the downstream *trans* alterations to pathways resulting from initial *cis* up-regulation. Following QC, normalization, and removal of doublets (cells containing more than one sgRNA), an average of 316 cells per sgRNA were successfully perturbated (ranging from 93-552) for a total of 3640 perturbed cells and 210 scramble controls (**SI Fig. 4-7**). Upregulation of eGenes by CRISPRa ranged from 0.2 to 3 log2 fold-change (Log2FC), comparable to the predicted effect sizes [SCZ1 (0.08 to 0.35); SCZ2 (0.2 to 0.77)] and eGene expression changes (Log2FC range 0.3-5.2) in the post-mortem dorsolateral prefrontal cortex (**Fig. 1B, C**; **SI Table 2,3; Fig. 8**). Effects of different gRNAs targeting the same eGenes were highly concordant, even when the degree of perturbation varied (**SI Fig. 15**). Differentially expressed genes (DEGs, p_FDR_<0.05) were enriched for neuroactive ligand-receptor interaction, protein processing in the endoplasmic reticulum, proteasome, and spliceosome Gene Ontology and KEGG Pathways terms (**SI Data 3**), suggesting that diverse eGenes might impact similar neural processes and pathways.

We define “convergence” as the independent development of transcriptomic changes in the same direction resulting from all eGene perturbations. DEGs were meta-analyzed (using METAL^78^, p< 1.92E-06), and “convergent” genes were defined as those with shared direction of effect across all eGene perturbations and with non-significant heterogeneity between eGenes (Cochran’s heterogeneity Q-test p_Het_ > 0.05) (**Fig. 2**). Across all schizophrenia eGenes, 790 significantly down-regulated genes and 10 significantly up-regulated genes were identified (Bonferroni meta p-value<=0.05) (**SI Data 3**), enriched for brain development, neuronal morphology, signaling, and transcriptional regulation (**SI Data 3**).

**Figure 2.**
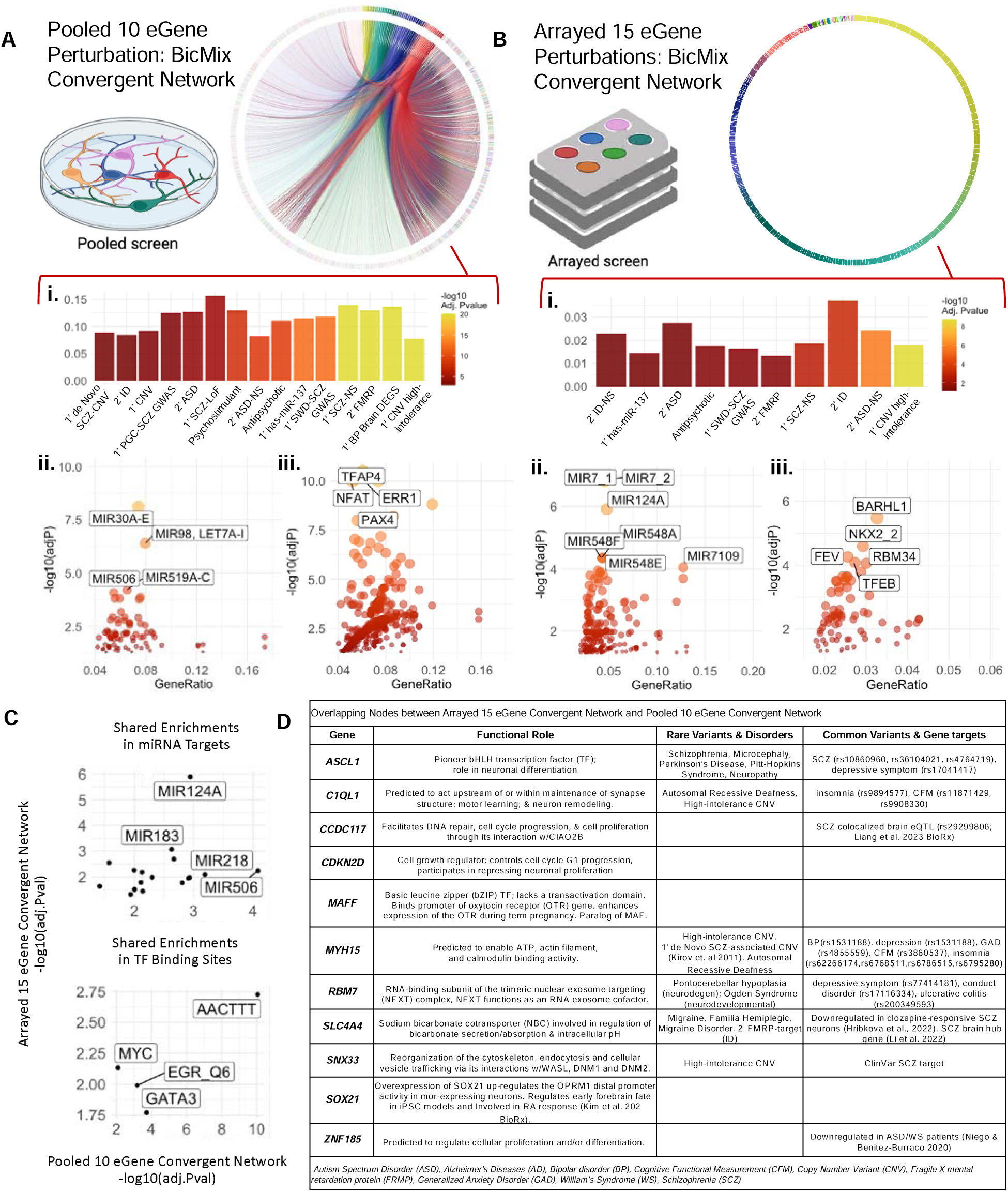
Downstream target-convergent networks identified by Bayesian bi-clustering resolve distinct networks enriched for schizophrenia common and rare variant target genes and transcription factor binding motifs. **A.** Convergent networks resolved across the downstream transcriptomic impacts of all ten target perturbations in the pooled screens **SCZ1** and **SCZ2** identified 1869 convergent genes with enrichments for **(i)** brain-related GWAS genes, **(ii)** transcription factor binding sites of know schizophrenia-associated TFs (TFAP4, NFAT and ERR1), and **(iii)** common and rare variant target genes. **B.** Convergent networks resolved across the downstream transcriptomic impacts of all fifteen target perturbations in the arrayed assay identified 255 convergent genes with enrichments for **(i)** miRNA targets and **(ii)** transcription factor binding sites of know schizophrenia-associated TFs (TFAP4, NFAT and ERR1), and **(iii)** common and rare variant target genes. **C.** While largely distinct, the resolved convergent networks from the arrayed and pooled screen shared 16 significant enrichments for miRNA targets and 4 significant enrichments for TF targets – many of which are thought to play a role in regulation of schizophrenia-associated genes. **D.** Overlapping nodes between the two networks were often involved in neuronal proliferation, and differentiation.

To identify groups of genes with similar expression patterns across eGene perturbations we define “convergent networks” as relationships between genes that are co-regulated by shared biological mechanisms. Unsupervised Bayesian bi-clustering^78^ and gene co-expression network reconstruction from the pooled CRISPRa single cell RNAseq (n=3850 cells, 16851 genes, donor/batch corrected and normalized to adjust for covariates such as cell heterogeneity) identified high-confidence co-expressed gene networks. Across the pooled single-cell experiments, 1048 protein-coding source node genes (>5 edges) were identified, with a total network membership of 1869 genes that clustered together in at least 20% of the runs (**Fig. 2A**, **SI Data 3**), and significant enrichments for gene targets of schizophrenia GWAS loci as well as transcription factors and miRNAs that regulate schizophrenia GWAS loci, such as *TCF4*^79,80^ (**Fig. 2A, i-iii)**. The cross-target convergent network was enriched in biological pathways implicated in schizophrenia etiology (**SI Fig. 9**); over representation analysis revealed schizophrenia, bipolar disorder, intellectual disability, and autism spectrum disorder common and rare risk genes to be significantly over-represented in node genes shared across all eGene perturbations (**Fig. 2A, i; SI Data 3**).

To study the strength and composition of convergent networks, we define “network convergence” as the sum of the network connectivity score (i.e., networks with fewer nodes and more interconnectedness have increased convergence). We endeavored to identify the biological factors (e.g., number of eGenes, functional similarity of eGenes, and eGene co-expression) that influenced network convergence. eGene number tested the number of eGenes used to generate a convergent network. Functional similarity (i.e., the degree of shared biological functions amongst eGenes) was calculated two ways: Gene Ontology semantic similarity scoring (within biological pathway, cellular component, and molecular function)^81^, and synaptic/signaling score (proportion of eGenes with annotated function as either “signaling” for pooled or “synaptic” for arrayed). The brain expression correlation was calculated as the strength of the correlation of eGene expression in the post-mortem dorsolateral prefrontal cortex^6^ (see **Methods**, **SI Fig. 10**). Bayesian reconstruction^82^ was performed across all random combinations of eGene perturbations from the pooled screen (1003 unique eGene-Convergent Network sets) and arrayed experiment described in the following section (32752 sets) and resolved distinct networks (**Fig. 3B,E**). Principal components analysis tested the effect of biological factors on the network convergence scores (**Fig. 3C-D, F-G**; **SI Fig. 10-11**). Only brain expression correlation and the proportion of synaptic/signaling genes were significantly positively correlated with network convergence across all sets in both the pooled [brain expression correlation: Pearson’s r=0.24, adj. p-value<0.001, signaling proportion: Pearson’s r=0.14, adj. p-value<0.01, n=826] and arrayed experiments [brain expression correlation: Pearson’s r=0.083, Bonferroni adjusted p-value<0.001, signaling proportion: Pearson’s r=0.25, adj. p-value<0.001, n=16319] (**Fig. 3D,G**). The average expression of perturbed eGenes was positively correlated with network convergence but is only significantly associated in the arrayed screen (**Fig. 3D, G**). Finally, although **SCZ1** and **SCZ2** pooled CRISPR screens were generated from distinct differentiation timepoints, the proportion of eGene perturbations by experiment did not correlate with the degree of network convergence, indicating that we have adequately controlled for variation in neuronal maturation (**Fig. 3D**; Pearson’s r=0.062, bonferroni p-value=1).

**Figure 3.**
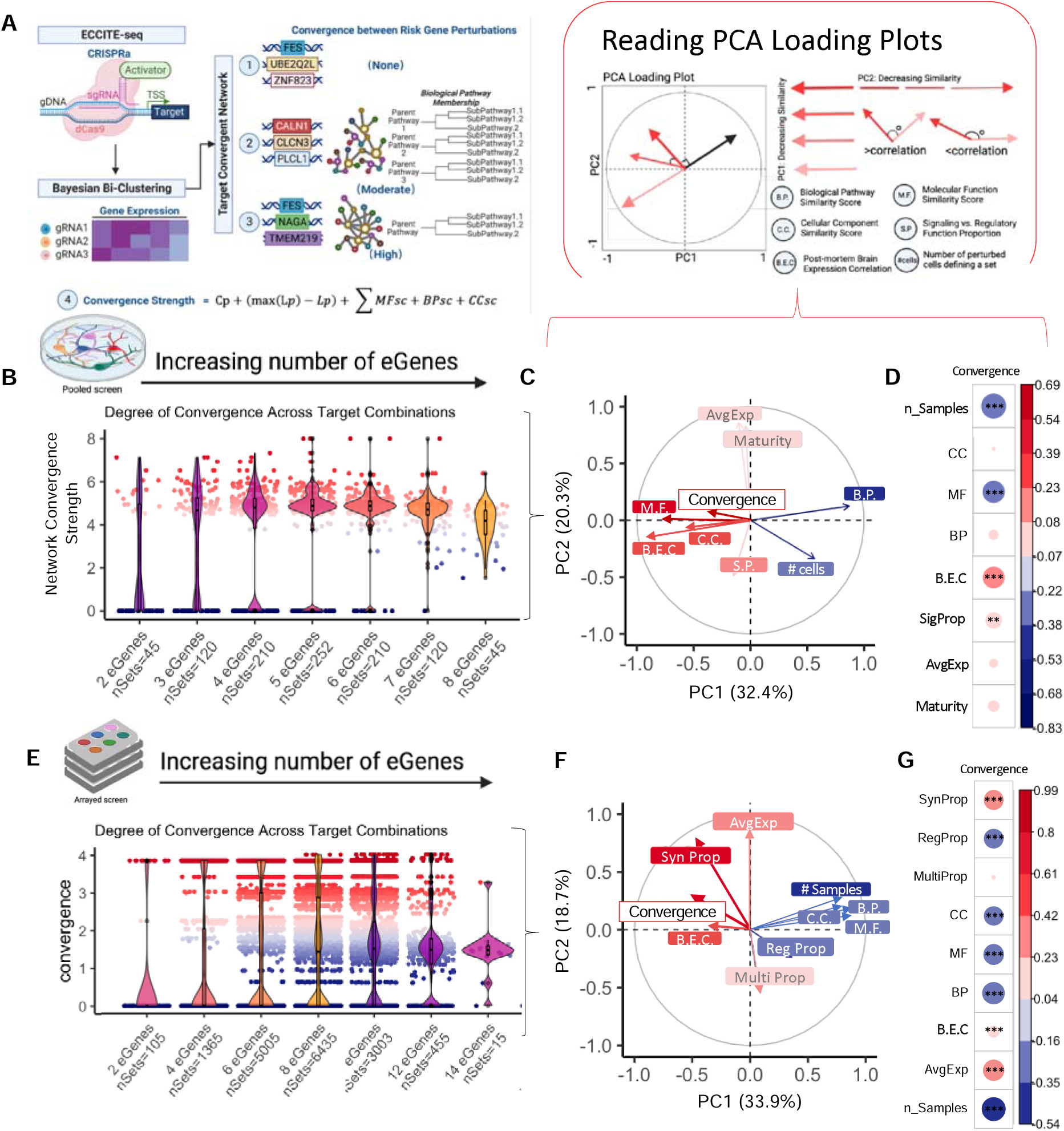
The degree of network convergence is influenced by functional similarity of target perturbations. **A.** Defining convergence and calculating convergent network strength. *<u>Here we define</u> <u>convergence as the independent development of transcriptomic similarities between</u> <u>separate “gene perturbations” that move towards union or uniformity of biological</u> <u>function</u>*. Bayesian bi-clustering identifies co-expressed genes that are shared across the downstream transcriptomic impacts of any given eGene perturbations, thus, the resolved networks are the transcriptomic similarities between distinct perturbations. While bi-clustering resolves convergent gene co-expression networks, the strength of convergence within a network can be defined by (i) the degree of network connectivity as define by two small-world network connectivity coefficients (edge density and average path length) and (ii) the degree of functional similarity or unity between genes represented within the network. Given this definition, **(1)** represents perturbations with no convergent downstream effects, **(2)** represents a network with a moderate degree of convergence because it (i) has resolved gene co-expression clusters that can be constructed into a network, (ii) has a moderate degree of network connectivity and (ii) is enriched in biological pathways with some redundancy, while **(3)** represents a highly convergent network because the degree of network connectivity is stronger and there is greater uniformity in biological pathway gene membership. Overall, we quantify the strength or degree of convergence using the function in **(4)**, where *Cp* is the edge density (the proportion of edges present given all possible edges) and *Lp* is the average path length (the mean of the shortest distance between each pair of nodes), *MFsc* is the average semantic similarity score between each pair of nodes in the network based on Molecular Function Gene Ontology, *BPsc* is the average semantic similarity score based on Biological Pathway Gene Ontology and *CCsc* is the average semantic similarity score based on Cellular Component Gene Ontology. Semantic similarity is based on the idea that genes with similar function have similar Gene Ontology annotations. Semantic similarity scores were calculated by aggregating four information content-based methods and one graph structure-based method with the R package GoSemSim. Created with BioRender.com. **B-G.** Principal component analysis (PCA) of the convergence scores, the three Gene Ontology scores (M.F., B.P., C.C.), brain expression correlation (B.E.C), and sample size across all resolved networks in both the pooled and arrayed assays revealed that some functional scores have similar influence on variance as convergence (**SI Fig. 19**). **B & E.** Distribution of the degree of convergence (x-axis) of networks across all possible combinations of 2 to 8 (y-axis; number of sets tested within each set) target perturbations from the single-cell pooled screen (**B**) and **(E)** across all possible combinations of 2 to 14 target perturbations from the arrayed screen show an influence of sample size on the ability to resolve a network. **C & F.** For both the pooled screen (**C**) and the **(F)** arrayed screen, PCs 1 (x-axis) and 2 (y-axis) explain ∼62% of the variance between networks. PC loadings demonstrate the influence of each variable on the variance between networks; within the first two PCs the influence of brain expression correlation (B.E.C) and proportion of signaling genes perturbed (S.P) on PCs 1 and 2 on variance explained are more strongly related to convergence degree compared to other functional scores. Since degree of convergence is influenced by number of eGene perturbed we ran PCA analysis within networks of the same set size and found that the pattern of influence of signaling proportion and brain expression correlation is maintained when convergence is ranked within set size shown in **SI Fig. 20**. **D & G.** This corresponds to an overall significant positive correlation between network convergence degree, signaling/synaptic proportion of perturbed genes in a set, and brain expression correlation between genes in a set (Bonferroni adjusted p-value of Pearson’s correlations: **\***<0.05, **\*\***<0.01, **\*\*\***<0.001.

### Combinatorial perturbation of schizophrenia eGenes results in sub-additive impacts on transcription that reflect convergence

To test if convergence between eGenes indeed impacts additive relationships between risk variants, we manipulated eGenes in combination to better approximate the polygenic nature of schizophrenia. Given that genes implicated in synaptic biology and epigenetic/transcriptional regulation are enriched for the schizophrenia risk^2,17,38–42^, we sought to generate three groups of eGenes, linked to synaptic biology, gene regulation, or neither (**Fig. 1A**, arrayed screen). Unconstrained by the unidirectionality of pooled CRISPR screens, we did not restrict our list to eGenes with a single direction of effect. From the 18 coding genes prioritized by the intersection of transcriptomic imputation and colocalization, eGenes were separated into discrete functional categories based on gene ontology annotations. Our final gene list included five synaptic genes (*SNAP91, CLCN3, PLCL1, DOC2A, SNCA*), five regulatory genes (*ZNF823, INO80E, SF3B1, THOC7, GATAD2A*), and five genes with non-synaptic, non-regulatory functions, termed “multi-function” (*CALN1, CUL9, TMEM219, PCCB, FURIN*) (**Table 1B**; **Fig. 1A**).

We applied an arrayed design (i.e., distinct conditions in each well) to manipulate schizophrenia eGenes alone and in combination, allowing us to capture cell autonomous and non-cell autonomous effects in a manner not possible in the pooled design (**Fig. 4, SI Table 4, SI Fig. 19**). Endogenous expression was increased and decreased (via CRISPRa and shRNAs, respectively) in the direction associated with schizophrenia risk. Three to five vectors per gene were tested in 7-day-old (D7) iGLUTs, identifying the single vector that best achieved the level of significant perturbation predicted by eQTL analyses as confirmed by qPCR (**SI Fig. 1E**). Each eGene was perturbed in 21-day-old (D21) iGLUTs for 72 hours (**Fig. 1D, SI Fig. 1F-G, SI Fig. 19A, SI Fig. 21A**), individually and jointly, including appropriate vector and scrambled controls, from two neurotypical donors with average polygenic risk scores (one experimental batch per donor). Three groups of five random genes, one group of ten random genes, and one group of all fifteen genes were also included. Significant (p<0.05) changes in eGene expression in iGLUTs were confirmed by RNAseq in 13/15 eGenes (**SI Fig. 1G, SI Fig. 19A**); we validated the magnitude and direction of experimental eGene perturbation relative to the dosage effects of the top predicted causal SNPs (e.g. eQTL effect size) and predicted eGene expression changes (**Fig. 1B, D; SI Tables 2,3,4**). Across donors, donor status did not significantly impact the degree of eGene perturbation (**SI Fig. 1H**, p=0.75, paired t test). Single perturbation of eGenes by CRISPRa ranged from 0.07 to 0.44 log2 fold change and RNAi ranged from -0.22 to -0.87 log2 fold change, comparable to EpiXcan effect sizes of 0.10 to 0.31 and -0.06 to -0.20 and PrediXcan effect sizes of 0.22 to 0.77 and -0.17 and -0.38 for corresponding eGenes.

**Figure 4.**
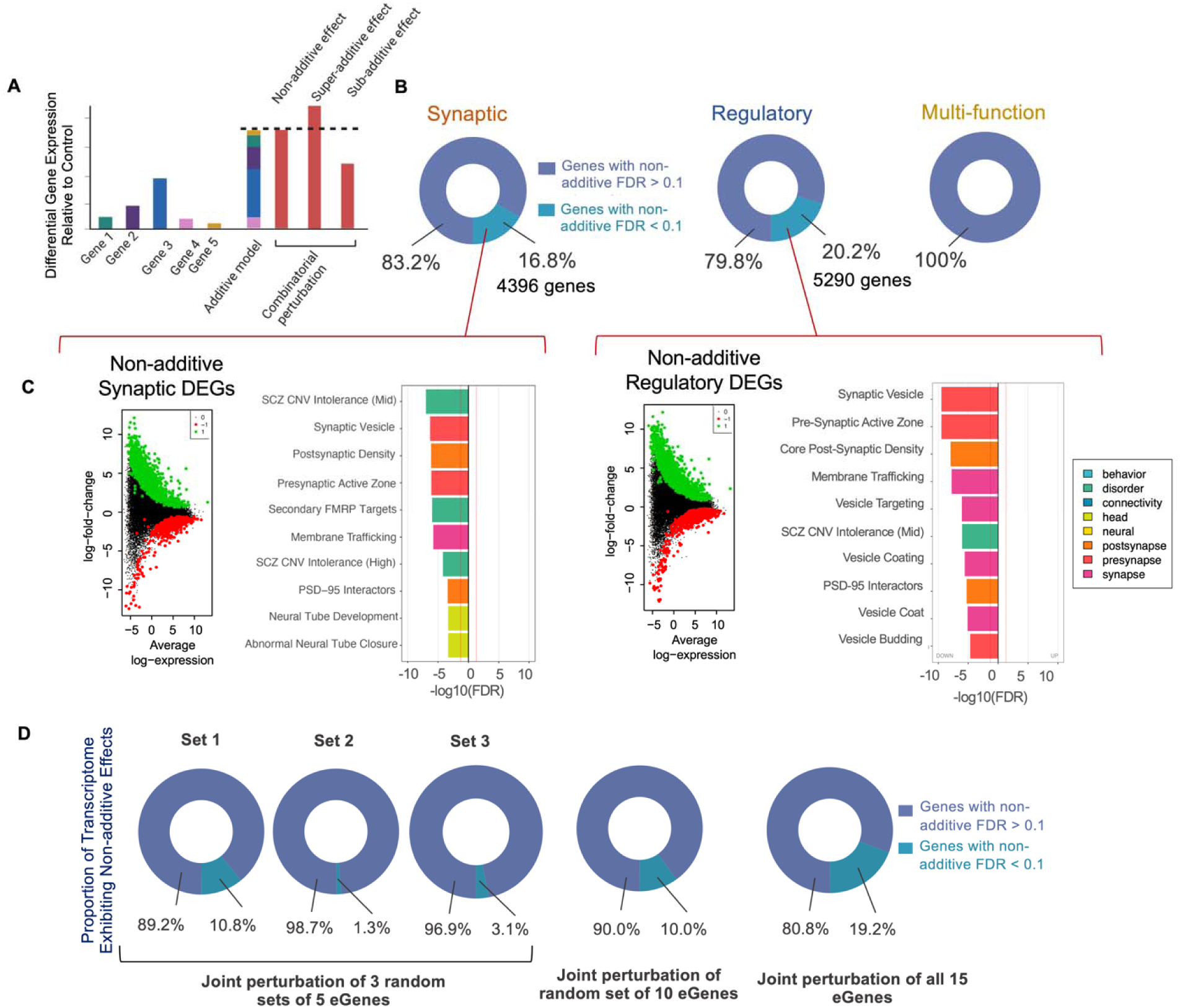
Perturbation of schizophrenia eGenes within functional categories results in non-additive effects on transcription impacting expression of genes linked to brain disorders and synaptic function. **A.** Schematic of differential expression analysis. Individual eGene perturbations, the implementation of the expected additive model based on the latter and the measured combinatorial perturbation permitting the detection of interactive effects through comparison with the additive model. **B.** Combinatorial perturbation of synaptic and regulatory eGenes resulted in non-additive effects on expression across 16.8% (synaptic) and 20.2% (regulatory) of the transcriptome. No significant non-additive effects were seen following joint perturbation of non-synaptic, non-transcriptional regulatory eGenes. **C.** GSEA of non-additive genes in the Synaptic eGene set demonstrated significant enrichment for genes relating to brain disorders and synaptic function. GSEA of non-additive genes in the Regulatory eGene set demonstrated significant enrichment for genes relating to brain disorders and synaptic function. SCZ = schizophrenia, CNV = copy number variant, FMRP = Fragile X Mental Retardation Protein, FDR = false discovery rate**. D.** Non-additive effects following combinatorial perturbation of sets of five, ten, and fifteen eGenes randomly assigned from the synaptic, regulatory, and multi-function eGene groups. The proportion of the transcriptome exhibiting significant non-additive effects increased with increasing numbers of perturbed eGenes (average of 5.1%, 10.0% and 19.2% of the transcriptome with non-additive FDR<0.1 after joint perturbations of five, ten, and fifteen eGenes respectively).

Across the majority of the schizophrenia eGenes in our arrayed screen, competitive gene-set enrichment analysis using 698 manually curated neural^60^ gene-sets (**SI Fig. 19B, C, SI Fig. 20A)** resulted in DEGs (p_FDR_<0.05) that were strongly enriched for gene-sets related to rare and common psychiatric disorder risk genes (11/15) (**SI Fig. 20B)**, pre-synaptic biology (10/15) (**SI Fig. 20C**), and glutamatergic neurotransmission (10/15) (**SI Fig. 20D**).

Differential expression following simultaneous perturbation of all five eGenes within each of the synaptic, regulatory, multi-function group (“measured combinatorial”), or random combinations thereof, was compared to the sum of differential expression for each single eGene perturbation (“expected additive”) (**Fig. 4; SI Fig. 21-23, Box 1; SI Table 4**). The difference between the measured combinatorial and expected additive model for each group represents the observed non-additive effects^83^ (**Fig. 4A**). Most genes were expressed as predicted by the expected additive model (i.e., did not differ significantly from the model), but combinatorial perturbations of synaptic regulatory, and multi-function eGenes resulted in significant non-additive effects (Bayes moderated t-statistics, FDR p < 0.1) across 16.8%, 20.2%, and 0% of the total transcriptome (π1 synergy coefficients^83^ of 43.86, 42.74, and 0.00, respectively) (**Fig. 4B**, non-additive p value distributions seen in **SI Fig 21E**). The log2FC of genes classified as “non-additive” was significantly larger than “additive” genes across all joint eGene perturbations (**SI Fig 23C**), and baseline expression of non-additive genes largely overlapped with those of additive genes (Log2(CPM) values within scramble control conditions, **SI Fig 23D**). Testing for overrepresentation in our curated neural gene sets showed that non-additive genes in the synaptic and regulatory joint eGene perturbations were significantly enriched for genes associated with risk for SZ as well as for pre- and postsynaptic gene sets (**Fig. 4C**). This was not explained by the magnitude of eGene perturbation between individual and combinatorial perturbations, which did not significantly differ (**SI Fig. 21B,** p>0.05 Wilcoxon ranked sum test). The degree of perturbation was strongly correlated with non-additivity for synaptic and regulatory eGenes, but not multifunctional eGenes – supporting that the type of eGene perturbed and not the magnitude of the perturbation alone drives non-additive effects (**SI Fig. 16A)**. Additionally, the strength of these correlations points to specific eGenes that may drive non-additive effects. For example, log2FC of *CLCN3* (synaptic) and *INO80E* (regulatory) are the most correlated with synergy coefficients (**SI Fig. 16C**). When evaluated across all eGene sets, the proportion of synaptic (Pearson’s r=0.49) and regulatory (r=0.45) eGenes in a set positively correlated with non-additivity, while proportion of multifunctional eGenes was strongly negatively correlated (r=-0.94). Additionally, higher semantic similarity scores based on molecular function (r=0.6), but not cellular component (r=0) or biological function (r=-0.05) positively correlated with non-additivity (**SI Fig. 16B**), overall indicating that the functional role of a given eGene strongly impacts interacting effects on transcription.

In addition to the impact of function on non-additivity, we also examined whether increasing the number of perturbed eGenes in a joint perturbation contributed to non-additive effects. Among randomised eGene sets, the size of the observed non-additive transcriptional effect increased directly with the total number of eGenes simultaneously perturbed, indicating that increasing the number of eGenes perturbed increases the degree of interactive effects on transcription (compare joint perturbations of random subsets of 5, 10 and 15 eGenes, **Fig. 4D, SI Fig 23A**).

Most (>95%) non-additive genes (whether up- or down-regulated, FDR p < 0.1) had “sub-additive” effects, defined as having less differential expression than predicted by the additive model (i.e., changes that were “less up” or “less down” than expected) (**SI Fig. 21C, Box 1**). Sub-additive genes were enriched for synaptic and brain disorder-associated gene-sets (**SI Data 1**).

Sub-additive effects do not reflect lack of combinatorial perturbation at the single cell level; observed non-additive effects were similar whether tested from a single multiplexed vector expressing all gRNAs^27^, or from independent expression vectors (**SI Fig. 22A-F**). Likewise, the strength and specificity to the non-additive effects resulting from pooled gRNA vectors were accurately recapitulated using a polycistronic gRNA vector (**SI Fig. 22G,H**), with 95% of non-additive effects validated. Notably, non-additive effects were more sensitively resolved using polycistronic vectors. Finally, modified ECCITE-seq confirmed a high number of unique gRNA integrations at the single cell level (**SI Fig. 22I)**.

To test the hypothesis that sub-additive effects arose from shared downstream signatures of individual eGene perturbations, we tested for convergent transcriptomic effects within each functional group (using METAL^78^, p< 1.92E-06). There was robust within-group convergence for the synaptic (1070 genes) and regulatory (1070 genes) eGene groups, but limited convergence across the multi-function eGenes (71 genes) (**Fig. 5B-E**). Most of the convergent genes overlapped with the non-additive genes (Fisher’s exact test, p<2.2E-16 for both synaptic and regulatory eGene groups), with 71% (761 of 1070) and 94% (1000 of 1070) of the convergent genes downstream of synaptic and regulatory eGenes, respectively, being including in their respective non-additive gene lists (**Fig. 5C,D**). Across all eight combinations of eGenes studied here (synaptic, regulatory, multi-function and random combinations of five, ten and fifteen eGenes), convergence across individual perturbations is correlated with the degree of non-additive effect seen in the corresponding joint perturbation condition (**Fig. 5A**, Pearson’s r^2^ = 0.6569, p=0.0147). Convergent effects of synaptic eGenes were enriched for synaptic function (e.g., mGluR5 interactors, p=1.64E-03) and brain disorders (e.g., schizophrenia GWAS, p=8.41E-05) gene-sets (**Fig. 5F**). Furthermore, convergent effects of regulatory eGenes were also enriched for brain disorder gene-sets (e.g., bipolar disorder, p=9.92E-06) (**Fig. 5G)**. Taken together, these findings highlight converging effects of schizophrenia eGenes on specific downstream genes relating to synaptic function and brain disorders.

**Figure 5.**
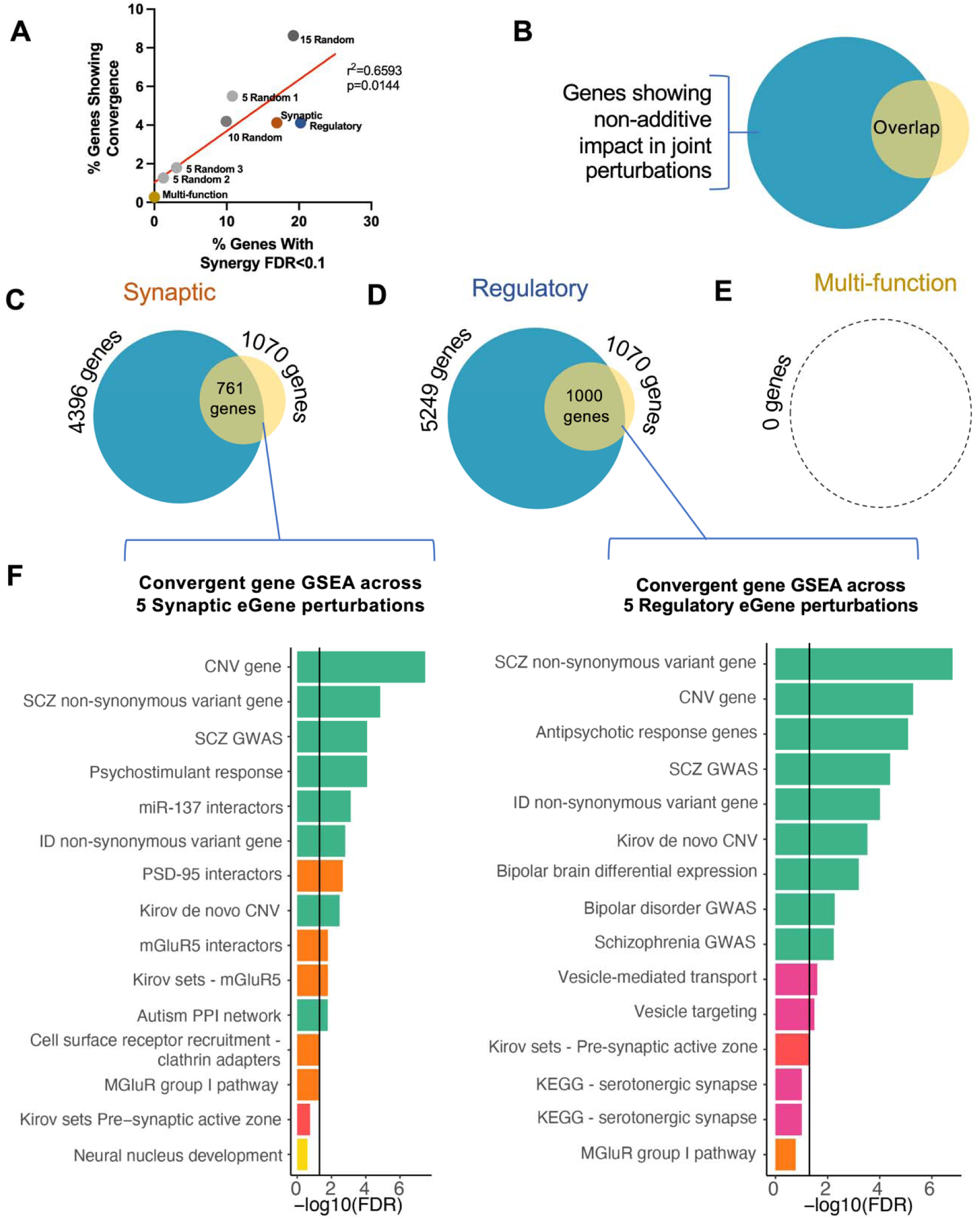
Convergence accounts for non-additive effects within functional pathways. A-E. Meta-analysis of differentially expressed genes (DEGs) elicited by individual eGene perturbations for each five-gene grouping using METAL to identify DEGs that showed altered expression consistently in the same direction across all five eGene perturbation conditions for each set of eGenes. **A.** Convergence across individual eGene perturbations is correlated with the degree of non-additive effect seen in the corresponding joint perturbation condition. Pearson’s r^2^ = 0.6569, p=0.0147. **B.** For each joint eGene perturbation group, non-additive impacts on transcription were compared with genes showing significant convergence across individual perturbations for the same eGene set. **C.** Evidence of convergence was found in 1070 genes across the synaptic eGene perturbations, 761 of which also exhibited non-additive effects in the additive-combinatorial comparison for the same set. **D.** Evidence of convergence was found in 1070 genes across the regulatory eGene perturbations, 1000 of which also exhibited non-additive effects in the additive-combinatorial comparison for the same set. **E.** No significant non-additive effects and only minimal convergence could be seen in eGene perturbations across functional pathways. **F.** GSEA of convergent genes in the synaptic and regulatory eGene groups demonstrated significant enrichment for genes relating to brain disorders and synaptic function.

Pathway-specific polygenic risk scores (PRS) (PRSet^84^) were calculated from non-additive signatures from synaptic (4,306 genes in PRS; R^2^=0.0431), regulatory (5,249 genes in PRS; R^2^=0.0419) and all fifteen eGenes (4,988 genes in PRS, R^2^=0.0425) (**SI Fig. 24A**). PRSet enrichment results explained the largest proportion of variance in schizophrenia case-control status, accounted for almost half of the total genome-wide PRS (19,340 genes plus SNPs in regions outside gene annotations in genome-wide PRS, R^2^= 0.0925) (**SI Fig. 24B**). Altogether, non-additive genes are enriched in GWAS signal in aggregation and harbor polygenic signal to the same degree as curated synaptic genes (**SI Fig. 24C-E**).

Bayesian network reconstruction^82^ from the arrayed experiment (n=63 samples, 4/sgRNA or shRNA, 25487 genes, and normalized to adjust for covariates such as donor) revealed a high-confidence co-expressed gene network that replicated across a maximum number of iterations (**Fig. 2B**). A densely interconnected network of 255 genes clustered together in at least 10% of the runs (**Fig. 2B**). This convergent network was significantly enriched for biological pathways implicated in schizophrenia etiology; over representation analysis revealed that target genes of schizophrenia, intellectual disability, and autism spectrum disorder common and rare variants were significantly over-represented in the network (**Fig. 2B,i**; **SI Data 3**), as well as genes regulated by miRNAs and transcription factors implicated in schizophrenia etiology, such as *hsa-miR-124a*^85^ and *NKX2*^86^ (**Fig. 2B,ii-iii)**. Separation of schizophrenia eGenes based on either signaling (**SI Fig. 9A**) or regulatory (**SI Fig. 9B**) function resolved unique convergent networks with no overlap in node genes, suggesting that the functional similarity of schizophrenia eGenes affects downstream convergence. Each of these networks included neuropsychiatric risk genes as well as those annotated for synaptic and immune signaling function (**SI Data 3**).

Although only nine node genes overlapped between convergent networks derived from arrayed and pooled experiments (**Fig. 2D**), both networks shared significant enrichments for targets of key miRNAs and transcription factors (**Fig. 2C**) – many with strong evidence of regulating gene expression associated with schizophrenia. The overlap in enrichments for regulatory factors suggests that while these unique groupings of eGene perturbations resolve largely disparate co-expression networks, they share an underlying regulatory structure.

### Prediction and empirical validation of drugs predicted to reverse convergent signatures in neurons

We identified drugs predicted to manipulate top node genes^87^. Across all eGene perturbations, reversers of convergent node signatures were enriched for mechanisms previously associated with psychiatric disorders, including HDAC inhibitors^88^ (normalized connectivity score (NCS)=-1.63; FDR adjusted pval<0.08), ATPase inhibitors^89^ (NCS=-1.61; FDR<0.08), and sodium channel blockers^90^ (NCS=-1.59; FDR<0.08). Mimickers of convergent node signatures were enriched for pathways associated with stress response, including glucocorticoid receptor agonists (NCS=1.66, FDR<0.08) and NFKB pathway inhibitors (NCS=1.60; FDR<0.2) (**SI Data 4**). Finding only nominally significant enrichments in non-neuronal cell lines suggests these may be neuron-specific drug responses.

Three drugs that opposed the transcription signatures of top convergent nodes specifically in neurons or neural progenitor cells (NPCs) were prioritized (see **Methods**, **Box 2**): anandamide (reverser of convergent network signature, NCS=-1.38, FDR=1 as well as *CALN1* signature alone, NCS=-1.23, FDR=0.16), simvastatin (NCS=-1.14, FDR=1; *TMEM219*, NCS=-0.8823, FDR=0.26), and etomoxir (NCS=-1.35, FDR=1; *CALN1,* NCS=-1.42, FDR=0.04; *TMEM219*, NCS=-1.09, FDR=0.01) (**SI Data 4**). These drugs were tested *in vitro* for their ability to reverse, or oppose, the effects of paired schizophrenia eGene perturbations. iGLUT CRISPRa for eGenes was followed by treatment with matched reverser drugs (*CALN1*: anandamide and etomoxir; *TMEM219*: simvastatin and etomoxir). Downstream transcriptomic and phenotypic assays were assessed to resolve eGene-drug effects on neuronal molecular, morphological, and physiological phenotypes (**Fig. 6**, **SI Fig. 28-30**). All drugs reversed or suppressed the transcriptomic impact of the CRISPRa perturbation alone. Notably, etomoxir not only limited the transcriptomic impact of perturbations of *CALN1* and *TMEM219*, but also impacted cell morphology (**Fig. 6C-D**), altering neurite length (adj. pval=0.0162) in a direction significantly opposed by *CALN1* perturbation (adj. pval=0.008) (**Fig. 6A-B**) and nominally opposed by *TMEM219* perturbation (adj. pval=0.082). Thus, it is possible to pharmacologically target convergent effects, even when schizophrenia eGenes themselves are not druggable.

**Figure 6.**
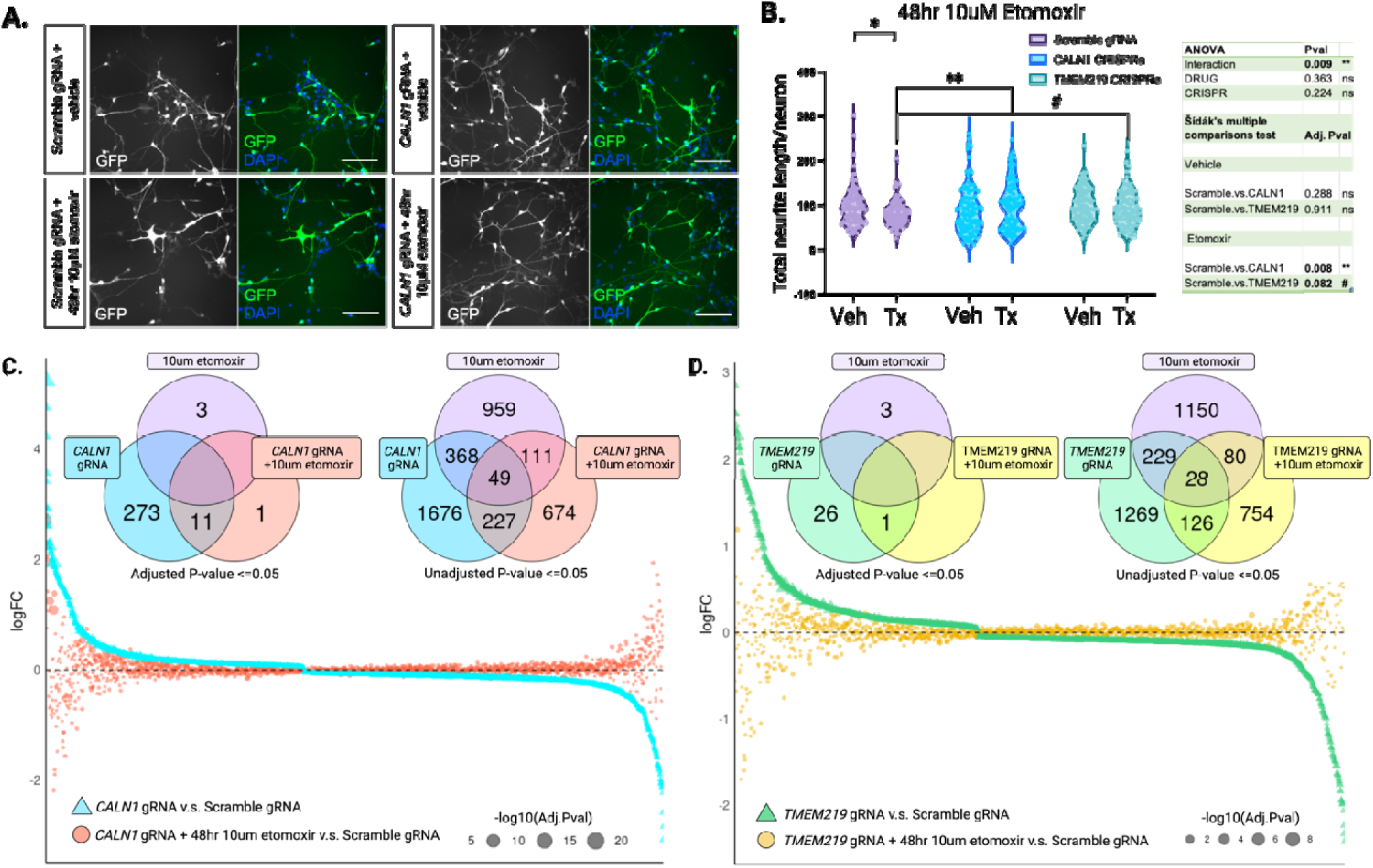
In vitro validation identifies opposing effects of in silico drug predictions and top schizophrenia eGenes. *In vitro* validation of drug-eGene phenotypic interactions. **A.** Imaging of Day 21 iGLUT neurons 48hrs after treatment with 10 µM etomoxir or vehicle after CRISPR activation of *CALN1* (compared to scramble gRNA control). Scale bar = 100 µm. **B.** 48-hour treatment with the small-molecule inhibitor of fatty acid oxidation (FAO) etomoxir significantly reduced neurite length in scramble control iGLUT neurons (Tukey’s multiple comparisons test; adj. pval=0.0162). Up-regulation of *CALN1* significantly rescued the effect of etomoxir, demonstrating opposing effect of the drug etomoxir and *CALN1* activation (adj. pval=0.008). Additionally, up-regulation of *TMEM219* demonstrated a nominal opposing effect to etomoxir (adj. pval=0.082) Volcano plots of total neurite length in CRISPRa Scramble/*CALN1/TMEM219* gRNA cells 48hrs after treatment with 10µM etomoxir or vehicle. ANOVA results table from a two-way Grouped ANOVA and Sidak’s multiple comparisons test of the interactive effects of CRISPRa perturbation and treatment with 48hr 10µM etomoxir. **C-D.** Treatment of cells perturbed with either *CALN1* CRISPRa **(C)** or *TMEM219* CRISPRa **(D)** with 10 µM etomoxir reverses or suppresses the transcriptomic impacts of the schizophrenia eGene perturbations alone (**SI Fig. 21-23**).

## DISCUSSION

Perturbations of schizophrenia eGenes resulted in convergent and non-additive transcriptional effects between functionally similar genes. First, a pooled CRISPRa study of twenty eGenes revealed a convergent effect concentrated on neurodevelopmental pathways, synaptic transmission, and voltage-gated ion channel activity. Second, across an independent but overlapping set of fifteen eGenes, within-function combinatorial perturbations resulted in sub-additive impacts on transcriptional profiles of hundreds of genes. The overlap of convergent genes and sub-additive genes suggests that redundant effects of eGenes with shared biological functions leads to saturation of directional impacts on gene expression. Overall, convergent signatures were robust, detected in three partially overlapping lists of schizophrenia eGenes, whether manipulated in arrayed or pooled experimental designs, and regardless of whether iGLUTs shared a common donor, cell type of origin, or developmental time point.

Would any large set of genetic perturbations uncover convergence and non-additivity? The issue of “negative control” is an important but challenging scientific question, particularly given the lack of power in some disease GWAS and the challenges in accurately mapping GWAS loci to gene targets highlighted by reviewers, but is one that must be addressed moving forward. Our results argue that future studies should expand beyond studying the effects of GWAS loci and their target eGenes in isolation, testing risk variant effects across multiple independent genetic backgrounds and in combination. We note that combinatorial eGene manipulations resulted in phenotypic changes in neuronal morphology, synaptic density, and neuronal activity that drastically differed from individual eGene perturbations (**SI Fig. 1F, SI Fig. 25-27**), reinforcing that polygenic risk cannot be extrapolated from experiments that test one risk gene at a time. These unexpected combinatorial effects could reflect compensatory homeostatic responses, or merely technical differences in the developmental timing of eGene perturbations and/or the inclusion of human astrocytes between the transcriptomic and phenotypic experimental paradigms. There is a need to investigate additivity, interactivity, and convergence across more neuronal and glia cell types and developmental stages.

Altogether, the experimental eGene perturbations approximated the magnitude and direction of predicted eGene effect associated with schizophrenia, and generally resulted in downstream gene expression changes related to synaptic biology and psychiatric disorder risk. Nonetheless, further gene set enrichment analysis using 493 inflammation and cell death gene-sets^47^ revealed enrichments related to cell stress and neurodegenerative diseases across many perturbations (**SI Data 1**). This enrichment was not seemingly associated with viral burden, being present whether single, combinatorial, or multiplexed vectors were applied. Indeed, if our *in vitro* system, defined by repeated lentiviral transduction, antibiotic selection, eGene perturbation, and single cell dissociation, stresses human neurons more than accounted for by the scramble gRNA controls, this would represent a concern of relevance to all applications of pooled CRISPR single screening experiments in human neurons. However, neither high content imaging nor multi-electrode array analyses indicated decreased cell survival or a cessation of neuronal activity (**SI Fig. 26, 27**). Moreover, inflammation^91^ and oxidative stress^92^, and particularly fetal exposures to inflammation, stress, and hypoxia^93,94^ are indeed associated with schizophrenia risk.

Mapping GWAS associations to eGene targets is challenging and can yield false positives. Our schizophrenia eGenes remain to be validated through newer methods (e.g., activity-by-contact model^95^) that do not rely on tissue-specific eQTL data. How well our three eQTL-based methods prioritized causal eGenes remains a critical question. In fact, there are frequent hotspots of multiple TWAS-associated genes in the same locus^72^, with co-regulation known to underlie pleiotropic TWAS associations^96^. We note that three eGenes prioritized in this study were linked to a single SNP (rs3814883) found within disease-associated copy number variant at 16p11.2, a locus that harbors the greatest excess of psychiatric common polygenic influences^97^. While it is possible that one or more eGenes at this locus were misidentified, an alternative possibility is that a causal GWAS SNP may co-regulate multiple adjacent and distal genes at this loci through chromatin contacts. For example, schizophrenia GWAS SNP rs2027349 altered expression of multiple genes: *VPS45*, IncRNA *AC244033*.2 and a distal gene, *C1orf54*; indeed, combinatorial perturbation of these eGenes results in non-additive impacts on transcriptomic and cellular phenotypes^28^.

Given the extent of polygenicity associated with schizophrenia, our conclusions are constrained by the small proportion of eGenes successfully manipulated relative to the total eGenes impacted by schizophrenia GWAS loci. Moreover, we selected for empirical evaluation only those schizophrenia eGenes with the very strongest evidence of genetically regulated gene expression. The generalizability of our observations to all schizophrenia eGenes is unclear, particularly if there are non-linear responses to gradual changes in gene dosage^98^. Thus, future investigation to test across larger gene sets, graded changes in expression^98^, *in vivo* brain regions^9^ and *in vitro* cell types^99^, developmental timespans^100^, drug/environmental contexts^101^ and donor backgrounds^102^ will inform the cell-type-specific and context-dependent nature of convergence and non-additivity. Of course, all of this must be considered within the caveat that *in vitro* CRISPR perturbations do not exactly recapitulate the possession of multiple genetic variants in human cases and controls. Despite this, it is worth noting that the limited number of perturbations used in our combinatorial conditions is still broadly relevant to studies of common variant interactions. When analyzing the full dataset of 105 S-PrediXcan SCZ eGenes in the post-mortem adult DLPFC^6^, a median of ten and a maximum of 37 eGenes had outlying expression in the direction of risk association per individual (**SI Fig. 17**). Across the twenty SCZ eGenes targeted in either the pooled or arrayed screens, a median of two and a maximum of eight eGenes had outlying expression in the direction of risk association per individual (**SI Fig. 18)**.

It is very challenging to extrapolate the extent to which our *in vitro* studies of CRISPR perturbations inform the physiological effects of polygenicity occurring in human cases and controls. Non-additive impacts remain difficult to assess using genomic approaches, despite the large number of methods that have been developed to improve detection^103^. Contrary to population-level additive models^104^, our findings suggest that at the individual-level some risk variants may have non-additive effects. We attempted to resolve this through eGene imputation in GWAS using two approaches. First, transcriptomic imputation of brain eGene expression (see **Methods, SI Fig. 1A**) revealed a dose-dependent effect: schizophrenia case-control status (p<0.01774, OR>1.10) was best predicted when three or more eGenes were perturbed (OR_3 eGenes_ = 1.47 vs. OR_1 eGene_ = 1.10). Second, transcriptomic risk scores were calculated (see **Methods**) to test whether the aggregate effect of eGenes was additive: schizophrenia risk was better predicted from larger (p<2.2 x 10^-16^) (**SI Fig. 1C**) or more biologically diverse (R=0.19, p<2.2×10^-16^) (**SI Fig. 1D**) gene groups. We speculate that population-level schizophrenia risk increases with the number of genes and pathways impacted (rather than within groups of functionally similar eGenes) due to the lack of individuals, either case or control, with strong imputed within-function perturbations. Pathway-specific polygenic risk scores^84^ that incorporate biological pathways, co-expression patterns, convergence, and/or non-additivity may improve patient stratification or better predict drug response.

How does our genetic analysis of convergent and combinatorial eGene effects inform precision medicine for patients with psychiatric disorders? How does our genetic analysis of convergent and combinatorial eGene effects inform precision medicine for patients with psychiatric disorders? First, convergence resolved across combinations of different SCZ eGenes may inform molecular subtypes of disease. For example, when we cluster individuals based on shared patterns of schizophrenia eGene up-regulation in the post-mortem DLPFC (**SI Data 5),** clusters were distinguished by the diagnosis of individuals within (Pearson’s Chi-squared; X^2^=140, df=21, p-value=9.51e-20) (e.g., cluster 8, SCZ, X^2^=3.286; cluster 12, affective disorders (AFF), X^2^=5.57; cluster 16, control, X^2^=3.014) (**SI Fig. 31-32**). A diagnosis of affective disorder (AFF) was significantly associated with up-regulation of *FES*, *NAGA*, *CALN1*, *CLCN3*, *SF3B1* and *ZNF804A* (cluster 12). Convergence analysis across these 6 eGenes in iGLUTs identifies the central node gene *ABCG2*—a biomarker associated with increased negative symptoms^105^, down-regulated in a neuroimmune molecular subtype (SCZ Type II)^106,107^, and associated with SCZ treatment resistance^108^. Second, we not only predicted drugs capable of reversing convergent transcriptomic signatures, but also demonstrated opposing action of drug treatment and schizophrenia eGene perturbations *in vitro*. We highlight statins, particularly simvastatin, which crosses the blood brain barrier and shows promise as an add-on treatment in schizophrenia^109^. Two double-blind placebo-controlled trials of simvastatin did not reduce overall symptom severity^110,111^ but highlighted the possibility that simvastatin may decrease negative symptoms in some patients. Secondary analysis highlights that baseline inflammatory profiles^112^ and treatment-induced changes in insulin receptor levels^113^ may serve as biomarkers capable of predicting simvastatin response.

That convergent genes were enriched for risk factors and transcriptomic signatures associated with a range of other brain disorders indicated that convergent effects may in part explain shared features of psychiatric disorders and cross-disorder pleiotropy of risk. Consistent with this, common and rare risk variants for schizophrenia^2,17,38–42,114–116^, autism spectrum disorder^117–119^ and more broadly across the neuropsychiatric disorder spectrum^30,120–122^ are all highly enriched for genes involved in synaptic biology and gene regulation, further highlighting the need to explore the impact of functional similarity in the interactions between rare and common variants. Our findings support the hypothesis that common and rare psychiatric risk variants converge on the same biological pathways^27^. There is great value in continuing to explore as many rare and common risk loci as possible, agnostic to previously defined function, in a cell-type-specific and context-dependent manner. The as-yet unidentified regulatory nodes that lead to the disruptions in synaptic function believed to underlie psychiatric endophenotypes represent exciting and unexplored therapeutic targets. Our overarching goal is to advance the field towards an era of precision medicine^123^, whereby not just each patient’s genetic variants, but also the expected interactions between them, can be used to predict symptom development, disorder trajectory, and potential therapeutic interventions.

## MATERIALS AND METHODS

### Schizophrenia eGene Prioritization

“eGenes: are defined as genes with significant genetic regulation of gene expression levels. In total, across the pooled and arrayed analyses, 20 unique eGenes were prioritized based on statistical and epigenetic evidence supporting genetic (dys)regulation of expression in schizophrenia (see, **Table 1**), rather than GWAS or eQTL effect size; predicted direction and magnitude of eGene effect available in **SI Table 1**.

i) “**SCZ1** eGenes”: EpiXcan^69^ was used to impute brain transcriptomes from Psychiatric Genomics Consortium 3 (PGC3)-SCZ GWAS^2^ at the level of genes and isoforms from the PsychENCODE post-mortem datasets of genotyped individuals (brain homogenate, n=924)^43,70^; EpiXcan increases power to identify trait-associated genes under a causality model by integrating epigenetic annotation^124^ (from REMC^125^); transcriptomes were imputed at the gene and isoform levels and features with training cross-validation R^2^≥0.01 were retained. The epigenetic imputation models were built with the PrediXcan^73^ method (using a 50kbp window instead of 1Mbp for transcripts) utilizing the recently described ChIPseq datasets^21^; summary-level imputation was performed with S-PrediXcan^71^. Peaks were assigned to genes with the ChIPseeker R package^126^. In addition, PrediXcan^73^ imputed H3K27ac (brain homogenate, n=122; neuronal, n=191) and H3K4me3 (neuronal, n=163)^21^ to more confidently identify *cis* regulatory elements associated with risk for SCZ. Overall, SCZ eGenes were prioritized from GWAS based on: i) significant genetic up-regulation of expression (z-score >6 for genes), ii) epigenetic support (imputed epigenetic activity (p<0.01) across at least one of the three assays), iii) exclusion of non-coding genes or those located in the MHC locus, iv) robust expression in our hiPSC neuron RNAseq. Genes were ranked based on the association z-score for imputed gene expression. For pooled experiments (day 7 hiPSC-derived iGLUT), six top coding genes and one top pseudo-gene were selected: *NEK4, PLCL1, UBE2Q2L, NAGA, FES, CALN1,* and *ZNF804* (**Table 1A**).

ii) “**SCZ2** eGenes”: First, transcriptomic imputation (prediXcan^71–73^) identified ∼250 significant genes (p<6×10^-6^) with predicted differential expression between SCZ-cases and controls using SCZ GWAS^2^ and post-mortem CommonMind Consortium (CMC)^6^ data (623 samples). Second, colocalization (COLOC^74,75^) of fine-mapped PGC3-GWAS^2^ loci (65,205 cases and 87,919 controls) with post-mortem brain^6^ eQTL (537 EUR samples)^6^ identified 25 loci with very strong evidence (high posterior probability that a single shared variant is responsible for both signals, PP4 > 0.8^74^). There was significant overlap between the two analyses (binomial test p-value 3.03×10^-112^); of the 25 COLOC genes, 22 were also significant by PrediXcan. For each eGene, the magnitude and direction of perturbation associated with SCZ risk was predicted, and expression confirmed in hiPSC neuron RNAseq^27^. eGenes were further separated into discrete functional categories based on gene ontology annotations (http://geneontology.org/). From these 22, we prioritized the top coding genes across three broad categories: synaptic, regulatory, and multifunction (defined as not synaptic, regulatory, and seemingly unrelated to each other). To complete selection of five genes from each category, three additional top-ranked synaptic genes from the prediXcan analysis were included: *DOC2A*^75^, *CLCN3*^75^ and *PLCL1*^122^. Overall, 15 SCZ eGenes were prioritized from GWAS based on i) significant genetic regulation by COLOC and/or PrediXcan, ii) exclusion of non-coding genes and those located in the major histocompatibility complex (MHC) locus, iii) robust expression in our hiPSC neuron RNAseq.

For arrayed experiments (day 21 NPC-derived iGLUT), our final gene list for combinatorial perturbations included five synaptic genes (*SNAP91, CLCN3, PLCL1, DOC2A, SNCA*), five regulatory genes (*ZNF823, INO80E, SF3B1, THOC7, GATAD2A*), and five genes with non-synaptic, non-regulatory functions, termed “multi-function” (*CALN1, CUL9, TMEM219, PCCB, FURIN*) (**Table 1B**). For pooled experiments (day 21 NPC-derived iGLUT), the ten coding genes with significant genetic up-regulation were selected: *CALN1*, *CLCN3, CUL9, DOC2A, PLCL1, INO8E0, SF3B1, SNAP91, TMEM219, ZNF823*. This list was combined with our eGene set previously evaluated in hiPSC-neurons ^27^; one functionally validated gRNA was included for each of these three genes (*SNAP91*, *TSNARE1*, and *CLCN3)*^27^.

### gRNA design

CRISPRa gRNA design and cloning were conducted as described previously^127^. For the fifteen eGenes prioritized by a combination of COLOC and PrediXcan, we designed three gRNAs each. For the seven eGenes prioritized by EpiXcan and PrediXcan, we designed ten gRNAs each. For the three previously tested eGenes^27^ (intended a positive control), we used one pre-validated gRNA each.

### iGLUT induction from hiPSC-derived NPCs^27,59–61^ or hiPSCs^66,128^

Validated control hiPSCs for eGene perturbation were selected from a previously reported case/control hiPSC cohort of childhood onset schizophrenia^129^. Informed consent was obtained from all fibroblast donors at the National Institute of Mental Health, under the review of the Internal Review Board of the NIMH. All hiPSC work was reviewed by the Internal Review Board of the Icahn School of Medicine at Mount Sinai. This work was also reviewed by the Embryonic Stem Cell Research Oversight Committee at the Icahn School of Medicine at Mount Sinai and Yale University. The following control hiPSC/NPCs were used: NSB553-S1-1 (male), NSB2607-2/NSB2607-1-4 (male), NSB690-2 (male). All fibroblast samples were genotyped by IlluminaOmni 2.5 bead chip genotyping^130,131^, PsychChip^129^, and exome sequencing^129^. Parental hiPSCs were validated by G-banded karyotyping (Wicell Cytogenetics), with ongoing genome stability monitored by Infinium Global Screening Array v3.0 (lllumina). Critically, SNP genotype is inferred from all RNAseq data using the Sequenom SURESelect Clinical Research Exome (CRE) and Sure Select V5 SNP lists to confirm that neuron identity matches donor.

i) Validated control hiPSC-derived NPCs for CRISPRa/shRNA were selected from a previously reported case/control hiPSC cohort of childhood onset SCZ (COS)^129^: NSB553-S1-1 (male, average SCZ PRS, European ancestry), NSB2607-1-4 (male, average SCZ PRS, European ancestry). hiPSC-NPCs expressing dCas9-VPR were generated and validated as previously described)^129^ and cultured in hNPC media (DMEM/F12 (Life Technologies #10565), 1x N2 (Life Technologies #17502-048), 1x B27-RA (Life Technologies #12587-010), 1x Antibiotic-Antimycotic, 20 ng/ml FGF2 (Life Technologies)) on Matrigel (Corning, #354230). hiPSC-NPCs at full confluence (1-1.5×10^7^ cells / well of a 6-well plate) were dissociated with Accutase (Innovative Cell Technologies) for 5 mins, spun down (5 mins X 1000g), resuspended, and seeded onto Matrigel-coated plates at 3-5×10^6^ cells / well. Media was replaced every two-to-three days for up to seven days until the next split.

At day -2, dCas9-VPR hiPSC-NPCs were seeded as 1.2×10^^6^ cells / well in a 12-well plate coated with Matrigel. At day -1, cells were transduced with rtTA (Addgene 20342) and *NGN2* (Addgene 99378) lentiviruses. Medium was switched to non-viral medium four hours post infection. At day 0 (D0), 1 µg/ml dox was added to induce *NGN2*-expression. At D1, transduced hiPSC-NPCs were treated with antibiotics to select for lentiviral integration (300 ng/ml puromycin for dCas9-effectors-Puro, 1 mg/ml G-418 for NGN2-Neo). At D3, NPC medium was switched to neuronal medium (Brainphys (Stemcell Technologies, #05790), 1x N2 (Life Technologies #17502-048), 1x B27-RA (Life Technologies #12587-010), 1 µg/ml Natural Mouse Laminin (Life Technologies), 20 ng/ml BDNF (Peprotech #450-02), 20 ng/ml GDNF (Peprotech #450-10), 500 µg/ml Dibutyryl cyclic-AMP (Sigma #D0627), 200 nM L-ascorbic acid (Sigma #A0278)) including 1 µg/ml Dox. 50% of the medium was replaced with fresh neuronal medium once every second day.

For pooled analysis, on day 5, young hiPSC-NPC *NGN2*-neurons were replated onto matrigel-coated plates and cells were dissociated with Accutase (Innovative Cell Technologies) for 5-10 min, washed with DMEM/10%FBS, gently resuspended, counted and centrifuged at 1,000·g for 5 min. The pellet was resuspended at a concentration of 1·10^6^ cells/mL in neuron media [Brainphys (StemCell Technologies #05790), 1·N2 (ThermoFisher #17502-048), 1·B27-RA (ThermoFisher #12587-010), 1 mg/ml Natural Mouse Laminin (ThermoFisher #23017015), 20 ng/mL BDNF (Peprotech #450-02), 20 ng/mL GDNF (Peptrotech #450-10), 500 mg/mL Dibutyryl cyclic-AMP (Sigma #D0627), 200 nM L-ascorbic acid (Sigma #A0278)] with doxycycline, puromycin, G418 [4µM Ara-C (Sigma #C6645)] and 1·Thiazovivin (Sigma #420220). Cells were seeded 5·10^5^ per 12-well plate. For arrayed analysis, neurons were not replated, owing to the complexity of conditions involved.

At D13, iGLUTs were treated with 200 nM Ara-C to reduce the proliferation of non-neuronal cells in the culture, followed by half medium changes. At D18, Ara-C was completely withdrawn by full medium change while adding media containing individual shRNA/gRNA vectors or pools of mixed shRNA and gRNA vectors (Addgene 99374), either targeting eGenes or scramble controls. Medium was switched to non-viral medium four hours post infection. At D19, transduced iGLUTs were treated with corresponding antibiotics to the gRNA lentiviruses (1 mg/ml HygroB for lentiguide-Hygro/lentiguide-Hygro-mTagBFP2) followed by half medium changes until neurons were harvested at D21.

ii) Clonal hiPSCs from two control donors of European ancestry (NSB690-2 (male, average SCZ PRS, European ancestry) and NSB2607-2 (male, average SCZ PRS, European ancestry)^129^ with lenti-EF1a-dCas9-VPR-Puro (Addgene #99373), pLV-TetO-hNGN2-eGFP-Neo (Addgene #99378), and lentiviral FUW-M2rtTA (Addgene #20342) were maintained in StemFlex™ Medium (ThermoFisher #A3349401) and passaged with EDTA (Life Technologies #15575-020). On day 1, induction media (DMEM/F12 (ThermoFisher #10565,), 1· N2 (ThermoFisher #17502-048), 1· B27-RA (ThermoFisher #12587-010), 1· Antibiotic-Antimycotic (ThermoFisher #15240096), and 1 µg/mL doxycycline) was prepared and dispensed 2 mL of suspension at 1.2·10^6^ cells/well in induction media onto a 6-well plate coated with matrigel (Corning #354230). On day 3, media is replaced with induction medium containing 1 μg/mL puromycin and 1 mg/mLG418. On day 5, split neurons were replated onto matrigel-coated plates and cells were dissociate with Accutase (Innovative Cell Technologies) for 5-10 min, washed with DMEM/10%FBS, gently resuspended, counted and centrifuged at 1,000·g for 5 min. The pellet was resuspended at a concentration of 1·10^6^ cells/mL in neuron media [Brainphys (StemCell Technologies #05790), 1·N2 (ThermoFisher #17502-048), 1·B27-RA (ThermoFisher #12587-010), 1 mg/ml Natural Mouse Laminin (ThermoFisher #23017015), 20 ng/mL BDNF (Peprotech #450-02), 20 ng/mL GDNF (Peptrotech #450-10), 500 mg/mL Dibutyryl cyclic-AMP (Sigma #D0627), 200 nM L-ascorbic acid (Sigma #A0278)] with doxycycline, puromycin, G418 [4µM Ara-C (Sigma #C6645)] and 1·Thiazovivin (Sigma #420220). Cells were seeded 5·10^5^ per 12-well plate. On day 7, neurons were harvested for scRNA sequencing.

### Neuronal Pooled CRISPRa screens

<u>E</u>xpanded <u>C</u>RISPR-compatible <u>CITE-seq</u> (ECCITE-seq)^76^, combines Cellular Indexing of Transcriptomes and Epitopes by sequencing (CITE-seq) and Cell Hashing for multiplexing and doublet detection^132^ with direct detection of sgRNAs to enable single cell CRISPR screens with multi-modal single cell readout. By capturing pol III-expressed guide RNAs directly, this approach overcomes limitations of other single-cell CRISPR methods, which detect guide sequences by a proxy transcript, resulting in barcode switching and lower capture rates^133–135^. CRISPRa hiPSC iGLUT neurons (2607 (male) and 690 (male)) were transduced with the pooled gRNA at day -1. After maturation, 7-day-old iGLUT neurons were dissociated to single cell suspensions with papain, antibody-hashed^132^, and bar-coded single cell cDNA generated using 10X Genomics Chromium^136^. NPC-derived iGLUT neurons (2607 (male) and 553 (male)) were transduced with the mixed-pooled gRNA vectors (Addgene 99374) at day 17. At day 21, media was replaced by 0.5ml/well accutase containing 10 μm Rock inhibitor, THX (catalog no. 420220; Millipore) for 1 hour to dissociate neurons. Neurons were spun down (3 mins X 300g) and resuspended in DMEM/F12 + THX before proceeding to single cell sequencing.

### Analysis of single-cell CRISPRa screens in DIV 7 and DIV 21 iGLUT Neurons

mRNA sequencing reads were mapped to the GRCh38 reference genome using the *Cellranger* Software. To generate count matrices for HTO and GDO libraries, the kallisto indexing and tag extraction (kite) workflow were used. Count matrices were used as input into the R/*Seurat* package^137^ to perform downstream analyses, including QC, normalization, cell clustering, HTO/GDO demultiplexing, and DEG analysis^76,138^.

Normalization and downstream analysis of RNA data were performed using the Seurat R package (v.2.3.0), which enables the integrated processing of multimodal single-cell datasets. Each ECCITE-seq experiment was initially processed separately. Cells with RNA UMI feature counts were filtered (200 < nFeature_RNA < 8000) and the percentage of all the counts belonging to the mitochondrial, ribosomal, and hemoglobin genes calculated using Seurat::PercentageFeatureSet. Hashtag and guide-tag raw counts were normalized using centered log ratio transformation, where counts were divided by the geometric mean of the corresponding tag across cells and log-transformed. For demultiplexing based on hashtag, Seurat::HTODemux function was used; and for guide-tag counts Seurat::MULTIseqDemux function within the Seurat package was performed with additional MULTIseq semi-supervised negative-cell reclassification. In both experiments, 8-10% of retained cells contained multiple gRNAs and were assigned as doublets after de-multiplexing. To remove variation related to cell-cycle phase of individual cells, cell cycle scores were assigned using Seurat::CellCycleScoring which uses a list of cell cycle markers^139^ to segregate by markers of G2/M phase and markers of S phase. RNA UMI count data was then normalized, log-transformed and the percent mitochondrial, hemoglobulin, and ribosomal genes, batch, donor (HTO-maxID; as a biological replicate), cell cycle scores (Phase) regressed out using Seurat::SCTransform. The scaled residuals of this model represent a ‘corrected’ expression matrix, that was used for all downstream analyses. To ensure that cells assigned to a guide-tag identity class demonstrated successful perturbation of the target gene, we performed ‘weighted-nearest neighbor’ (WNN) analysis, to assign clusters based on both guide-tag identity class and gene expression^77^. To identify successfully perturbed cells, we calculated a p-value based on the Wilcox rank sum test and Area Under the Curve (AUC) statistic, which reflects the power of each gene (or gRNA) to serve as a marker of a given cluster using Presto. WNN Clusters were then filtered based on two criteria (1) single gRNA-identity with an AUC statistic of >= 0.8 (where 1 means the gRNA is a perfect marker of a given cluster) and (2) a logFC >= 2 standard deviations of the mean or logFC > 0 and p-val > 0.05, of the target gene (but no other target genes) compared to scramble (non-targeting sgRNAs) controls (**SI Fig. 4-8**). These clusters were then used for downstream analyses^140^.

### Cell Fraction Imputation and Quantification of Heterogeneity in Composition of iGLUT neurons

Using CiberSortx, we imputed the cell-faction identity of randomly sampled scramble control cells from each experiment (n=100/exp) using the PsychEncode scRNAseq dataset as a reference (100 permutations). To determine if the level of heterogeneity of iGLUT neuron maturity and subtype was similar between DIV7 and DIV21 iGLUT neurons in the given experiments, we performed a non-parametric Levene’s Test for Homogeneity of Variance (LT-test) on the imputed cell fraction matrices. Although we observed heterogeneity in relative central and peripheral nervous system marker expression across the cell fractions, this heterogeneity was not due to gRNA identity and the level of variance in our data due to cellular heterogeneity was not significantly different by time-point. We were underpowered to compare gRNAs between cells with higher expression of different cell markers.

### Meta-analysis of gene expression across perturbations

We performed a meta-analysis and Cochran’s heterogeneity Q-test (METAL^78^) using the p-values and direction of effects (t-statistic), weighted according to sample size across all sets of perturbations in both the arrayed and pooled assays (Target vs. Scramble DEGs). Genes were defined as “convergent” if they (1) had the same direction of effect across all 5, 10, or 15 target combinations, (2) were Bonferroni significant in our meta-analysis (Bonferroni adjusted p-value <= 0.05), and (3) had a heterogeneity p-value = >0.05.

### Bayesian Bi-clustering to identify Target-Convergent Networks

eGene-Convergent gene co-expression Networks (eGCN)^82^ were built using an unsupervised Bayesian biclustering model, BicMix^141^, on the log2CPM expression data from all the replicates across each of the 5-target sets and scramble gRNA jointly or all the cells across 10 targets and scramble gRNA jointly. for the arrayed and pooled assays respectively. To account for neuronal maturity differences in the single-cell screen, expression matrices were batch corrected and normalized and the scramble cells from both experiments (matched scramble gRNA across experiments) used as a single control population. To perform this as a joint analysis across two experiments, (1) Count matrices from each experiment were combined and RNA transcripts, mitochondrial, ribosomal, and hemoglobin genes were removed ([‘^MT-|^RP[SL][[:digit:]]|^RPLP[[:digit:]]|^RPSA|^HB[ABDEGMQZ][[:digit:]]’) as well as genes that had at fewer than 2 read counts in 90% of samples, (2) and limma:voom normalization and transformation was used to compute the log2cpm counts from the effective library sizes of each cell (16851 genes). 40 runs of BicMix were performed on these data and the output from iteration 400 of the variational Expectation-Maximization algorithm was used. The hyperparameters for BicMix were set based on previous extensive simulation studies^142^. Convergent networks were identified across all possible combinations of 2-14 as well as all 15 of the targets (n=32752 combinations) in the arrayed assay, and all possible combinations of 2,3,4,5,6,7 or 8 as well as all 10 of the targets (n=1003 combinations) in the pooled screen. Network connections that did not replicate in more than 10% of the runs were excluded. Nodes with less than 5 edges or non-coding genes were removed from gene set enrichment analysis (GSEA). (The threshold of >5 edges is based on the likelihood of more than 5 edges being present by chance, with 10% being the percentage of runs where the connection was identified, see^82,141^. Duplication thresholds are network-dependent and a metric of confidence in the connections; including those with especially low duplication rates were not included in downstream analysis.) Of all random sets tested in the pooled screen, 64.8% resolved a convergent network passing at least a 10% duplication threshold; of all random sets tested in the arrayed experiment, ∼50% resolved a convergent network with a 5-255 threshold of duplication depending on the node-edge connection. Using FUMAGWAS: GENE2FUNC, the protein-coding genes were functionally annotated and overrepresentation gene-set analysis for each network gene set was performed^143^. Using WebGestalt (WEB-based Gene SeT AnaLysis Toolkit)^144^, over-representation analysis (ORA) was performed on all convergent network gene sets against a curated list of common and rare variant target genes across ASD, BIP, SCZ, and ID^27^.

### Influence of Functional Similarity on Convergence Degree

Functionally similarity scores across the eGenes represented in each set was calculated using three metrics: (1) Gene Ontology Scores: the average semantic similarity score based on Gene Ontology pathway membership (within Biological Pathway (BP), Cellular Component (CC), and Molecular Function (MF) between genes in a set^81^, (2) Brain expression correlation (B.E.C.) score: based on the strength of the correlation in gene expression in the CMC (n=991 after QC) post-mortem dorsa-lateral prefrontal cortex (DLPFC) gene expression data^6^, and (3) Signaling Score: based on the proportion of eGenes whose basic functional annotation was categorized as “signaling” (*CALN1, CLCN3, FES, NAGA, PLCL1, TMEM219*; with *PLCL1* and *CLCN3* further separated as specific synaptic genes) or four “epigenetic/regulatory” target genes (*SF3B1, UBE2Q2L, ZNF823, ZNF804A*; with *ZNF823, ZNF804A* as specific transcription factors) using FUMAGWAS: GENE2FUNC^143^ (**SI Fig. 10**).

Bi-clustering identifies co-expressed genes shared across the downstream transcriptomic impacts of any given set of eGene perturbations, thus, the resolved networks are the transcriptomic similarities between distinct perturbations (convergence). While bi-clustering resolves convergent gene co-expression networks, the strength of convergence within a network can vary. We assigned each network a “degree of convergence” based on (1) network connectivity and (2) similarity of network genes based on biological pathway membership. We performed a principal components analysis on the functional similarity scores and the degree of network convergence. PCA loadings determined the effect of the included variables on the variability across all resolvable sets (arrayed=16320, pooled=827, variables=6). To quantify this, we calculated two small world connectivity network coefficients: the cluster connectivity coefficient based on the proportion of edges present out of all possible edges (*Cp*) and the average path length (*Lp*)^145^.

Here we define convergence as (1) increased connectivity of the resolved networks, and (2) functional similarity of genes within the network. Network connectivity was defined by the sum of the clustering coefficient (Cp) and the difference in average length path (Lp) from the maximum average length path resolved across all possible sets [(max)Lp-Lp]. Network functional similarity was scored by taking the sum of the mean semantic similarity scores between all genes in the network. Overall, convergence degree represented the sum of the network connectivity score and the network functional similarity score.

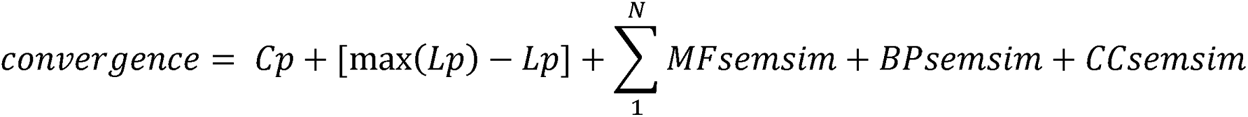

### Convergent networks with matched patterns of gene expression in the post-mortem brain

We clarify that this approach asks how often eGenes are up-regulated together in individual post-mortem brains. To do this, we ran target-convergent network reconstruction in our scRNA-seq data, not the CMC bulk tissue data, for sets of eGenes defined by the clustering observed in the CMC bulk tissue data. We found zero individuals in the CMC data with significant upregulation of all ten risk eGenes. Instead of only evaluating convergence on the basis on eGene functional similarity as in the first portion of the manuscript, we define eGene pairings more broadly based on the signatures of these eGenes in the post-mortem DLPFC – increasing the relevancy to risk at the individual level. Target sets based on gene expression patterns in the CMC (n=991 after QC) post-mortem dorsa-lateral pre-frontal cortex (DLPFC). We performed K-means clustering to subset the data into clusters based on the Z-scored gene expression of the 10 target genes. Although initial silhouette analysis identified the optimal number of clusters as two, visualization by a scree plot suggested the optimal number to be between 4-6 clusters. Given that data clustered by case/control status (2 clusters), and sub-diagnosis of BP, SCZ, AFF, and Controls (4 clusters), to assess clustering based on 10 eGenes, we tested the impact of using 10 clusters and 20 clusters (**SI Fig. 32**). Perturbation identities were assigned based on average positive Z-scores of >=0.5 within each cluster. We then assigned our single-cell data to clusters based on the overlap of perturbations and performed network reconstruction to replicate our convergent analysis using groups based on CMC post-mortem data. We retained clusters that resolved networks with at least 10% duplication rate and calculated convergence scores and performed GSEA using protein-coding network genes. Of the twenty clusters, networks were recovered for the combination of targets represented in cluster 4 (2 targets; 913 cells; 15% duplication; 13 node genes), cluster 5 (3 targets; 1260 cells; 15% duplication; 13 node genes), cluster 6 (6 targets; 2035 cells; 15% duplication; 34 node genes), cluster 9 (6 targets, 1822 cells, 20% duplication, 108 node genes), cluster 11 (5 targets; 1640 cells; 15% duplication; 25 node genes), cluster 12 (6 targets; 2357 cells; 20% duplication, 152 node genes), cluster 13 (5 targets; 1741 cells; 17.5% duplication, 17 node genes), cluster 18 (6 targets; 1884 cells, 15% duplication, 25 nodes), cluster 19 (6 targets, 2327 cells, 20% duplication, 153 nodes), cluster 20 (6 targets, 2015 cells, 20% duplication, 33 nodes), while low confidence convergence was resolved for cluster 1 (5 targets, 1600 cells; 7.5% duplication; 38 node genes), cluster 8 (3 targets, 1233 cells, 7.5% duplication, 38 node genes), cluster 14 (3 targets, 1020 cells, 5% duplication, 23 nodes) and 16 (4 targets, 1177 cells, 2.5% duplication, 16 nodes). To determine if convergent networks were distinct between diagnostic groups, we first performed a Pearson’s chi-squared test to determine whether there was a significant difference between the expected frequencies and the observed frequencies in diagnosis of AFF, BIP and SCZ within the clusters and then calculated Jaccard Similarity Indices between clusters based on convergent network gene membership.

### Drug prioritization based on perturbation signature reversal in LiNCs Neuronal Cell Lines

To identify drugs that could reverse the resolved convergent perturbation signature across all ten targets, and within each target individually, we used the Query tool from The Broad Institute’s Connectivity Map (Cmap) Server. Briefly, the tool computes weighted enrichment scores (WTCS) between the query set and each signature in the Cmap LINCs gene expression data (dose, time, drug, cell-line), normalizes the WRCS by dividing by signed mean w/in each perturbation (NCS), and computes FDR as fraction of “null signatures” (DMSO) where the absolute NCS exceeds reference signature^146^. We prioritized drugs that reversed signatures specifically in neuronal cells (either neurons (NEU) of neural progenitor cells (NPCs) with NCS <= -1.00) and filtered for (i) drugs that cross the blood-brain barriers, (ii) drugs that target genes expressed in iGLUT neurons based on bulk RNA-sequencing data from our lab and (ii) drugs that are currently launched or in clinical trial according the cMAP Drug Repurposing database and without evidence of neurotoxicity (**Box 2**).

### CRISPRa/shRNA Validation^27^

At day -2, dCas9-VPR hiPSC-NPCs were seeded as 0.6×10^^6^ cells / well in a 24-well plate coated with Matrigel. At day -1, cells were transduced with rtTA (Addgene 20342) and *NGN2* (Addgene 99378) lentiviruses. Medium was switched to non-viral medium four hours post infection. At D0, 1 µg/ml dox was added to induce *NGN2*-expression. At D1, transduced hiPSC-NPCs were treated with corresponding antibiotics to the lentiviruses (1 mg/ml G-418 for *NGN2*-Neo) in order to increase the purity of transduced hiPSC-NPCs. At D3, NPC medium was switched to neuronal medium (Brainphys (Stemcell Technologies, #05790), 1x N2 (Life Technologies #17502-048), 1x B27-RA (Life Technologies #12587-010), 1 µg/ml Natural Mouse Laminin (Life Technologies), 20 ng/ml BDNF (Peprotech #450-02), 20 ng/ml GDNF (Peptrotech #450-10), 500 µg/ml Dibutyryl cyclic-AMP (Sigma #D0627), 200 nM L-ascorbic acid (Sigma #A0278)) including 1 µg/ml Dox. 50% of the medium was replaced with fresh neuronal medium once every second day. At D4 individual shRNA/gRNA vectors (Addgene 99374), either targeting eGenes or scramble controls. 3-5 vectors were tested per eGene. Medium was switched to non-viral medium four hours post infection. At D5, transduced iGLUTs were treated with corresponding antibiotics to the gRNA lentiviruses (1 mg/ml HygroB for lentiguide-Hygro/lentiguide-Hygro-mTagBFP2) before harvesting at D7 in order to assess eGene perturbation efficacy via qPCR.

### Real time-quantitative PCR

Real time qPCR was performed as previously described^127^. Specifically, cell cultures were harvested with Trizol and total RNA extraction was carried out following the manufacturer’s instructions. Quantitative transcript analysis was performed using a QuantStudio 7 Flex Real-Time PCR System with the Power SYBR Green RNA-to-Ct Real-Time qPCR Kit (all Thermo Fisher Scientific). Total RNA template (25 ng per reaction) was added to the PCR mix, including primers listed below. qPCR conditions were as follows; 48°C for 15 min, 95°C for 10 min followed by 45 cycles (95°C for 15 s, 60°C for 60 s). All qPCR data is collected from at least three independent biological replicates of one experiment. A one-way ANOVA with posthoc Dunnett’s multiple comparisons test was performed on data for the set of targeting vectors for each eGene relative to the scramble control vector. Data analyses were performed using GraphPad PRISM 6 software.

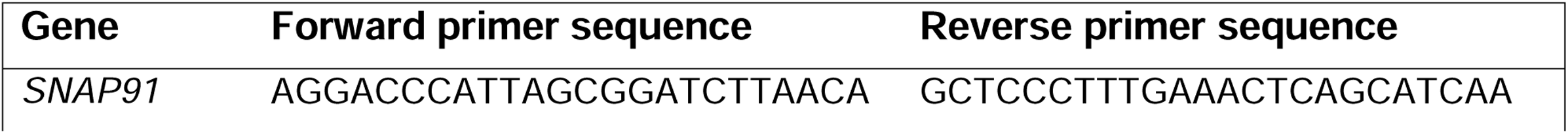
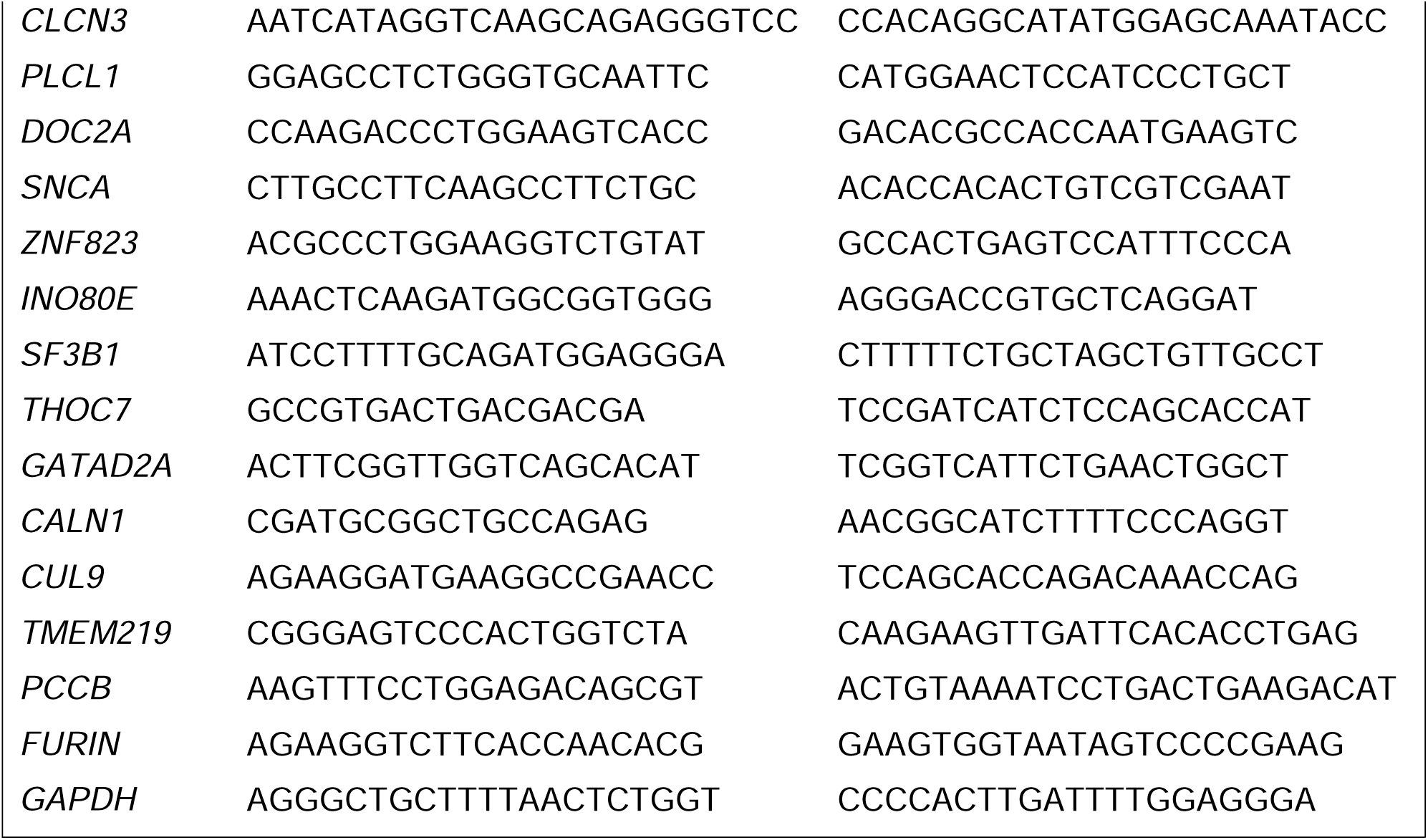

### Immunostaining and high-content imaging microscopy

#### Neurite analysis

Immature iGLUTs were seeded as 1.5×10^4^ cells/well in a 96-well plate coated with 4x Matrigel at day 3. iGLUTs were plated in media containing individual shRNA/gRNA vectors or pools of mixed shRNA and gRNA vectors (Addgene 99374), either targeting eGenes or scramble controls. Medium was switched to non-viral medium four hours post infection. At day 4, transduced iGLUTs were treated with corresponding antibiotics to the gRNA lentiviruses (1 mg/ml HygroB for lentiguide-Hygro/lentiguide-Hygro-mTagBFP2) followed by half medium changes until the neurons were fixed at day 7. At day 7, cultures were fixed using 4% formaldehyde/sucrose in PBS with Ca^2+^ and Mg^2+^ for 10 minutes at room temperature (RT). Fixed cultures were washed twice in PBS and permeabilized and blocked using 0.1% Triton/2% Normal Donkey Serum (NDS) in PBS for two hours. Cultures were then incubated with primary antibody solution (1:1000 MAP2 anti chicken (Abcam, ab5392) in PBS with 2% NDS) overnight at 4°C. Cultures were then washed 3x with PBS and incubated with secondary antibody solution (1:500 donkey anti chicken Alexa 647 (Life technologies, A10042) in PBS with 2% NDS) for 1 hour at RT. Cultures were washed a further 3x with PBS with the second wash containing 1 μg/ml DAPI. Fixed cultures were then imaged on a CellInsight CX7 HCS Platform with a 20x objective (0.4 NA) and neurite tracing analysis performed using the neurite tracing module in the Thermo Scientific HCS Studio 4.0 Cell Analysis Software. 12-24 wells were imaged per condition across a minimum of 2 independent cell lines, with 9 images acquired per well for neurite tracing analysis; each “N” therefore represents an average of hundreds of neurons per image. A one-way ANOVA with a post hoc Bonferroni multiple comparisons test was performed on data for neurite length per neuron using Graphpad Prism.

#### Synapse analyses

Commercially available primary human astrocytes (pHAs, Sciencell, #1800; isolated from fetal female brain) were seeded on D3 at 0.85×10^4^ cells per well on a 4x Matrigel-coated 96 W plate in neuronal media supplemented with 2% fetal bovine serum (FBS). iGLUTs were seeded as 1.5×10^5^ cells/well in a 96-well plate coated with 4x Matrigel at day 5. Half changes of neuronal media were performed twice a week until fixation. At day 13, iGLUTs were treated with 200 nM Ara-C to reduce the proliferation of non-neuronal cells in the culture. At day 18, Ara-C was completely withdrawn by full medium change while adding media containing individual shRNA/gRNA vectors or pools of mixed shRNA and gRNA vectors (Addgene 99374), either targeting eGenes or scramble controls. Medium was switched to non-viral medium four hours post infection. At day 19, transduced iGLUTs were treated with corresponding antibiotics to the gRNA lentiviruses (1 mg/ml HygroB for lentiguide-Hygro/lentiguide-Hygro-mTagBFP2) followed by half medium changes until the neurons were fixed at day 21. At day 21, cultures were fixed and immunostained as described previously, with an additional antibody stain for Synapsin1 (primary antibody: 1:500 Synapsin1 anti mouse (Synaptic Systems, 106 011); secondary antibody: donkey anti mouse Alexa 568 (Life technologies A10037)). Stained cultures were imaged and analyzed as above using the synaptogenesis module in the Thermo Scientific HCS Studio 4.0 Cell Analysis Software to determine SYN1+ puncta number, area, and intensity per neurite length in each image. 20 wells were imaged per condition across a minimum of 2 independent cell lines, with 9 images acquired per well for synaptic puncta analysis. A one-way ANOVA with a post hoc Bonferroni multiple comparisons test was performed on data for puncta number per neurite length using Graphpad Prism.

| Antibody | Species | Supplier | Product number | Dilution |
| --- | --- | --- | --- | --- |
| MAP2 | Ck | Abcam | ab5392 | 1:500 |
| SYNAPSIN1 | Ms | Synaptic Systems | 106 011 | 1:500 |
| Alexa 568 anti-Mouse | Ms | Life technologies | A10037 | 1:500 |
| Alexa 647 anti-Chicken | Ck | Life technologies | A10042 | 1:500 |

#### Multiple Electrode array (MEA)

Commercially available primary human astrocytes (pHAs, Sciencell, #1800; isolated from fetal female brain) were seeded on D3 at 1.7×10^4^ cells per well on a 4x Matrigel-coated 48 W MEA plate (catalog no. M768-tMEA-48W; Axion Biosystems) in neuronal media supplemented with 2% fetal bovine serum (FBS). At D5, iGLUTs were detached, spun down, and seeded on the pHA cultures at 1.5×10^5^ cells per well. Half changes of neuronal media supplemented with 2% FBS were performed twice a week until day 42. At day 13, co-cultures were treated with 200 nM Ara-C to reduce the proliferation of non-neuronal cells in the culture. At Day 18, Ara-C was completely withdrawn by full medium change. At day 25, a full media change was performed to add media containing individual shRNA/gRNA vectors or pools of mixed shRNA and gRNA vectors (Addgene 99374), either targeting eGenes or scramble controls. Medium was switched to non-viral medium four hours post infection. If drug treatments were included, D26 neurons were treated for 48hrs with either Anandamide (10µM), Etomoxir (10µM), Simvastatin (10µM), or matched vehicles. Electrical activity of iGLUTs was recorded at 37°C twice every week from day 28 to day 42 using the Axion Maestro MEA reader (Axion Biosystems). Recording was performed via AxiS 2.4. Batch mode/statistic compiler tool was run following the final recording. Quantitative analysis of the recording was exported as Microsoft excel sheet. Data from 6-12 biological replicates were analyzed using GraphPad PRISM 6 software or R.

#### RNAseq

RNA Sequencing libraries were prepared using the Kapa Total RNA library prep kit. Paired-end sequencing reads (100bp) were generated on a NovaSeq platform. Raw reads were aligned to hg19 using STAR aligner^147^ (v2.5.2a) and gene-level expression were quantified by featureCounts^148^ (v1.6.3) based on Ensembl GRCh37.70 annotation model. Genes with over 10 counts per million (CPM) in at least four samples were retained. After filtering, the raw read counts were normalized by the voom^149^ function in limma and differential expression was computed by the moderated t-test implemented in limma^150^. Differential gene expression analysis was performed between each CRISPRa/shRNA target group and scramble control group. Bayes shrinkage (limma::eBayes) estimated modified t- and p-values and identified differentially expressed genes (DEGs) based on an FDR <= 0.05 (limma::TopTable)^151^. Gene Ontology/pathways were evaluated using Gene-set Enrichment Analysis (GSEA)^152^, with genes expressed in iGLUTs as our baseline comparison. In these analyses, the t-test statistics from the differential expression contrast were used to rank genes in the GSEA using the R package ClusterProfiler^153^. Permutations (up to 100,000 times) were used to assess the GSEA enrichment P value. Log2 fold changes in expression were calculated across all RNA-seq samples in our arrayed dataset.

#### Analysis of additive and non-additive effects

We applied our published approach to resolve distinct additive and non-additive transcriptomic effects after combinatorial manipulation of genetic variants and/or chemical perturbagens, developed^27^, applied^59^, and described in detail^83^. The expected additive effect was modeled through addition of the individual comparisons; the non-additive effect was modeled by subtraction of the additive effect from the combinatorial perturbation comparison. Fitting of this model for differential expression identifies genes that show a difference in the expected differential expression computed for the additive model compared to the observed combinatorial perturbation. Briefly, the non-additive effect between eGenes was identified using a limma’s linear model analysis. The coefficients, standard deviations and correlation matrix were calculated, using *contrasts.fit,* in terms of the comparisons of interest. Empirical Bayes moderation was applied using the *eBayes* function to obtain more precise estimates of gene-wise variability. P-values were adjusted for multiple hypotheses testing using false discovery rate (FDR) estimation, and differentially expressed genes were determined as those with FDR ≤ 10%, unless stated otherwise. Two methods were used to compare the extent of synergy between data sets. First, we calculated the fraction of synergistic genes (FDR<10%) measure the extent of synergy. Second, we calculated a synergy coefficient, π1, as the fraction of non-null synergistic P-values, to inform the existence of a synergistic component, even if the P-values themselves are not significant genome-wide.

However, interpretation of the resulting DEGs depends on several factors, such as the direction of fold change (FC) in all three models. To identify genes whose magnitude of change is larger in the combinatorial perturbation vs. the additive model, we categorized all genes by the direction of their change in both models and their log_2_(FC) in the non-additive model. First, log_2_(FC) standard errors (SE) were calculated for all samples. Genes were then grouped into ‘positive non-addition’ if their FC was larger than SE and ‘negative non-addition’ if smaller than -SE. If the corresponding additive model log_2_(FC) showed the same or no direction, the gene was classified as “more” differentially expressed in the combinatorial perturbation than predicted. GSEA was performed on a curated subset of the MAGMA collection using the limma package camera function, which tests if genes are ranked highly in comparison to other genes in terms of differential expression, while accounting for inter-gene correlation. Due to the small sample size in this study and moderate fold changes in some eGene perturbations, changes in gene expression may be small and distributed across many genes. However, powerful enrichment analyses in the limma package may be used to evaluate enrichment based on genes that are not necessarily genome-wide significant and identify sets of genes for which the distribution of t-statistics differs from expectation. Over-representation analysis (ORA) was performed when subsets of DEGs were of interest; genes of interests were ranked by –log10 (p-value) and enrichment was performed against a background of all expressed genes using the WebGestaltR package.

#### Dataset for population-level analysis of synergy

Individuals from the Sweden-SCZ Population-Based cohort were obtained from the database of Genotypes and Phenotypes, Study Accession: phs000473.v2.p2 (N_Cases_ = 5,232, N_Controls_ = 6,468)^154^.

#### Pathway polygenic risk scores

Pathway-specific polygenic risk score (PRS) analyses were performed using PRSice-2 (v2.3.5) on individual genotype data for the Sweden-SCZ Population based cohort. A total of 4,834 individuals diagnosed with SCZ and 6,128 controls were included after quality control. To calculate the scores, we used a version of the summary statistics from the PGC SCZ GWAS that excludes the Sweden-SCZ data to prevent inflation of results. SNPs were annotated to genes and pathways based on GTF files obtained from ENSEMBL (GRCh37.75). To include potential gene regulatory elements, gene coordinates were extended 35 kilobases (kb) upstream and 10 kb downstream of each gene. We excluded from analyses the MHC region (chr6:25Mb-34Mb), ambiguous SNPs (A/T and G/C), and SNPs not present in both GWAS summary statistics and genotype data.

To obtain empirical “competitive” *P-*values, that assess GWAS signal enrichment while accounting for pathway size, we performed the following permutation procedure: first, a “background” pathway containing all genic SNPs is constructed, and clumping is performed within this pathway. For each pathway with *m* SNPs, *N=*10,000 null pathways are generated by randomly selecting *m* SNPs from the “background” pathway. The competitive *P*-value can then be calculated as

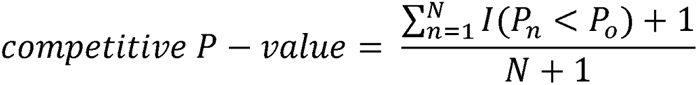

where *I*(.) is an indicator function, taking a value of 1 if the association *P*-value of the observed pathway (*P_0_*) is larger than the one obtained from the *n*^th^ null pathway (*P_n_*), and 0 otherwise (see^84^ for additional details).

For the analyses testing whether non-additive genes from synaptic/regulatory pathways explain larger R^2^ than *<u>the same number</u>* of non-additive genes from random combinations (**SI Fig. 24**), we took 2,799 random genes from the non-additive synaptic and regulatory transcriptome, which corresponds to the number of genes with non-additive effects in one of the random joint perturbations. For the GTF NULL permutation analyses, we selected n=2,799 random genes from the GTF file GRCh37.75. Pathway-specific PRS for each sample of 2,799 genes was calculated using PRSet^84^, as described above. This procedure was repeated 1000 times.

#### Transcriptomic Risk Score (TRS) Analyses

In order to test the impact of non-additive genetic effects *in silico*, we used transcriptomic imputation methods to calculate genetically-regulated gene expression (GREX) for individuals from the Sweden-SCZ Population-Based cohort (**SI Table 3**). Brain GREX was calculated using PrediXcan^73^ with CMC dorsolateral prefrontal cortex (CMC-DLPFC) models^6^. Predicted GREX levels were calculated for the fifteen eGenes. An initial test of aberrant gene expression was performed by counting the number of genes with dysregulated GREX (defined as predicted GREX in the top or bottom decile of overall expression of that gene, defined in the direction of effect of that gene’s association with SCZ from S-PrediXcan analyses (top decile for positive effect, bottom decile for negative effect) for each of the five-gene groups (synaptic, regulatory, multi-function), and summed the number of aberrant genes present in each individual for each perturbed gene group (Synaptic, Regulatory, and Multi-function). We then looked at the SCZ case/control proportion within each group of individuals with 3+, 1-2, and any genes with aberrant GREX.

#### Association of Synaptic, Regulatory, and Multi-function gene-sets with SCZ

We tested for association of each of the fifteen eGene GREX individually with SCZ (SCZ ∼ GREX), and then calculated composite scores of group GREX (Synaptic, Regulatory, and Multi-function) using a Transcriptomic Risk Score (TRS), calculated as the sum of each GREX weighted by the direction of gene perturbation (1 for activation, -1 for inhibition) from *in vivo* experiments, divided by the total number of genes (N) in the gene-set (Equation X):

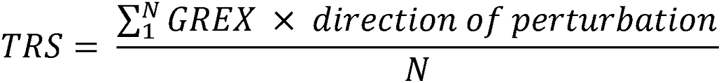

We then tested for association of each TRS (Synaptic, Regulatory, and Multi-function) with SCZ status in the Swedish cohort.

#### Permutation tests

We performed permutation tests to assess the impact of (1) the number of genes included in our TRS gene group and (2) the number of pathways impacted by those genes on SCZ case status. We used S-PrediXcan to find genes with CMC-DLPFC GREX associated with SCZ in a large SCZ cohort (N_Cases_ = 11,260, N_Controls_ = 24,542)^39^. From this resulting list of genes, we assigned genes to two groups: nominally-significant genes (N=1,963, Bonferroni p<0.05), and tissue-specific significant genes (N=144, p<0.05/N_Genes in CMC-DLPFC PrediXcan model_). We created pathway sets affected by these genes using the overlap with Kyoto encyclopedia of genes and genomes (KEGG)^155^ and gene ontology (GO)^156,157^. This gave us a sampling pool of 1,465 genes affecting 8,324 pathway sets for the nominally-significant group, and 110 genes affecting 2,382 pathway sets for the tissue-specific group. We then performed permutation sampling analyses (for nominally-significant and tissue-specific significant gene-pathway set pools) where we randomly sampled sets of five, ten, or fifteen genes from the sampling pool (adjusted for the size of each pathway set), calculated TRS from the sampled gene-set, and looked at the association of TRS with SCZ. We performed sampling 100,000 times for each gene-set size. For this analysis, TRS was calculated by taking the sum of each gene in the gene-sets GREX weighted by the direction of effect of the gene association with SCZ from our S-PrediXcan analysis (1 or -1) (Equation X):

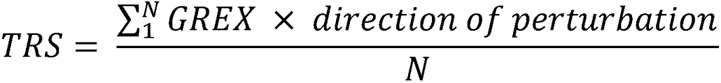

We then looked at the overall association the number of pathways hit by each TRS (based on the annotated lists) with SCZ variance explained (**SI Fig. 1A-C**). To determine if the type of pathways hit by our perturbed genes was important to SCZ risk (i.e. is it more important to hit multiple, similar pathways or more diverse pathways to increase SCZ variance explained), we additionally assessed whether the similarity in make-up of pathways affected by the TRS was associated with SCZ. To do this, we used the R “GeneOverlap” package to calculate the average Jaccard Index of pathways for each TRS, and looked at the association of that index with SCZ.

## STATEMENT OF ETHICS

Yale University Institutional Review Board waived ethical approval for this work. Ethical approval was not required because the hiPSC lines, lacking association with any identifying information and widely accessible from a public repository, are thus not considered to be human subjects research. Post-mortem data are similarly lacking identifiable information and are not considered human subjects research.

## AUTHOR CONTRIBUTIONS

SCZ eGene lists were prioritized by E.S., L.H., W.Z., G.V. and P.R, together with K.J.B. Epigenome data provided by K.G., S.A. and P.R. iGLUT transcriptomic and phenotypic studies were conducted by P.J.M.D, with assistance by E.C. Synergy analysis was conceptualized by N.S. and applied by P.J.M.D; convergent analyses were conceptualized by K.G.T. and applied by C.S. Transcriptomic imputation was conducted by J.J. and L.H.; pathway-specific PRS by J.G.G. and P.O. The ECCITE-seq pipeline was adapted to hiPSC-neurons by A.L., supported by P.J.M.D., J.F., A.Y. and S.C. ECCITE-seq iGLUT neuron studies were conducted by A.L. and P.J.M.D. 10x analyses were conducted by J.F, and preliminary ECCITE-seq quality control conducted by MW. Convergent analyses were conceptualized by K.J.B. and L.H., developed and conducted entirely by K.G.T. The paper was written by P.J.M.D, K.G.T., L.H., and K.J.B., with input from all authors.

Special thanks to Michael Talkowski and Douglas Ruderfer for countless discussions on convergence.

## DATA AND CODE AVAILABILITY

All source donor hiPSCs have been deposited at the Rutgers University Cell and DNA Repository (study 160; http://www.nimhstemcells.org/); dCas9-VPR hiPSCs are in the process of being submitted in advance of publication.

The accession number for sc-RNA sequencing data reported in this paper is Gene expression omnibus (GEO): GSE200774. Processed data and accompanying code can be accessed through Synapse (syn27819129)

