## Supplemental Figures & Tables for "Convergent and non-additive impact of schizophrenia risk genes in human neurons"

#### SUPPLEMENTAL INFORMATION

**Box 1.** *Summary of additive and non-additive effects of joint eGene perturbation on transcription, related to Figure 4, Supplemental Figures 21-23*.

**Box 2**. *In vitro validation of in silico drug predictions, related to Figure 6, Supplemental Figures 28-30.*

**Supplemental Figure 1.** *Prioritization and manipulation of synaptic, regulatory, and multi-function brain eGenes regulated by SCZ, related to Figure 1.*

**Supplementary Figure 2.** *Degree of variance in NGN2-induction is not significantly different between Day 7 and Day 21 iGLUTs.*

**Supplementary Figure 3.** *Degree of heterogeneity of iGLUTs is not significantly correlated with perturbation identity.*

**Supplementary Figure 4.** *Weighted Nearest Neighbor (WNN) dimensional reduction identifies successfully perturbed cells.*

**Supplementary Figure 5.** *Multi-modal analysis allows for identification of successfully perturbed cells by dimensional reduction and weighted clustering in D7 iGLUTs.*

**Supplementary Figure 6.** *Multi-modal analysis allows for identification of successfully perturbed cells by dimensional reduction and weighted clustering in D21 iGLUTs.*

**Supplemental Figure 7.** *Successful CRISPR-activation of top SCZ eGenes compared to predicted effects of eQTL causal SNPs.*

**Supplemental Figure 8.** *Successful CRISPR-activation of top SCZ eGenes leads to significant changes in gene expression.*

***Supplemental Figure 9.*** *Downstream target-convergent networks identified by Bayesian bi-clustering resolve distinct networks enriched for SCZ common and rare variant target genes and transcription factor binding motifs, related to Figure 2, SI Data 3.*

**Supplemental Figure 10**. *Functional scoring of perturbation sets***,** *related to Figure 3, SI figure 11.*

**Supplementary Figure 11.** *The degree of network convergence is influenced by functional similarity of target perturbations***,** *related to Figure 3, SI figure 10.*

**Supplementary Figure 12.** *Gene expression of D7 and D21 NGN2-neurons significantly correlate with gene expression of the postmortem adult DLPFC.*

**Supplementary Figure 13.** *Gene expression of D7 and D21 NGN2-neurons significantly correlate with gene expression of adult cortical neurons and fetal trans and excitatory neurons.*

**Supplementary Figure 14.** *Basal gene expression of SCZ eGenes perturbed in this study in D7 and D21 NGN2-neurons and fetal and adult cell-types in the prefrontal cortex.*

**Supplementary Figure 15.** *Concordance of differential gene expression patterns across individual gRNAs targeting CALN1.*

**Supplementary Figure 16.** *Similarity in molecular function and degree of perturbation of synaptic and regulatory genes drives non-additivity.*

**Supplementary Figure 17.** *Across ~100 SCZ risk genes, a median of 10 were classified as perturbed per individual.*

**Supplementary Figure 18.** *Across 20 SCZ risk genes targeted in the arrayed and pooled screens, an average of 2 and a maximum of 8 were classified as perturbed per individual.*

**Supplemental Figure 19.** *Perturbation of SCZ eGenes results in differential expression of genes relating to brain disorders and synaptic function (representative genes), related to Figure 1.*

**Supplemental Figure 20.** *Perturbation of SCZ eGenes results in differential expression of genes relating to brain disorders and synaptic function (comprehensive analysis), related to Figure 1.*

**Supplemental Figure 21.** *Combinatorial perturbation of synaptic and regulatory eGenes results in non-additive effects on transcription, related to Figure 4.*

**Supplemental Figure 22.** *Comparison of non-additive effects of joint perturbations using combined single vectors vs multiplexed vector, related to Figure 4.*

**Supplemental Figure 23.** *Combinatorial perturbation of random eGenes results in non-additive impacts on transcription which scale with the number of perturbed eGenes, related to Figure 4.*

**Supplemental Figure 24.** *Within-pathway non-additive genes are associated with SCZ risk.*

**Supplemental Figure 25.** *Combinatorial perturbation of SCZ eGenes with synaptic functions results in impaired neurite outgrowth, synaptic expression, and neuronal hyperactivity.*

**Supplemental Figure 26.** *Detailed analysis of effect of combinatorial perturbation of SCZ eGenes on impaired neurite outgrowth and synaptic expression, related to SI Figure 25, Figure 4.*

**Supplemental Figure 27***. Detailed analysis of effect of combinatorial perturbation of SCZ eGenes on neuronal hyperactivity, related to SI Figure 25, Figure 4***.**

**Supplementary Figure 28.** *In vitro validation of drug-eGene interactions at the transcriptomic level demonstrate an opposing effect of the predicted drugs on eGene activation***,** *related to Figure 6.*

**Supplementary Figure 29.** *Comparison of the transcriptomic effects of eGene perturbations alone and the combined drug-eGene perturbations, related to Figure 6.*

**Supplementary Figure 30***. In vitro validation of drug-eGene interactions on neuronal morphology, related to Figure 6.*

**Supplementary Figure 31.** *Convergence across different eGenes resolve distinct networks that may inform molecular subtypes related to schizophrenia.*

**Supplementary Figure 32.** *K-means clustering resolution influences identification or diagnostic subgroups defined by eGene patterns of expression.*

**Supplemental Table 1**. Selection of PGC3 SCZ-GWAS eGenes. Top 25 genes prioritized by COLOC further informed by PrediXcan (grey indicates not significant by PrediXcan.) Selected synaptic (green), regulatory/epigenetic (blue), and multi-function (purple) eGenes indicated; multi-function genes were chosen to limit any potential overlapping function.

**Supplemental Table 2.** Effect sizes and p-values of eGenes from predicted, arrayed, and pooled methods corresponding *to Figure 1B-D.*

**Supplemental Table 3.** Prioritization and manipulation of synaptic, regulatory, and multi-function brain eGenes regulated by SCZ, *related to Figure 1*.

**Supplemental Table 4.** List of perturbed eGenes, no. of non-additive genes at each FDR cutoff and no. of convergent genes for each combinatorial perturbation condition, *related to Figures 4 & 5.*

**Supplemental Data 1.** *DEGs, GSEA tables, synergy sub-categories, and synergy sub-category over-representation analysis for arrayed screen RNA-seq data.*

**Supplemental Data 2.** *Individual scRNA-seq Perturbation DEGs and Pathway Enrichments from pooled experiment.*

**Supplemental Data 3.** *Reconstructed Convergent Networks and Convergent network enrichment results (FUMA, ClusterProfiler, ORA of common/rare/variants) from arrayed and pooled screens.*

**Supplemental Data 4.** *CMAP drug prioritization queries and GSEA for 10 target and each individual perturbation signature used in CMAP query***.**

**Supplemental Data 5.** *Z-scored expression of targeted eGenes in the CMC DLPFC.*

#### SUPPLEMENTAL TABLE

**SI Table 1.** SCZ eGene selection. Highlighted in bold are eGenes prioritized as synaptic (green), regulatory/epigenetic (blue), and unrelated multi-function (purple). (Grey indicates not significant by PrediXcan.)

| | GWAS | | | | | COLOC | PrediXcan | |
|---|---|---|---|---|---|---|---|---|
| Function | Chr | SNP | P | LD range | Gene | PP4 | z | P |
| proprotein convertase | 15 | rs4702 | 2.15E-23 | 90414642…92426654 | ***FURIN*** | 0.999943 | -10.06869 | 7.60E-24 |
| regulatory | 19 | rs72986630 | 3.07E-10 | 10832225…12839819 | ***ZNF823*** | 0.999253 | 6.350252 | 2.15E-10 |
| signalling | 1 | rs56335113 | 3.15E-15 | 28563084…30653243 | *PTPRU* | 0.985465 | -6.850981 | 7.33E-12 |
| metabolism | 8 | rs10957321 | 4.18E-10 | 64500933…66710532 | *CYP7B1* | 0.978623 | -6.202272 | 5.57E-10 |
| development | 17 | rs6504163 | 1.87E-09 | 60555375…62597717 | *ACE* | 0.967474 | -5.869283 | 4.38E-09 |
| pseudogene | 6 | rs2153960 | 9.22E-10 | 108103765…110108063 | *ZNF259P1* | 0.961803 | -5.466066 | 4.60E-08 |
| ubiquitination | 6 | rs113113059 | 2.29E-11 | 42150013…44192158 | ***CUL9*** | 0.953677 | 5.754229 | 8.70E-09 |
| calcium signalling | 7 | rs2944821 | 1.90E-09 | 70244499…72912040 | ***CALN1*** | 0.913966 | 5.547481 | 2.90E-08 |
| signalling | 14 | rs722637 | 9.64E-15 | 103201875…105313861 | *PPP1R13B* | 0.913695 | -5.132138 | 2.86E-07 |
| signalling | 13 | rs9545047 | 3.05E-11 | 79057053…81130204 | *NDFIP2* | 0.907569 | -6.536768 | 6.29E-11 |
| metabolism | 3 | rs7432375 | 5.32E-15 | 134969406…137055316 | ***PCCB*** | 0.906413 | -7.547429 | 4.44E-14 |
| chromatin remodeling | 16 | rs3814883 | 8.82E-15 | 29007489…31016970 | ***INO80E*** | 0.894347 | 7.639276 | 2.18E-14 |
| RNA transport | 3 | rs704364 | 8.41E-10 | 62819885…64848612 | ***THOC7*** | 0.893413 | -5.236222 | 1.64E-07 |
| synapse | 6 | rs2022265 | 3.74E-10 | 83262613…85419243 | ***SNAP91*** | 0.893265 | 5.185282 | 2.16E-07 |
| protein binding | 1 | rs61828917 | 7.95E-10 | 172580537…174637937 | *ANKRD45* | 0.891423 | 3.614873 | 0.0003 |
| splicing | 2 | rs2914983 | 1.10E-14 | 197256700…199299078 | ***SF3B1*** | 0.888092 | 7.256793 | 3.96E-13 |
| signalling | 3 | rs75968099 | 5.16E-11 | 35754053…37781278 | *DCLK3* | 0.88304 | -4.628441 | 3.68E-06 |
| DNA repair | 3 | rs75968099 | 5.16E-11 | 36034966…38107017 | *MLH1* | 0.877109 | 6.376857 | 1.81E-10 |
| metabolism | 11 | rs58950470 | 1.10E-08 | 64482546…66486143 | *RNASEH2C* | 0.869117 | -5.203533 | 1.96E-07 |
| regulatory | 1 | rs11121172 | 7.15E-10 | 7412645…9876635 | *RERE* | 0.867634 | 2.078589 | 0.037655 |
| regulatory | 19 | rs1858999 | 7.97E-14 | 18497024…20619093 | ***GATAD2A*** | 0.858178 | -7.214675 | 5.41E-13 |
| synapse | 4 | rs356183 | 3.37E-08 | 89645368…91759130 | ***SNCA*** | 0.847582 | -4.494898 | 6.96E-06 |
| signalling | 16 | rs3814883 | 8.82E-15 | 28952638…30984212 | ***TMEM219*** | 0.846736 | 6.291546 | 3.14E-10 |
| pseudogene | 2 | rs6546857 | 2.80E-09 | 72872890…74911073 | *ALMS1P* | 0.830632 | 5.282995 | 1.27E-07 |
| ubiquitination | 12 | rs4766428 | 2.61E-17 | 109813508…111837285 | *ANAPC7* | 0.824378 | -0.962298 | 0.3359 |
| synapse | 16 | rs3814883 | 8.82E-15 | 30016830…30034591 | ***DOC2A*** | 0.521296 | 6.088335 | 1.14E-09 |
| synapse | 4 | rs61405217 | 5.39E-11 | 170533784…170644824 | ***CLCN3*** | 0.718835 | 5.771783 | 7.84E-09 |
| synapse | 2 | rs1451488 | 6.72E-17 | 198669426…199437305 | ***PLCL1*** | 0.035058 | 4.910285 | 9.09E-07 |

**SI Table 2:** Effect sizes and p-values of eGenes from predicted, arrayed, and pooled methods corresponding to **Figure 1B-D**. A larger version of this table can be found in the excel file with the same name.

| GENE | Assay | CALN1 | CLCN3 | CUL9 | DOC2A | FES | FURIN | GATAD2A | INO80E | NAGA | PCCB | PLCL1 | SF3B1 | SNAP91 | SNCA | THOC7 | TMEM219 | UBE2Q2L | ZNF804A | ZNF823 |
|---|---|---|---|---|---|---|---|---|---|---|---|---|---|---|---|---|---|---|---|---|
| avgEXP single | Arrayed - Single Perturbations | 1.326890191 | 1.2242146 | 1.05582609 | 1.279257883 | | 0.85683871 | 0.85085943 | 1.06121914 | | 0.54680778 | 1.30992171 | 1.15827609 | 1.353036456 | 0.63439452 | 0.8195733 | 1.17711224 | | | 1.19400766 |
| logFC single | Arrayed - Single Perturbations | 0.408048983 | 0.29185648 | 0.07837222 | 0.355307124 | | -0.2229044 | -0.2330073 | 0.08572259 | | -0.8708943 | 0.38948059 | 0.21197917 | 0.436200711 | -0.6565478 | -0.2870551 | 0.2352519 | | | 0.2558121 |
| pVal single | Arrayed - Single Perturbations | 0.091550744 | 0.00062434 | 0.16617555 | 0.000573473 | | 0.04062416 | 0.01919266 | 0.17557752 | | 7.3997E-05 | 0.02269349 | 0.03988319 | 0.037725271 | 0.04374504 | 0.05402839 | 0.10534946 | | | 0.0060221 |
| aveEXP multiplex | Arrayed - Functional Combinatorial | 1.417515372 | 1.22894618 | 0.87702935 | 1.489726178 | | 0.88928338 | 0.55742356 | 1.03769214 | | 0.36690876 | 1.42838328 | 1.11943569 | 1.168488328 | 0.37103278 | 0.72975383 | 1.17172222 | | | 1.20621332 |
| logFC multiplex | Arrayed - Functional Combinatorial | 0.50336438 | 0.29742174 | -0.189303 | 0.575047178 | | -0.1692849 | -0.8431541 | 0.05337849 | | -1.4465067 | 0.51438315 | 0.16277165 | 0.224643324 | -1.4303814 | -0.4545182 | 0.22863059 | | | 0.27048507 |
| pVal multiplex | Arrayed - Functional Combinatorial | 0.017699985 | 8.248E-05 | 0.01753236 | 0.000771384 | | 0.05677642 | 0.00016711 | 0.16036789 | | 0.00054478 | 0.00548584 | 0.03527222 | 0.017363832 | 0.00088377 | 0.01380475 | 0.05553635 | | | 0.03916232 |
| aveEXP random | Arrayed - Random Combinatorial | 1.426486024 | 1.0624696 | 0.88324314 | 1.206167628 | | 0.83641821 | 0.52746289 | 1.006443 | | 0.39062242 | 1.44614879 | 1.08914284 | 1.125290714 | 0.35313322 | 0.75450864 | 1.12955589 | | | 1.26198769 |
| logFC random | Arrayed - Random Combinatorial | 0.512465612 | 0.08742156 | -0.1791175 | 0.270430421 | | -0.2577036 | -0.9228585 | 0.00926547 | | -1.3561533 | 0.532216 | 0.12319318 | 0.170297764 | -1.5017155 | -0.4063907 | 0.17575566 | | | 0.33569784 |
| pVal random | Arrayed - Random Combinatorial | 0.002787659 | 0.02136499 | 0.02710188 | 0.066144203 | | 0.01844009 | 0.00053979 | 0.44182476 | | 0.00011511 | 0.00740362 | 0.02703743 | 0.072358678 | 0.00081174 | 0.05822668 | 0.01601603 | | | 0.0038822 |
| aveEXP all15 | Arrayed - All Combinatorial | 1.374260825 | 1.22224843 | 0.92870175 | 1.582078056 | | 1.06226865 | 0.61359371 | 1.08443611 | | 0.41736776 | 1.42350033 | 1.16000986 | 1.180447533 | 0.3790281 | 0.87059165 | 1.18800132 | | | 1.16904556 |
| logFC all15 | Arrayed - All Combinatorial | 0.458655843 | 0.28953755 | -0.1067127 | 0.661820781 | | 0.08714867 | -0.7046444 | 0.11694505 | | -1.2606089 | 0.50944283 | 0.21413707 | 0.23933392 | -1.3996233 | -0.1999319 | 0.24853644 | | | 0.22533115 |
| pVal all15 | Arrayed - All Combinatorial | 0.008272496 | 2.4318E-05 | 0.07761502 | 0.008191476 | | 0.06997431 | 0.00040737 | 0.11313436 | | 0.00017466 | 0.0171118 | 0.00760967 | 0.004698015 | 0.00076735 | 0.10690761 | 0.03992569 | | | 0.00476586 |
| CIRSPRa_1vAll_avgExpr | Pooled - 1 gRNA vs All other gRNAs | 1.021856312 | 3.6690882 | | | 1.94781735 | | | | 1.88230086 | | 0.93439928 | 9.55186681 | | | | 5.83895365 | 0.59668856 | 1.92216903 | 0.55518047 |
| CIRSPRa_1vAll_logFC | Pooled - 1 gRNA vs All other gRNAs | 0.848550656 | 1.322297 | | | 1.77470725 | | | | 1.40469839 | | 0.72426524 | 0.9065338 | | | | 3.43631815 | 0.42829866 | 1.4954021 | 0.19686882 |
| CIRSPRa_1vAll_pval | Pooled - 1 gRNA vs All other gRNAs | 4.58E-05 | 0.0002309 | | | 4.05E-11 | | | | 1.70E-07 | | 0.00044931 | 1.24E-17 | | | | 4.91E-17 | 0.01219014 | 9.35E-08 | 0.13253058 |
| CIRSPRa_1vAll_padj | Pooled - 1 gRNA vs All other gRNAs | 0.385617465 | 0.04645993 | | | 6.83E-07 | | | | 0.00286224 | | 0.14285567 | 6.30E-16 | | | | 4.86E-14 | 0.9341992 | 0.00157624 | 0.25129658 |
| CRISPRa_1vScram_avgExpr | Pooled - 1 gRNA vs Scramble gRNA c | 1.021856312 | 3.6690882 | | | 1.94781735 | | | | 1.88230086 | | 0.93439928 | 9.55186681 | | | | 5.83895365 | 0.59668856 | 1.92216903 | 0.55518047 |
| CRISPRa_1vScram_logFC | Pooled - 1 gRNA vs Scramble gRNA c | 0.835398169 | 0.93324216 | | | 1.61905306 | | | | 1.30160873 | | 0.84798869 | 1.87182196 | | | | 3.51100258 | 0.40859644 | 1.52545299 | 0.37027793 |
| CRISPRa_1vScram_pval | Pooled - 1 gRNA vs Scramble gRNA c | 4.16E-24 | 0.00279192 | | | 3.04E-47 | | | | 2.62E-29 | | 8.32E-30 | 1.83E-57 | | | | 1.89E-20 | 2.89E-06 | 5.63E-37 | 1.48E-05 |
| CRISPRa_1vScram_padj | Pooled - 1 gRNA vs Scramble gRNA c | 7.02E-20 | 0.23175641 | | | 5.12E-43 | | | | 4.42E-25 | | 3.51E-26 | 1.90E-55 | | | | 1.59E-16 | 0.0243815 | 9.49E-33 | 5.84E-05 |
| S.PrediXcan_Zscore | Predicted | -5.54748126 | -5.7717826 | -5.7542287 | -6.08826863 | -6.5874518 | 10.0686914 | 7.21467511 | -7.6392764 | -6.8493887 | 7.54742906 | -4.9102854 | -7.2567934 | -5.185282301 | 4.49489825 | 5.23622222 | -6.2915463 | | | -6.3502525 |
| S.PrediXcan_Effect | Predicted | -0.70123495 | -0.4391492 | -0.291785 | -0.468836561 | -0.3057547 | 0.3848464 | 0.16564488 | -0.2937592 | -0.1980673 | 0.19935838 | -0.7706139 | -0.6119333 | -0.22317725 | 0.24194434 | 0.20844776 | -0.4253221 | | | -0.6107145 |
| S.PrediXcan_pval | Predicted | 2.90E-08 | 7.84E-09 | 8.70E-09 | 1.14E-09 | 4.47E-11 | 7.60E-24 | 5.41E-13 | 2.18E-14 | 7.42E-12 | 4.44E-14 | 9.09E-07 | 3.96E-13 | 2.16E-07 | 6.96E-06 | 1.64E-07 | 3.14E-10 | | | 2.15E-10 |
| EpiXcan_zscore | Predicted | 6.569210514 | 5.9912689 | 5.99003637 | 5.692455809 | 6.63188602 | -9.0837964 | -7.2032957 | 7.0995603 | 6.6697758 | -7.6450732 | 7.19984476 | 7.26505287 | 6.164942734 | -4.5969384 | -5.5065661 | 7.24046319 | 6.88921821 | | 5.24913999 |
| EpiXcan_effect_size | Predicted | 0.195827388 | 0.17535088 | 0.10333009 | 0.195494208 | 0.16386562 | -0.2048402 | -0.0730481 | 0.11931399 | 0.08489799 | -0.0865378 | 0.3168569 | 0.12979483 | 0.126558919 | -0.1893319 | -0.0651137 | 0.3539311 | 0.33313469 | | 0.16657278 |
| EpiXcan_pvalue | Predicted | 5.06E-11 | 2.08E-09 | 2.10E-09 | 1.25E-08 | 3.31E-11 | 1.05E-19 | 5.88E-13 | 1.25E-12 | 2.56E-11 | 2.09E-14 | 6.03E-13 | 3.73E-13 | 7.05E-10 | 4.29E-06 | 3.66E-08 | 4.47E-13 | 5.61E-12 | | 1.53E-07 |

**SI Table 3.** Prioritization and manipulation of synaptic, regulatory, and multi-function brain eGenes regulated by SCZ, related to **Figure 1.**

Predicted GREX levels in dorsolateral prefrontal cortex (DLPFC) calculated for the fifteen eGenes in a Swedish SCZ cohort. Aberrant expression of eGenes is predicted to impact SCZ case-control status in a dose-dependent manner. Number of SCZ cases with 0, 1, 2, 3, 4, or 5 aberrantly expressed eGenes as follows: synaptic (3150, 1509, 290,26, 1, 0), regulatory (1925, 2040, 852, 139, 18, 2), and multi-pathway (3084, 1555, 300, 37, 0, 0).

**Synaptic SCZ GREX**

| # "perturbed" eGenes | 0 | 1 | 2 | 3 | 4 | 5 |
|---|---|---|---|---|---|---|
| CONTROL | 3742 | 1951 | 398 | 45 | 2 | 0 |
| CASE | 3150 | 1509 | 290 | 26 | 1 | 0 |

**Regulatory SCZ GREX**

| # "perturbed" eGenes | 0 | 1 | 2 | 3 | 4 | 5 |
|---|---|---|---|---|---|---|
| CONTROL | 2233 | 2591 | 1109 | 182 | 23 | 0 |
| CASE | 1925 | 2040 | 852 | 139 | 18 | 2 |

**Multi-function SCZ GREX**

| # “perturbed” eGenes | 0 | 1 | 2 | 3 | 4 | 5 |
|---|---|---|---|---|---|---|
| CONTROL | 3622 | 2019 | 441 | 55 | 1 | 0 |
| CASE | 3084 | 1555 | 300 | 37 | 0 | 0 |

**SI Table 4.** List of perturbed eGenes, no. of non-additive genes at each FDR cutoff and no. of convergent genes for each combinatorial perturbation condition, related to **Figures 4 & 5.** A larger version of this table can be found in the excel file with the same name.

| Geneset name | Perturbed eGenes | No. DEGs additive model (FDR<0.1) | Proportion of expressed genes | No. DEGs joint perturbation (FDR<0.1) | Proportion of expressed genes | No. non-additive DEGs (FDR<0.1) | Proportion of expressed genes | No. non-additive DEGs (FDR<0.05) | Proportion of expressed genes | No. non-additive DEGs (FDR<0.01) | Proportion of expressed genes | Synergy coefficient ($\pi$) | % Convergent DEGs |
|---|---|---|---|---|---|---|---|---|---|---|---|---|---|
| Synaptic | SNCA, SNAP91, CLCN3, DOC2A, PLCL1 | 7857 | 30.27 | 4649 | 17.91 | 4396 | 16.93 | 2726 | 10.50 | 720 | 2.77 | 43.86 | 3.73 |
| Regulatory | THOC7, GATAD2A, ZNF823, INO80E, SF3B1 | 6876 | 26.49 | 1498 | 5.77 | 5249 | 20.22 | 3690 | 14.21 | 1340 | 5.16 | 42.74 | 3.46 |
| Multi-function | FURIN, PCCB, CALN1, CUL9, TMEM219 | 22 | 0.08 | 723 | 2.79 | 1 | 0.00 | 1 | 0.00 | 0 | 0.00 | 0.00 | 0.20 |
| 5 Random 1 | SNCA, THOC7, SNAP91. ZNF823, CALN1 | 7156 | 27.57 | 4611 | 17.76 | 2799 | 10.78 | 1353 | 5.21 | 110 | 0.42 | 38.80 | 4.82 |
| 5 Random 2 | PCCB, CLCN3, DOC2A, CUL9, SF3B1 | 3283 | 12.65 | 83 | 0.32 | 324 | 1.25 | 2 | 0.01 | 0 | 0.00 | 30.30 | 1.05 |
| 5 Random 3 | FURIN, GATAD2A, PLCL1, INO80E, TMEM219 | 4967 | 19.13 | 691 | 2.66 | 799 | 3.08 | 216 | 0.83 | 4 | 0.02 | 28.77 | 1.49 |
| 10 Random | PCCB, CLCN3, DOC2A, CUL9, SF3B1, FURIN, GATAD2A, PLCL1, INO80E, TMEM219 | 5265 | 20.28 | 6057 | 23.33 | 2581 | 9.94 | 1274 | 4.91 | 97 | 0.37 | 36.35 | 3.66 |
| All 15 | SNCA, SNAP91, CLCN3, DOC2A, PLCL1, THOC7, GATAD2A, ZNF823, INO80E, SF3B1, FURIN, PCCB, CALN1, CUL9, TMEM219 | 6755 | 26.02 | 8761 | 33.75 | 4988 | 19.21 | 3165 | 12.19 | 1030 | 3.97 | 42.58 | 7.89 |

### SUPPLEMENTAL BOXES & FIGURES

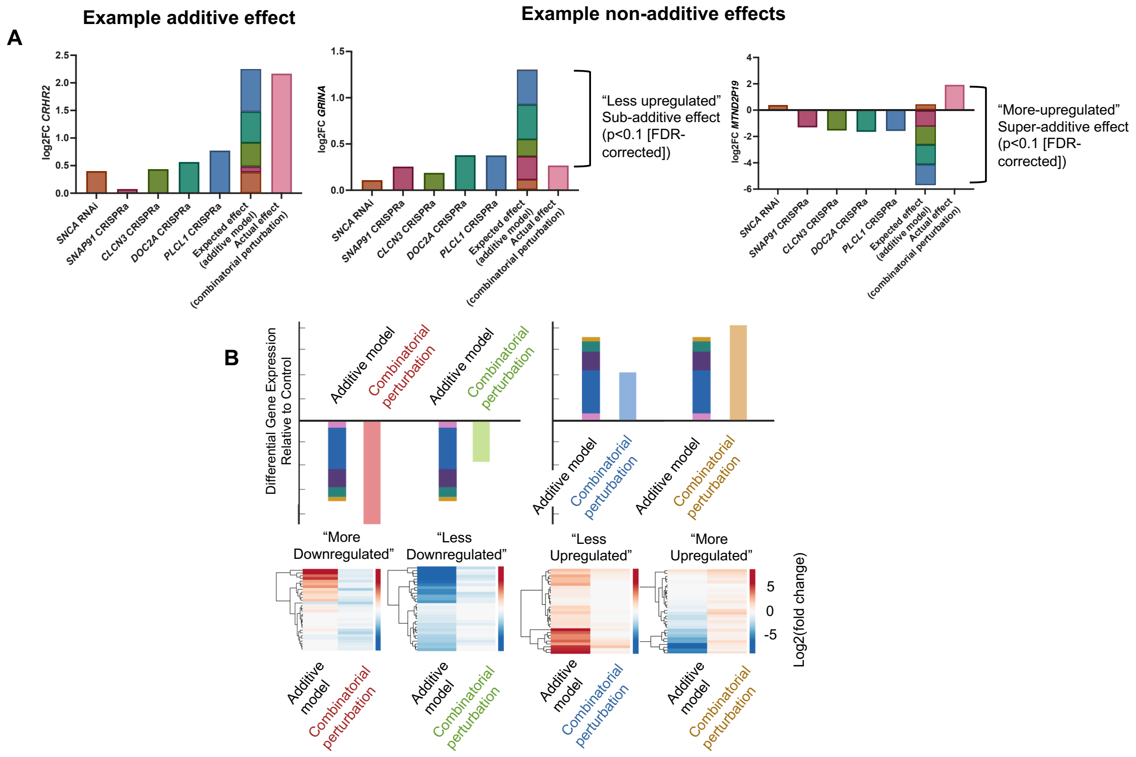


**Box 1.** ***Summary of additive and non-additive effects of joint eGene perturbation on transcription, related to Figure 4, Supplemental Figures 21-23.***

**A.** Example of genes demonstrating additive (*CRHR2*) and non-additive (*GRINA, MTNDP19*) effects following joint perturbation of 5 synaptic eGenes. The summed individual impact of eGene perturbations on *CRHR2* expression did not significantly differ from the impact of joint eGene perturbation, while the impact of joint eGene perturbation on *GRINA* expression was significantly reduced relative to the summed impact of individual eGene perturbations, suggesting saturation or redundancy of individual effects. In contrast to this, the impact of joint eGene perturbation on *MTNDP19* expression was significantly reversed relative to the summed impact of individual eGene perturbations, suggesting antagonistic summation of individual effects. **B.** Example figure showing different categories of non-additive effect found when comparing the impact of combinatorial perturbation relative to that in the predicted additive model. The top row shows exemplar effects on gene expression in the predicted additive model and in the joint perturbation condition. The bottom row shows exemplar heatmaps of differential gene expression for both conditions in each category. "More downregulated" and "more upregulated" categories show more dramatic or opposing effects on gene expression relative to the additive model, while "less downregulated" and "less upregulated" categories show smaller impacts on transcription in the joint perturbation condition when compared to the additive model. In the current study, we highlight those non-additive effects resulting in smaller changes on transcription than the summed individual effects predicted in the additive model ("less downregulated" and "less upregulated" categories, collectively termed “sub-additive” effects).

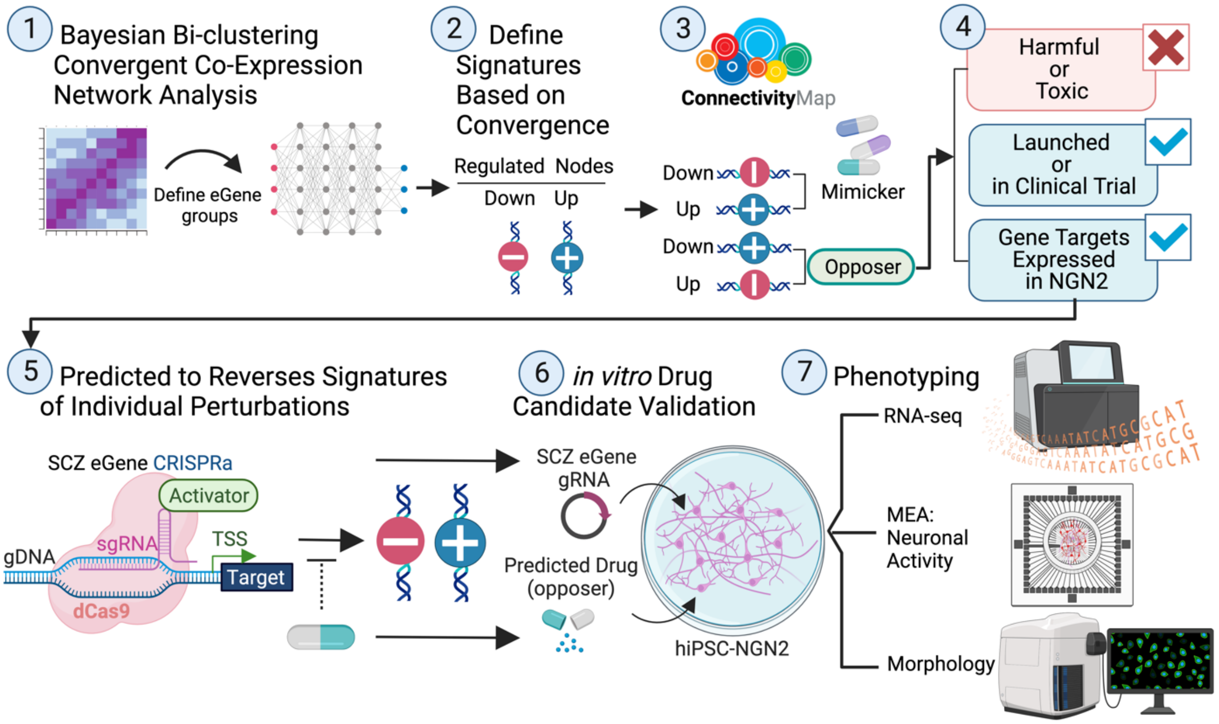


***Box 2. In vitro validation of in silico drug predictions, related to Figure 6, Supplemental Figures 28-30.***

Workflow of drug prioritization using LINCS drug database. (1) Bayesian bi-clustering performed across all 10 eGene perturbations identifies gene expression shared across all perturbations. (2) Un-supervised target convergent network reconstruction of the bi-clustering results resolves network relationships of convergent genes. (3) Top 150 up-regulated and down-regulated nodes identified by TCN reconstruction were used to define a network signature. (4) Network signatures were input into the Broads Institute's Connectivity Map query tool to predict drugs and perturbagens that mimic or reverse the target network signature. (5) Drugs that reversed the convergent network signature where filtered based on CS scores less than -1.00 in both neuronal (NEU) and neural progenitor cell (NPC) culture. Top drugs were filtered based on repurposing potential (Launched or in clinical trial), neurotoxicity and negative side effects, and expression of target genes in hiPSC-derived *NGN2*-neurons. (6) Following drug prioritization, their capacity to reverse individual perturbation signatures (based on the top up -and down-regulated genes) was assessed and eGene perturbations most likely to be reversed by the prioritized drugs were used in validation experiments *in vitro*. (7) CRISPR activation experiments of prioritized (based on CS score and significance in NEU and NPC culture) were carried out in iGLUT neurons treated with matched drug reversers for 48hrs. (8) Following CRISPRa and drug treatment, phenotypic assays (i.e. multi-electrode array (MEA) for neuronal activity; High-resolution microscopy for morphological measurements) to assess the impact of (i) perturbation, (ii) drug treatment, and (ii) perturbation by drug treatment on morphology, activity, and downstream gene expression of glutamatergic neurons were performed. Created with BioRender.com.

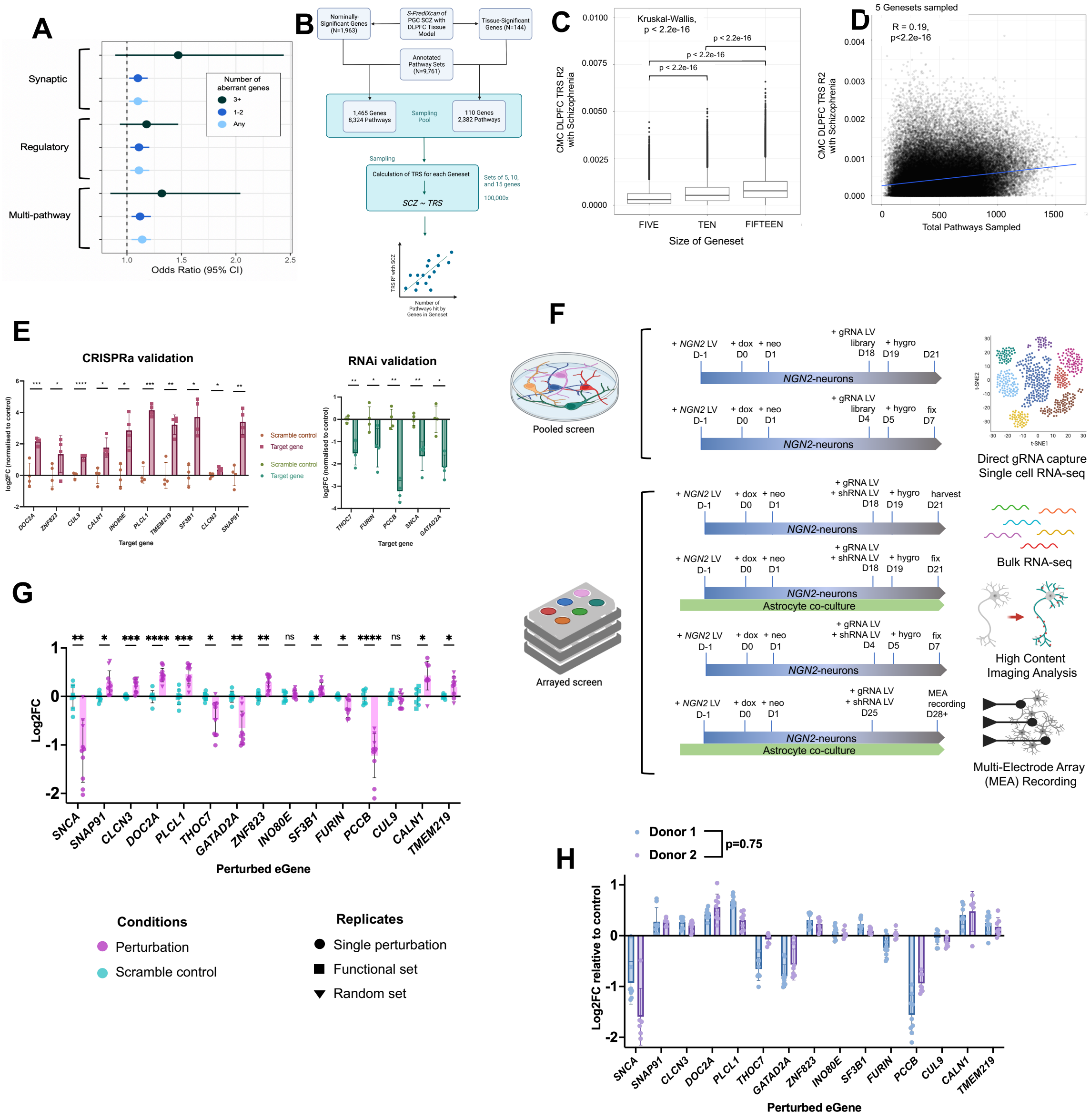

***Supplemental Figure 1. Prioritization and manipulation of synaptic, regulatory, and multi-function brain eGenes regulated by SCZ, related to Figure 1.***

**A**. Predicted GREX levels in dorsolateral prefrontal cortex (DLPFC) calculated for the fifteen eGenes in a Swedish SCZ cohort. Aberrant expression of eGenes is predicted to impact SCZ case-control status in a dose-dependent manner. Bars indicated the 95% confidence intervals for each odds ratio value. **B**. Schematic of random sampling of SCZ gene-sets for TRS. Random sets of 5, 10, and 15 genes were sampled from a pool of either nominally-significant, or tissue-specific-significant SCZ eGenes. TRS were calculated for each gene set, and pathways annotated for the genes included in each randomly sampled gene set. **C**. Boxplot demonstrating increasing TRS $R^2$ with SCZ with increasing number of genes sampled. **D**. Correlation of the number of pathways hit by randomly sampled sets of 5 nominally-significant SCZ genes with the geneset TRS $R^2$

with SCZ. **E**. QPCR validation of CRISPR activation and RNA interference in perturbing target eGenes in D7 hiPSC-NPC derived iGLUTs. 3-5 gRNA or shRNA vectors were tested per target eGene; successful vectors used for subsequent experiments shown in panel. One way ANOVA with posthoc Dunnett's multiple comparisons test, * = $p<0.05$; ** = $p<0.01$; *** = $p<0.001$; **** = $p<0.0001$. **F**. Schematic showing iGLUT induction, differentiation and perturbation timelines for each phenotyping method. For pooled screens: direct gRNA capture and single cell RNA-seq using independent CRISPRa libraries. For arrayed screens: bulk RNA-seq; synapse detection and neurite tracing using high content imaging; and MEA recording. All eGene perturbations were initiated 3 days prior to harvesting, fixing or recording glutamatergic cultures. A minimum of two independent experiments were performed on two healthy donor lines for each phenotyping method. **G.** CRISPR activation and RNA interference of target eGenes in D21 hiPSC-NPC derived iGLUTs, bulk RNA-seq samples. 13/15 eGenes showed significant perturbation in bulk RNA-seq data after correcting for multiple comparisons. Multiple t tests with Holm-Sidak correction for multiple comparisons. * = $p<0.05$; ** = $p<0.01$; *** = $p<0.001$; **** = $p<0.0001$. **H.** Comparison of perturbation efficacy across donors in D21 hiPSC-NPC derived iGLUT bulk RNA-seq data. Donor status did not significantly impact the degree of perturbation of target eGenes. Paired two-tailed test, $p=0.75$.

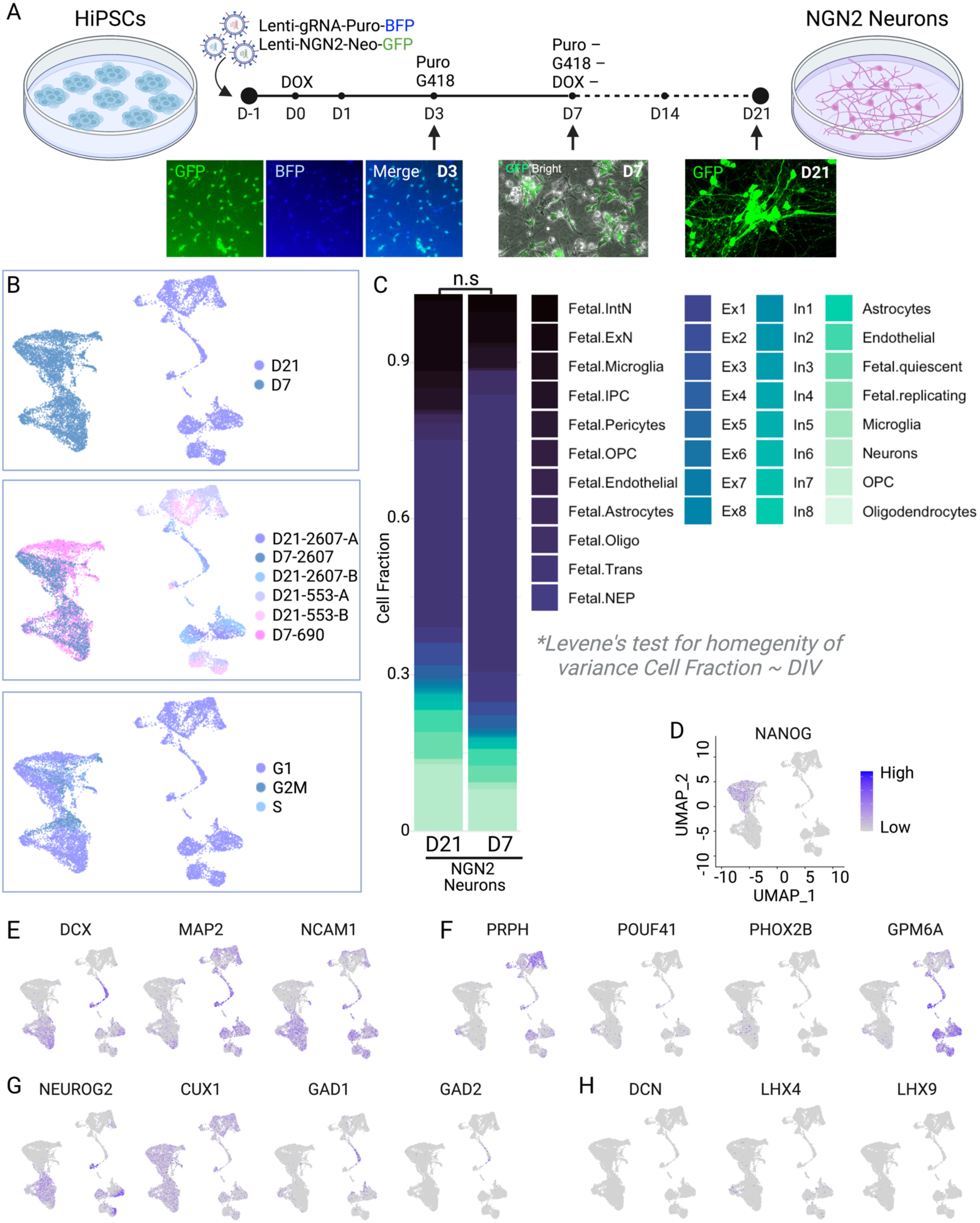

A
HiPSCs
Lenti-gRNA-Puro-BFP
Lenti-NGN2-Neo-GFP
DOX
Puro
G418
Puro –
G418 –
DOX –
D-1
D0
D1
D3
D7
D14
D21
NGN2 Neurons
GFP
BFP
Merge
D3
GFP Bright
D7
GFP
D21
B
D21
D7
D21-2607-A
D7-2607
D21-2607-B
D21-553-A
D21-553-B
D7-690
G1
G2M
S
C
n.s
Cell Fraction
0.9
0.6
0.3
0
D21
D7
NGN2
Neurons
Fetal.IntN
Fetal.ExN
Fetal.Microglia
Fetal.IPC
Fetal.Pericytes
Fetal.OPC
Fetal.Endothelial
Fetal.Astrocytes
Fetal.Oligo
Fetal.Trans
Fetal.NEP
Ex1
Ex2
Ex3
Ex4
Ex5
Ex6
Ex7
Ex8
In1
In2
In3
In4
In5
In6
In7
In8
Astrocytes
Endothelial
Fetal.quiescent
Fetal.replicating
Microglia
Neurons
OPC
Oligodendrocytes
*Levene's test for homegenity of variance Cell Fraction ~ DIV
D
NANOG
UMAP_2
UMAP_1
High
Low
E
DCX
MAP2
NCAM1
F
PRPH
POUF41
PHOX2B
GPM6A
G
NEUROG2
CUX1
GAD1
GAD2
H
DCN
LHX4
LHX9

***Supplementary Figure 2. Degree of variance in NGN2-induction is not significantly different between Day 7 and Day 21 iGLUTs.***

**A.** Diagram representing the differentiation time-course for both ECCITE-seq experiments from hiPSC induction at D0 to harvesting at D7 and D21, respectively. Fluorescence of GFP (*NGN2*-expression) and BFP (CRISPR gRNAs) demonstrate successful induction and infection with gRNAs. **B.** UMAP plots based on scRNA expression for D7 and D21 iGLUTs, color-coded by **(i)** day, **(ii)** donor and batch, **(iii)** cell-cycle phase. **C.** Bar plots of the average imputed cell fractions for gRNA-Scramble control cells across DIV and screening experiments with the proportion of each cell color-coded. While there are differences in the cell fraction represented by each cell type between experiments, a Levene's test for the homogeneity of variance (Cell Fraction ~ DIV; DF=1, F-value=0, Pr(>F)=1) was not significant and a Kruskal-Wallis test to compare distributions of cell-type fractions was nominally significant (Cell Fraction ~ DIV ; Kruskal-Wallis $X^2$=3.05, df=1, p-value=0.08062). **D-G.** Feature plots represent the expression of key cell-type markers for pluripotency seen in D0-D1 neurons (**D**), for general neuronal markers seen in D5-D28 iGLUTs **(E)**, for PNS and CNS neurons and neural projects **(F)**, for *NGN2*/glutamatergic specific markers **(G)**, and off-target (non-glutamatergic) neurons **(H)** overlaid atop of the UMAP plots for each Day/experiment. Expression of sub-type specific markers in certain clusters indicates cell-type heterogeneity in cell marker expression, with D7 neurons having unsurprisingly greater expression of NANOG than D21 neurons and D21 iGLUTs demonstrating some subtype heterogeneity (specifically PRPH a marker of PNS neurons). Neurons at both time points did not show expression of the off-target markers DCN, LHX4, and LHX9.

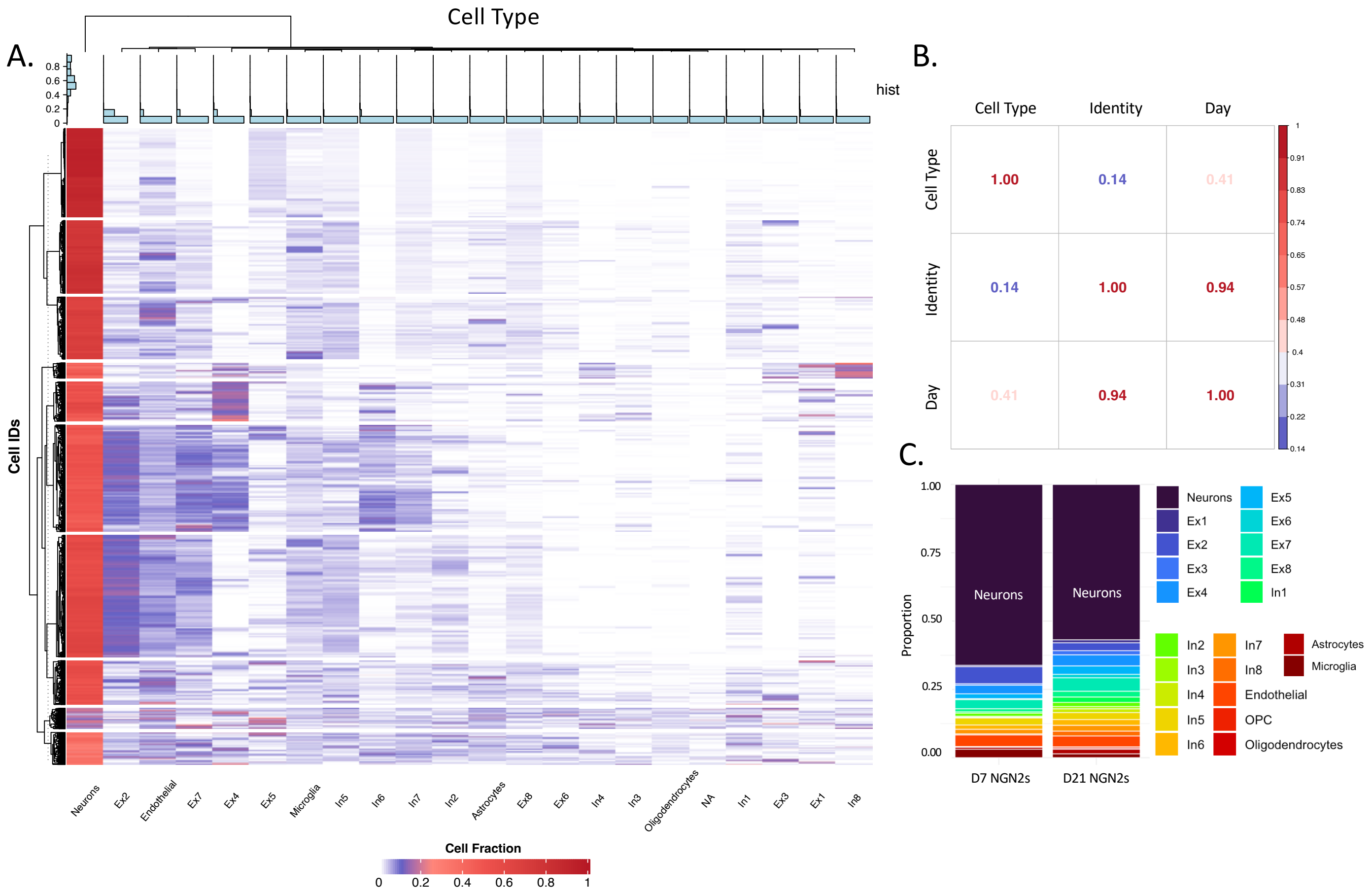


***Supplementary Figure 3. Degree of heterogeneity of iGLUTs is not significantly correlated with perturbation identity.***

**A.** Heatmap of adult cell-type fraction estimates across all cells in each experiment shows that transcriptomic signatures most strongly match neuronal cells followed by type-3 Excitatory neurons. **B.** There was no significant association between cell-type fraction estimates and gRNA identity (Pearson's correlation coefficient=0.14), with a nominal association between cell-type fraction and day (Pearson's correlation coefficient =0.41). **C.** Average cell-type fraction estimates in adult brain cells for D7 iGLUTs and D21 iGLUTs. Dox induced iGLUTs are transcriptomically most like neurons and excitatory neuron sub-types as expected.

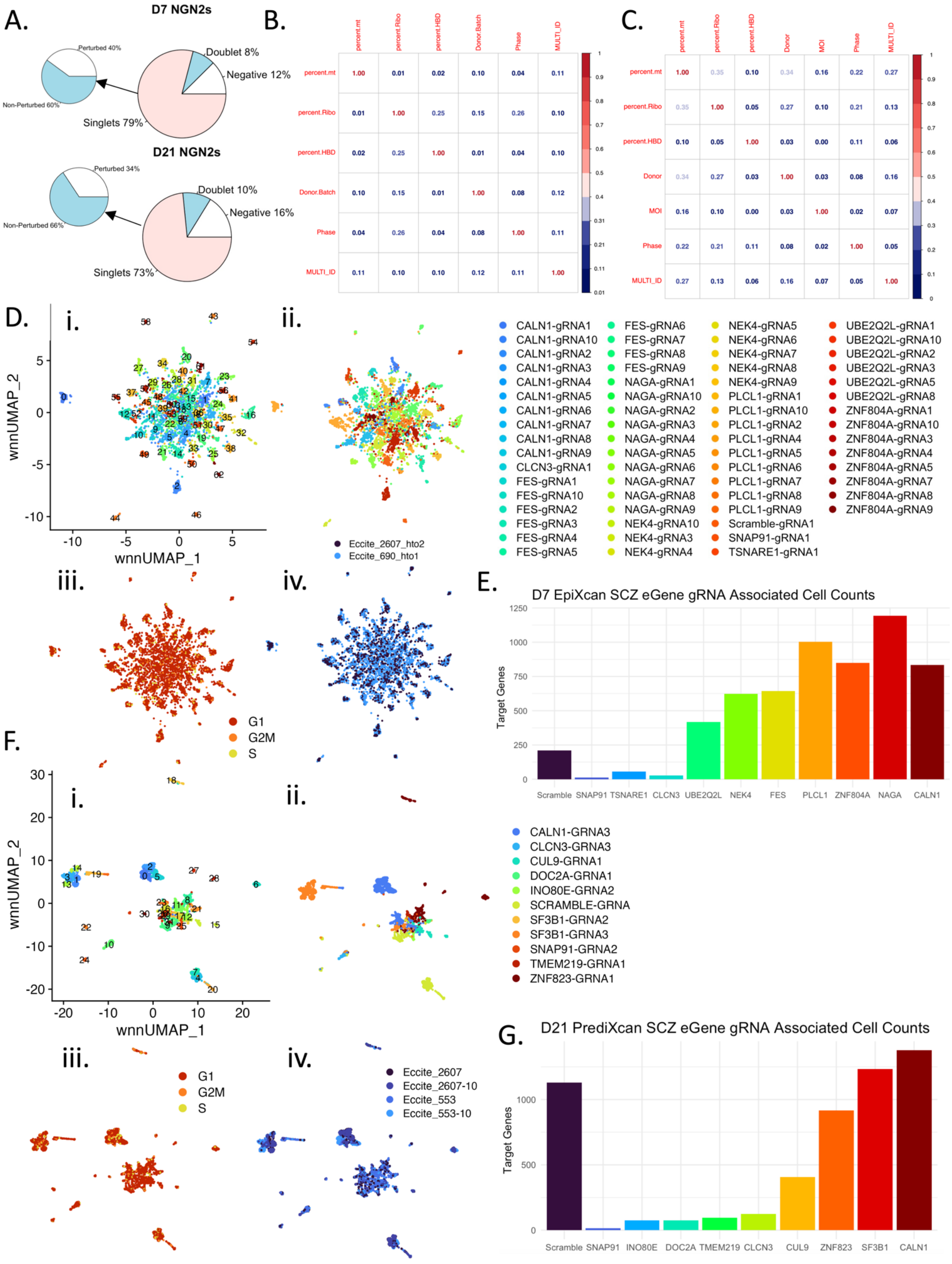
A.
D7 NGN2s
Perturbed 40%
Non-Perturbed 60%
Doublet 8%
Negative 12%
Singlets 79%
D21 NGN2s
Perturbed 34%
Non-Perturbed 66%
Doublet 10%
Negative 16%
Singlets 73%
B.
percent.mt percent.Ribo percent.HBD Donor.Batch Phase MULTI_ID
percent.mt 1.00 0.01 0.02 0.10 0.04 0.11
percent.Ribo 0.01 1.00 0.25 0.15 0.26 0.10
percent.HBD 0.02 0.25 1.00 0.01 0.04 0.10
Donor.Batch 0.10 0.15 0.01 1.00 0.08 0.12
Phase 0.04 0.26 0.04 0.08 1.00 0.11
MULTI_ID 0.11 0.10 0.10 0.12 0.11 1.00
C.
percent.mt percent.Ribo percent.HBD Donor MOI Phase MULTI_ID
percent.mt 1.00 0.35 0.10 0.34 0.16 0.22 0.27
percent.Ribo 0.35 1.00 0.05 0.27 0.10 0.21 0.13
percent.HBD 0.10 0.05 1.00 0.03 0.00 0.11 0.06
Donor 0.34 0.27 0.03 1.00 0.03 0.08 0.16
MOI 0.16 0.10 0.00 0.03 1.00 0.02 0.07
Phase 0.22 0.21 0.11 0.08 0.02 1.00 0.05
MULTI_ID 0.27 0.13 0.06 0.16 0.07 0.05 1.00
D. i. ii. iii. iv.
wnnUMAP_1
wnnUMAP_2
Eccite_2607_hto2
Eccite_690_hto1
G1
G2M
S
CALN1-gRNA1, CALN1-gRNA10, CALN1-gRNA2, CALN1-gRNA3, CALN1-gRNA4, CALN1-gRNA5, CALN1-gRNA6, CALN1-gRNA7, CALN1-gRNA8, CALN1-gRNA9, CLCN3-gRNA1, FES-gRNA1, FES-gRNA10, FES-gRNA2, FES-gRNA3, FES-gRNA4, FES-gRNA5
FES-gRNA6, FES-gRNA7, FES-gRNA8, FES-gRNA9, NAGA-gRNA1, NAGA-gRNA10, NAGA-gRNA2, NAGA-gRNA3, NAGA-gRNA4, NAGA-gRNA5, NAGA-gRNA6, NAGA-gRNA7, NAGA-gRNA8, NAGA-gRNA9, NEK4-gRNA10, NEK4-gRNA3, NEK4-gRNA4
NEK4-gRNA5, NEK4-gRNA6, NEK4-gRNA7, NEK4-gRNA8, NEK4-gRNA9, PLCL1-gRNA1, PLCL1-gRNA10, PLCL1-gRNA2, PLCL1-gRNA4, PLCL1-gRNA5, PLCL1-gRNA6, PLCL1-gRNA7, PLCL1-gRNA8, PLCL1-gRNA9, Scramble-gRNA1, SNAP91-gRNA1, TSNARE1-gRNA1
UBE2Q2L-gRNA1, UBE2Q2L-gRNA10, UBE2Q2L-gRNA2, UBE2Q2L-gRNA3, UBE2Q2L-gRNA5, UBE2Q2L-gRNA8, ZNF804A-gRNA1, ZNF804A-gRNA10, ZNF804A-gRNA3, ZNF804A-gRNA4, ZNF804A-gRNA5, ZNF804A-gRNA7, ZNF804A-gRNA8, ZNF804A-gRNA9
E.
D7 EpiXcan SCZ eGene gRNA Associated Cell Counts
Target Genes
Scramble SNAP91 TSNARE1 CLCN3 UBE2Q2L NEK4 FES PLCL1 ZNF804A NAGA CALN1
F. i. ii. iii. iv.
wnnUMAP_1
wnnUMAP_2
CALN1-GRNA3
CLCN3-GRNA3
CUL9-GRNA1
DOC2A-GRNA1
INO80E-GRNA2
SCRAMBLE-GRNA
SF3B1-GRNA2
SF3B1-GRNA3
SNAP91-GRNA2
TMEM219-GRNA1
ZNF823-GRNA1
G1
G2M
S
Eccite_2607
Eccite_2607-10
Eccite_553
Eccite_553-10
G.
D21 PrediXcan SCZ eGene gRNA Associated Cell Counts
Target Genes
Scramble SNAP91 INO80E DOC2A TMEM219 CLCN3 CUL9 ZNF823 SF3B1 CALN1

***Supplementary Figure 4. Weighted Nearest Neighbor (WNN) dimensional reduction identifies successfully perturbed cells.***

**A.** Pie chart of distribution of negatives, doublets, and singlets (further subset by perturbed for a single eGene and non-perturbed based on up-regulation of eGene compared to scramble controls) identified within each pooled CRISPR screen. **B-C.** Correlation coefficients of covariates corrected for in the Day 7 (B) and Day 21 (C) pooled CRIPSRa screens and gRNA identity (MULTI_ID). **D.** UMAP of D7 CRISPRa cells follow WNN analysis, clustered by (i) WNN cluster, (ii) gRNA identity, (iii) cell-cycle phase, and (iv.) donor and batch. **E.** Total number of cells per EpiXcan eGene in D7 CRISPRa screen. **F.** UMAP of D21 CRISPRa cells follow WNN analysis, clustered by (i) WNN cluster, (ii) gRNA identity, (iii) cell-cycle phase, and (iv.) donor and batch. **G.** Total number of cells per PrediXcan eGene in D21 CRISPRa screen.

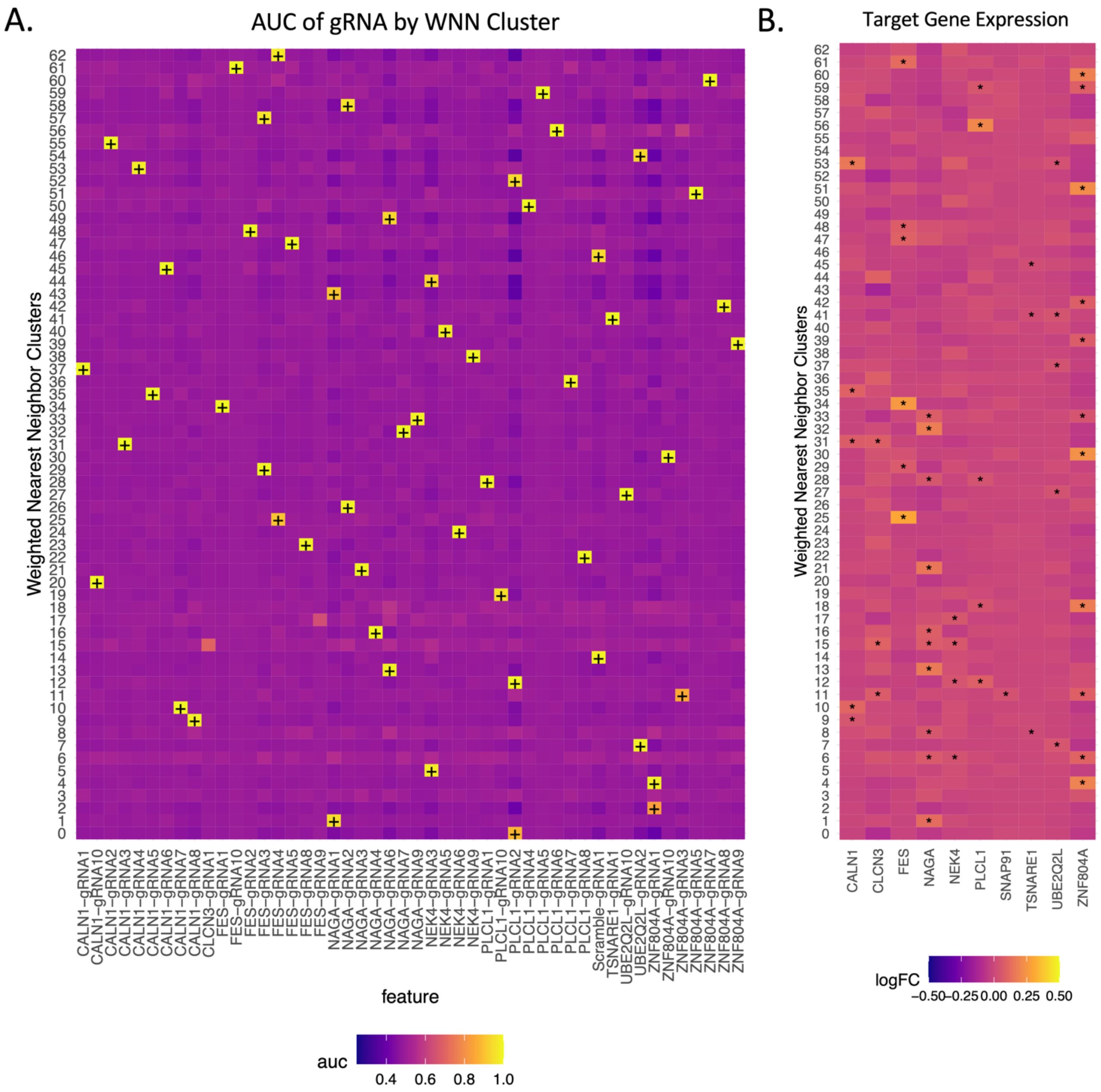


***Supplementary Figure 5. Multi-modal analysis allows for identification of successfully perturbed cells by dimensional reduction and weighted clustering in D7 iGLUTs.***

**A.** Wilcox-rank sum analysis of gRNA count matrices calculated an Area Under the Curve (AUC) statistic, which reflects the power of each gRNA to serve as a marker o the cell cluster (how well a specific gRNA defines the cell identity). Heatmap colors are coded by AUC with WNN clusters on the x-axis and gRNA identities on the y-axis. Black plus signs (+) represent AUC scores greater than 0.80. **B.** Heatmap of log fold-change (LogFC) of eGenes by WNN cluster. Wilcox rank sum test of RNA count matrix to identify successfully upregulated gene targets by WNN cluster (as compared to all other clusters). Black stars (*) represent significantly up-regulated gene expression.

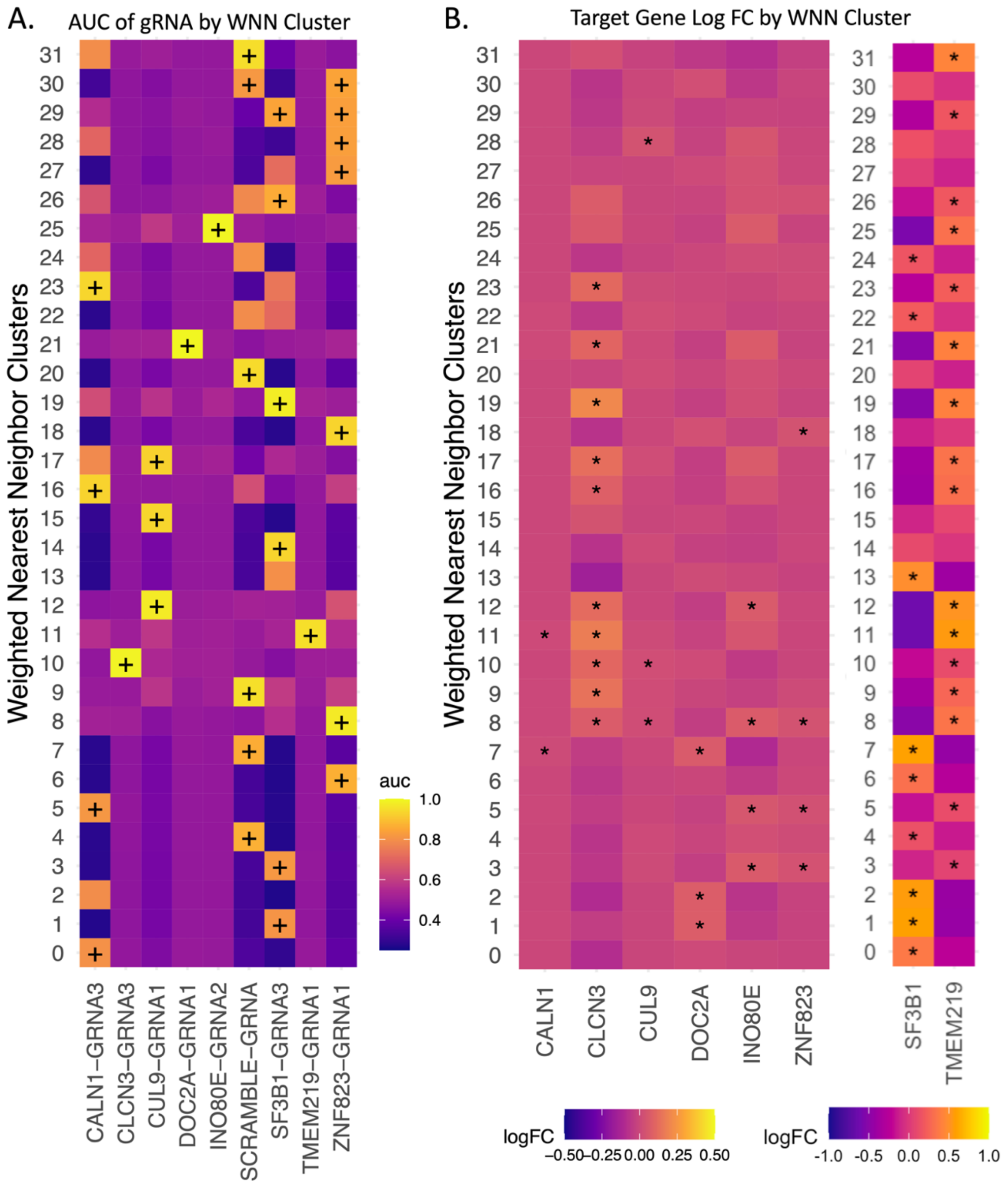


***Supplementary Figure 6. Multi-modal analysis allows for identification of successfully perturbed cells by dimensional reduction and weighted clustering in D21 iGLUTs.***

**A.** Heatmap of AUC based on gRNA count matrices with WNN clusters on the x-axis and gRNA identities on the y-axis. Black plus signs (+) represent AUC scores greater than 0.80. **B.** Heatmaps of logFC based on Wilcox-rank sum analysis of gene expression within each WNN cluster compared to all other clusters. Black stars (*) represent significantly up-regulated gene expression.

A.

Number of Successfully Perturbed Cells
by gRNA Targets

| Variable | N | Percent |
|---|---|---|
| Identity | 2355 | |
| ... CALN1-gRNA | 435 | 18.5% |
| ... FES-gRNA | 361 | 15.3% |
| ... NAGA-gRNA | 552 | 23.4% |
| ... PLCL1-gRNA | 244 | 10.4% |
| ... Scramble-gRNA | 164 | 7% |
| ... UBE2Q2L-gRNA | 224 | 9.5% |
| ... ZNF804A-gRNA | 375 | 15.9% |

B.

Number of Successfully Perturbed Cells
by gRNA Targets

| Variable | N | Percent |
|---|---|---|
| Identity | 1623 | |
| ... CLCN3-gRNA | 123 | 7.6% |
| ... CUL9-gRNA | 55 | 3.4% |
| ... INO80E-gRNA | 73 | 4.5% |
| ... Scramble-gRNA | 421 | 25.9% |
| ... SF3B1-gRNA | 511 | 31.5% |
| ... TMEM219-gRNA | 93 | 5.7% |
| ... ZNF8043-gRNA | 347 | 21.4% |

C.

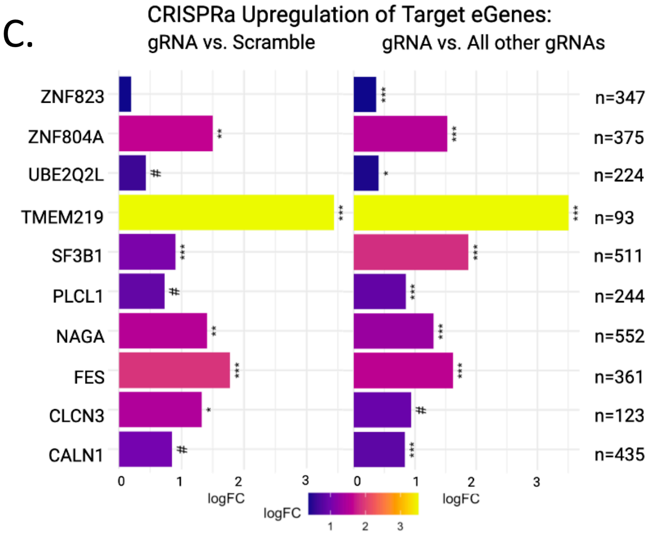


***Supplemental Figure 7. Successful CRISPR-activation of top SCZ eGenes compared to predicted effects of eQTL causal SNPs.***

**A-B.** Cell number per perturbation where gRNA AUC >=0.80 for a single target gene and the matched eGene was significantly upregulated compared to all other clusters, indicating successful perturbation for DIV 7 **(A)** and DIV 21 **(B)** CRISPRa screens. **C.** A one-way pairwise Wilcox Rank Sum test (AUC statistic on the x-axis) comparing perturbed cells with matched scramble controls found that 10 eGenes (n=93-552) out of the 12 resolvable targets across both experiments were significantly upregulated (unadjusted p-value **#** <= 0.05; adjusted p-value: * < 0.05, ** <= 0.01, ***<0.001).

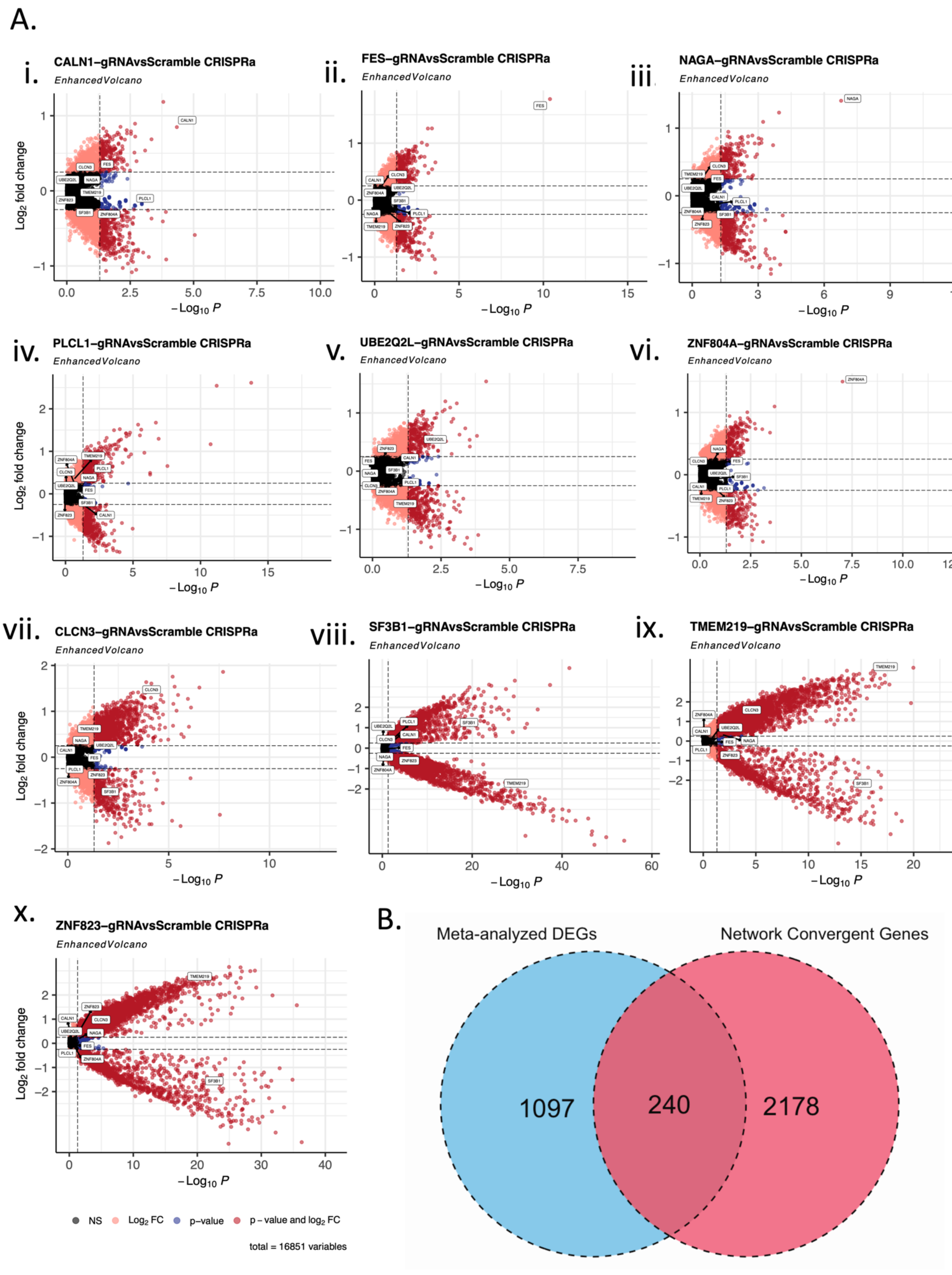

A.
i.
CALN1–gRNAvsScramble CRISPRa
EnhancedVolcano
ii.
FES–gRNAvsScramble CRISPRa
EnhancedVolcano
iii.
NAGA–gRNAvsScramble CRISPRa
EnhancedVolcano
iv.
PLCL1–gRNAvsScramble CRISPRa
EnhancedVolcano
v.
UBE2Q2L–gRNAvsScramble CRISPRa
EnhancedVolcano
vi.
ZNF804A–gRNAvsScramble CRISPRa
EnhancedVolcano
vii.
CLCN3–gRNAvsScramble CRISPRa
EnhancedVolcano
viii.
SF3B1–gRNAvsScramble CRISPRa
EnhancedVolcano
ix.
TMEM219–gRNAvsScramble CRISPRa
EnhancedVolcano
x.
ZNF823–gRNAvsScramble CRISPRa
EnhancedVolcano
$\log_2$ fold change
$-\log_{10} P$
NS
$\log_2$ FC
p – value
p – value and $\log_2$ FC
total = 16851 variables
B.
Meta-analyzed DEGs
Network Convergent Genes
1097
240
2178

***Supplemental Figure 8. Successful CRISPR-activation of top SCZ eGenes leads to significant changes in gene expression.***

**6-Target ECCITE-seq (D7 iGLUTs): A. (i-vi)** Volcano plots of differential gene expression (DGE) for each target perturbation compared to scramble controls.

**4-Target ECCITE-seq (D21 iGLUTs): A. (vi-x)** Volcano plots of differential gene expression (DGE) for each target perturbation compared to scramble controls. **B.** Overlap of mt-analyze DEGs across all ten targets and convergent genes identified by Bayesian bi-clustering and network reconstruction.

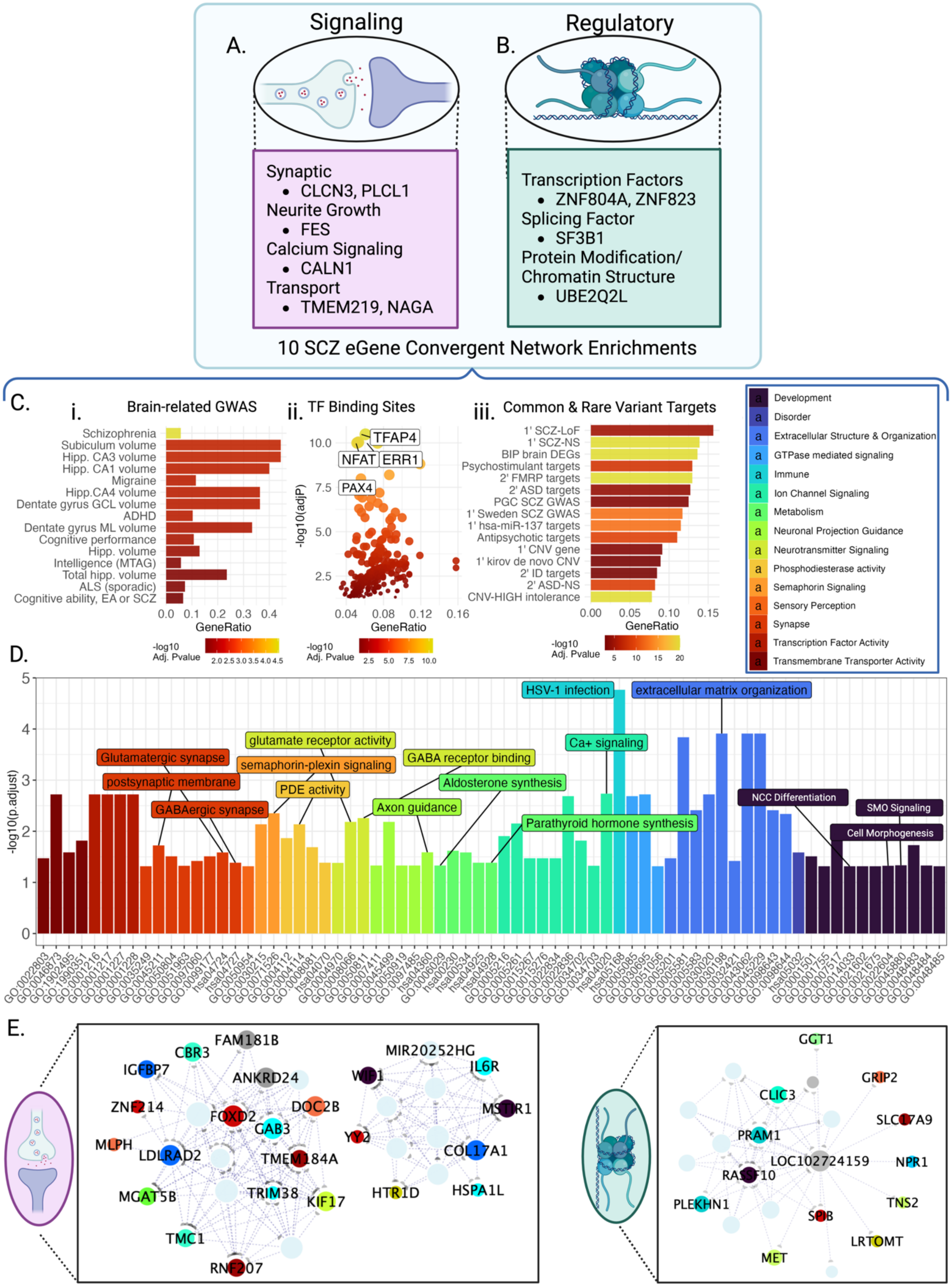


***Supplemental Figure 9. Downstream target-convergent networks identified by Bayesian bi-clustering resolve distinct networks enriched for SCZ common and***

***rare variant target genes and transcription factor binding motifs, related to Figure 2, SI Data 3***.

For the pooled screen, the resolved ten SCZ eGene could be separated based on general function as **(A)** signaling or (**B)** regulatory genes. **C.** Convergent networks resolved across the downstream transcriptomic impacts of all ten target perturbations identified 2418 convergent genes with enrichments for **(i)** brain-related GWAS genes, **(ii)** transcription factor binding sites of know SCZ-associated TFs (TFAP4, NFAT and ERR1), and **(iii)** common and rare variant target genes. **D.** Gene set enrichment analysis of convergent genes shared across all ten eGene perturbation were significantly enriched in multiple terms related to synaptic signaling and structure, neuronal projection and development, voltage-gated ion channel signaling, phosphodiesterase signaling, and semaphoring signaling. Full enrichment results are found in Supplemental Data 3. **E.** Unique convergent networks were resolved across signaling (left) and regulatory (right) SCZ Genes with node genes involved in synaptic signaling and immune response. Node color corresponds with general functional term found in **(D)**, with RNA genes colored in pale blue, while node size indicates node degree, and the line thickness represents weight of the connection (calculated as the percent duplication of the connection across bi-clustering runs). Created with BioRender.com

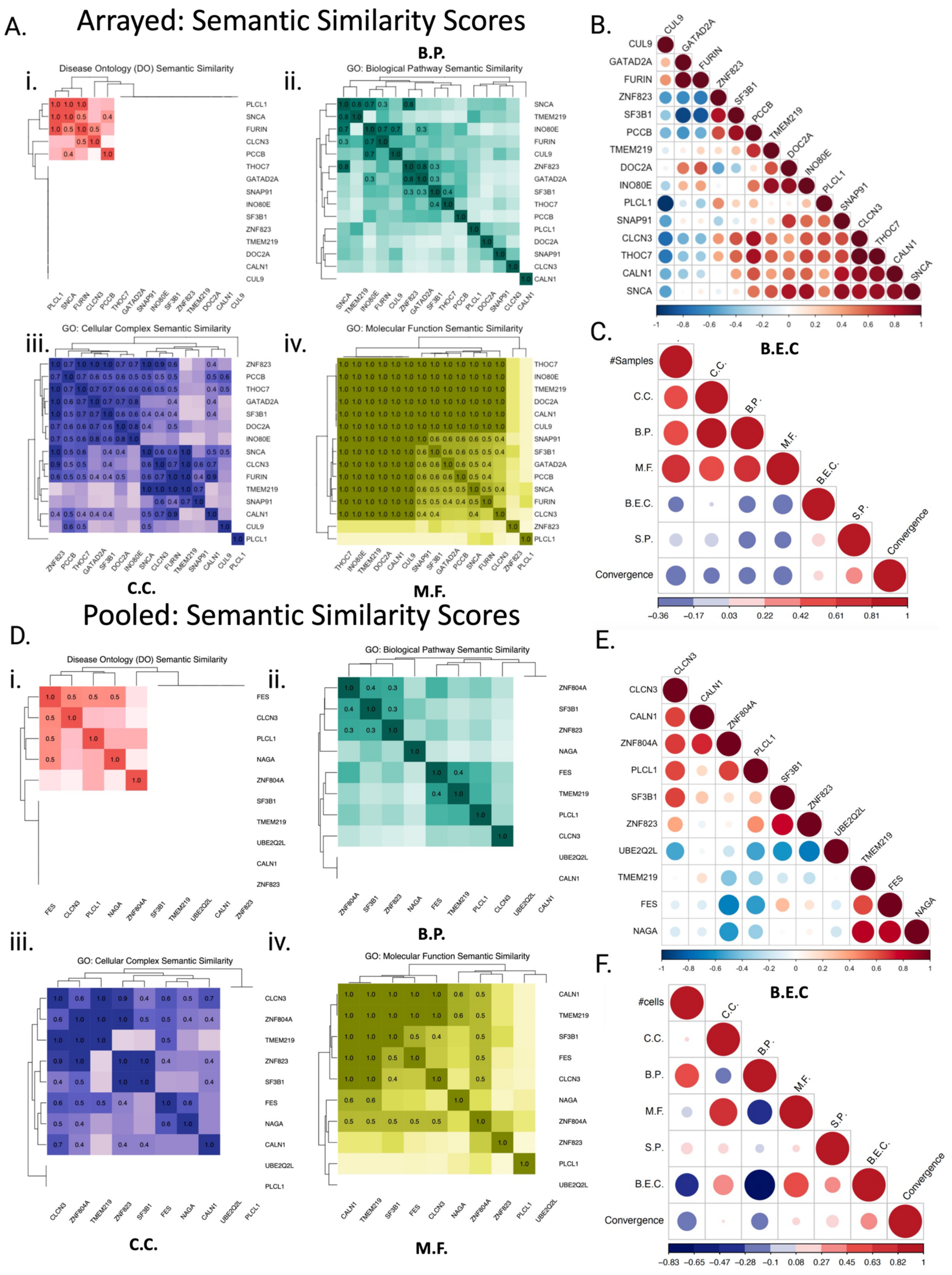
A.
Arrayed: Semantic Similarity Scores
i.
Disease Ontology (DO) Semantic Similarity
PLCL1
SNCA
FURIN
CLCN3
PCCB
THOC7
GATAD2A
SNAP91
INO80E
SF3B1
ZNF823
TMEM219
DOC2A
CALN1
CUL9
B.P.
ii.
GO: Biological Pathway Semantic Similarity
iii.
GO: Cellular Complex Semantic Similarity
C.C.
iv.
GO: Molecular Function Semantic Similarity
M.F.
B.
C.
#Samples
C.C.
B.P.
M.F.
B.E.C.
S.P.
Convergence
B.E.C
Pooled: Semantic Similarity Scores
D.
i.
Disease Ontology (DO) Semantic Similarity
FES
CLCN3
PLCL1
NAGA
ZNF804A
SF3B1
TMEM219
UBE2Q2L
CALN1
ZNF823
ii.
GO: Biological Pathway Semantic Similarity
B.P.
iii.
GO: Cellular Complex Semantic Similarity
C.C.
iv.
GO: Molecular Function Semantic Similarity
M.F.
E.
F.
#cells
B.E.C
-0.36 -0.17 0.03 0.22 0.42 0.61 0.81 1
-0.83 -0.65 -0.47 -0.28 -0.1 0.08 0.27 0.45 0.63 0.82 1

***Supplemental Figure 10. Functional scoring of perturbation sets, related to Figure 3, SI Figure 11.***

**A & D.** Heatmaps visualizing the degree of semantic similarity (0-1) between perturbed SCZ eGenes from the arrayed (**A**) or pooled (**D**) screens based on Disease Ontology (DO) and Gene Ontology (GO) Biological Pathway (B.P.), Cellular Component (C.C.), and Molecular Function (M.F.) respectively. **B & E.** Brain Expression Correlation (B.E.C.) represents the correlation strength (-1 to 1) of gene expression of perturbed SCZ eGenes from the arrayed (B) or pooled (**E**) screens in the postmortem CMC dorsolateral pre-frontal cortex (DLPFC). **C & F.** Correlations between sample size of the set, functional scores, and convergence degree demonstrate positive correlations between network convergence degree and DLPFC gene expression and signaling proportion of perturbed eGenes in both the arrayed (**C**) and pooled (**F**) screen.

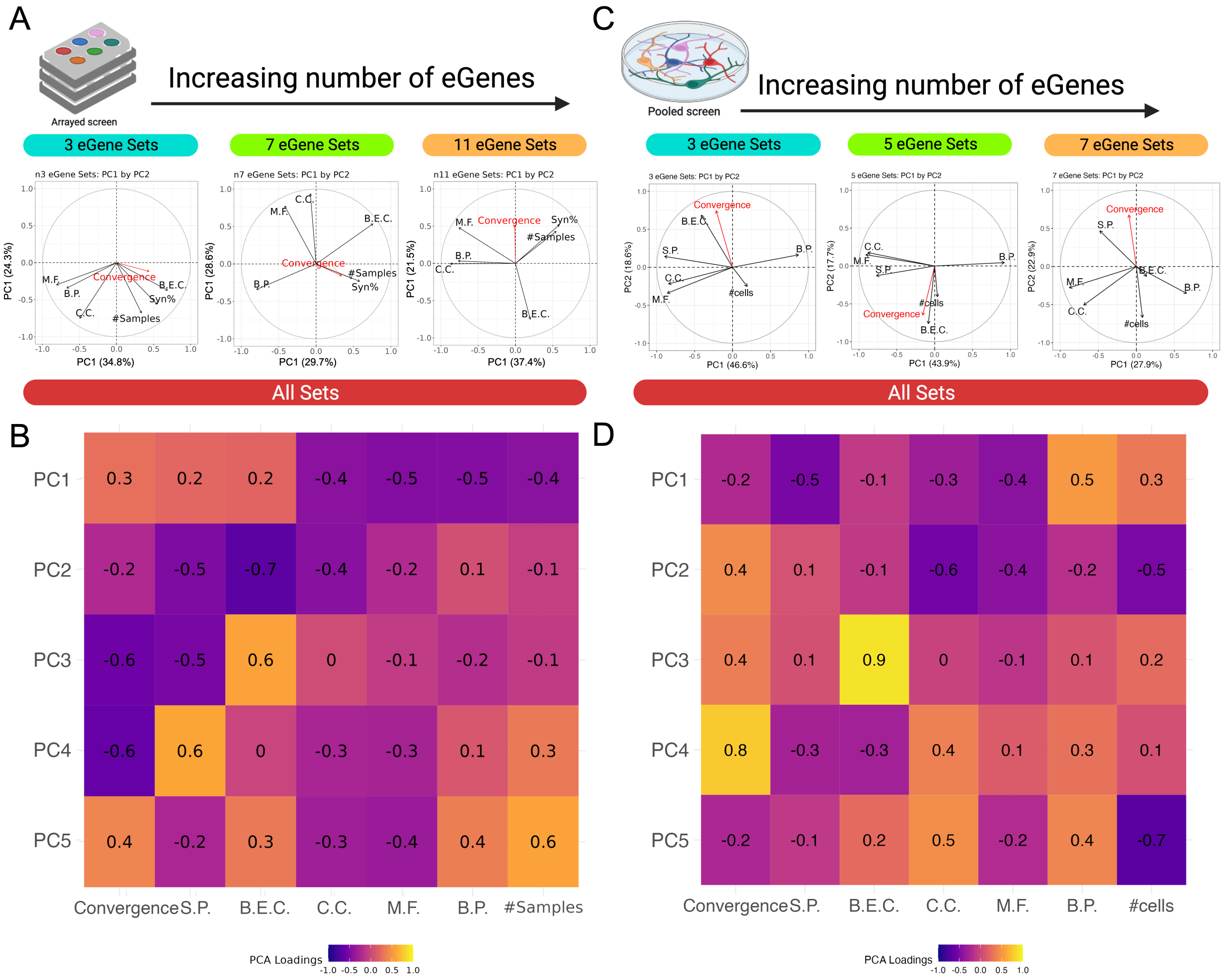


***Supplementary Figure 11. The degree of network convergence is influenced by functional similarity of target perturbations, related to Figure 3, SI fig 10***

*Principal Component Analysis across PCs demonstrate the degree of variance between convergent networks each functional score influences.* **Arrayed Experiment: A.** PCA biplots visualizing the loadings of PCs 1 and 2 for networks reconstructed across an increasing number of eGene perturbations from the arrayed experiment. The loadings are represented as labeled maroon lines that demonstrate how strongly each characteristic (convergence degree, sample size, functional score) influences the principal component **B.** Heatmap of PCA loadings of PCs 1-5 of each label when including all possible random sets. **Pooled Experiments: C.** PCA biplots visualizing the loadings of PCs 1 and 2 for networks reconstructed across an increasing number of eGene perturbations from the arrayed experiment. The loadings are represented as labeled maroon lines that demonstrate how strongly each characteristic (convergence degree, sample size, functional score) influences the principal component **D.** Heatmap of PCA loadings of PCs 1-5 of each label when including all possible random sets.

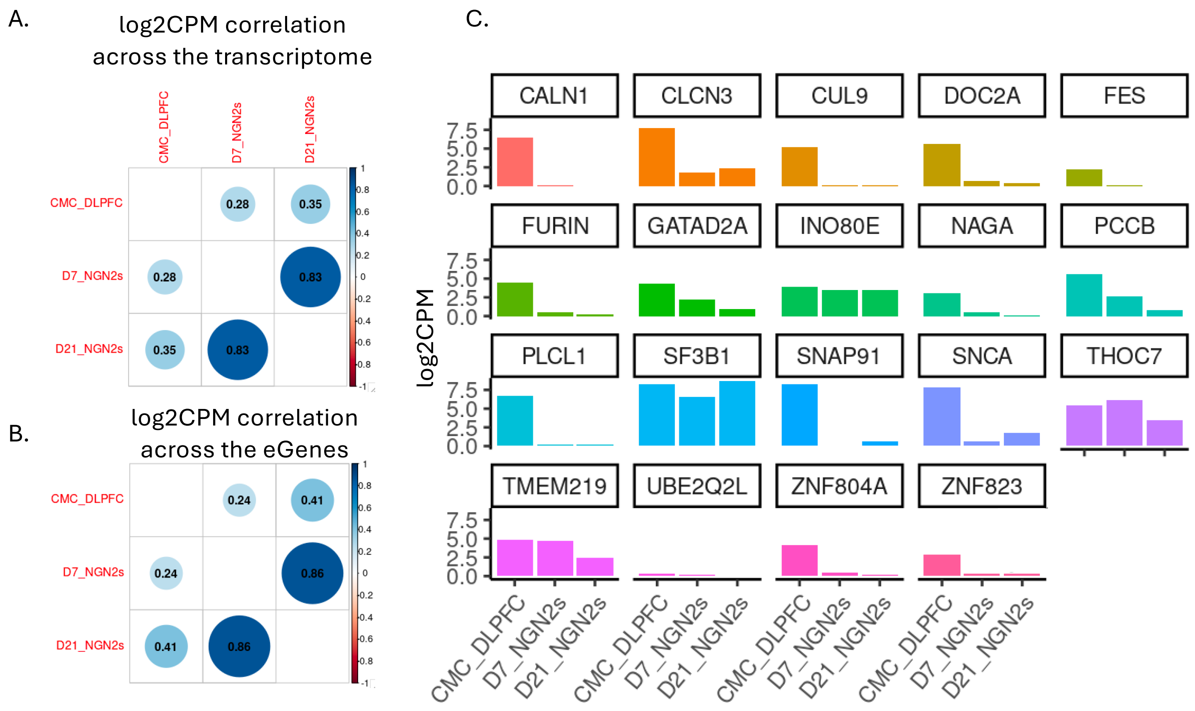


***Supplementary Figure 12**. Gene expression of D7 and D21 NGN2-neurons significantly correlate with gene expression of the postmortem adult DLPFC.* **A.** Correlation of log2CPM transcriptome-wide gene expression signatures across adult DLPFC bulk RNA-sequence from neurotypical donors (n=541) and NGN2 single-cell RNA-seq in scramble controls. **B.** Correlation of log2TPM gene expression signatures of CRIPSR-targeted SCZ eGenes between across adult DLPFC bulk RNA-sequence from neurotypical donors (n=541) and NGN2 single-cell RNA-seq in scramble controls. **C.** Average expression (log2CPM) of CRIPSR-targeted SCZ eGenes in the adult DLPFC (neurotypical donors, n=541) and NGN2 single-cell RNA-seq in scramble controls.

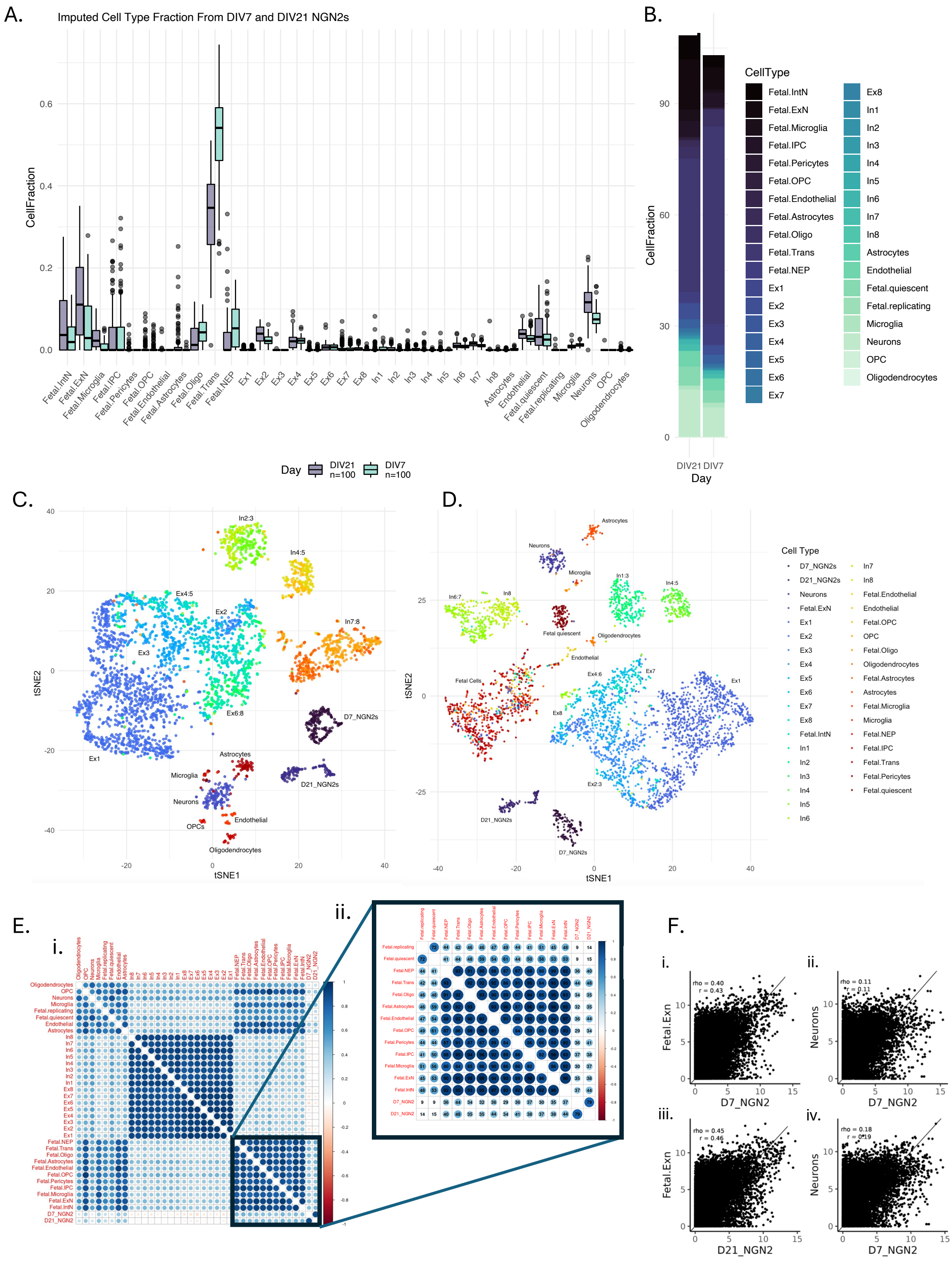

**Supplementary Figure 13. Gene expression of D7 and D21 NGN2-neurons significantly correlate with gene expression of adult cortical neurons and fetal trans and excitatory neurons. A-B.** Average cell-type fraction estimates of adult and fetal brain cell-types in the frontal cortex in D7 *NGN2*-neurons and D21 *NGN2*-neurons.

Dox induced *NGN2*-neurons are transcriptomically most like neurons and excitatory neuron sub-types. **C-D.** *Uniform Manifold Approximation and Projection (UMAP)* for Dimension Reduction of gene expression (TPM) from adult and fetal cortical cells and D7 and D21 NGN2 scramble controls used in this study. **E.** Correlation of log2TPM gene expression signatures across adult and fetal brain cell types and NGN2 scramble controls. **F.** Correlation of log2TPM gene expression signatures between Fetal excitatory neurons and forebrain neurons in the cortex with D7 and D21 NGN2s (Pearson's rho and Spearman's r correlation coefficients annotated in top left corner).

A.

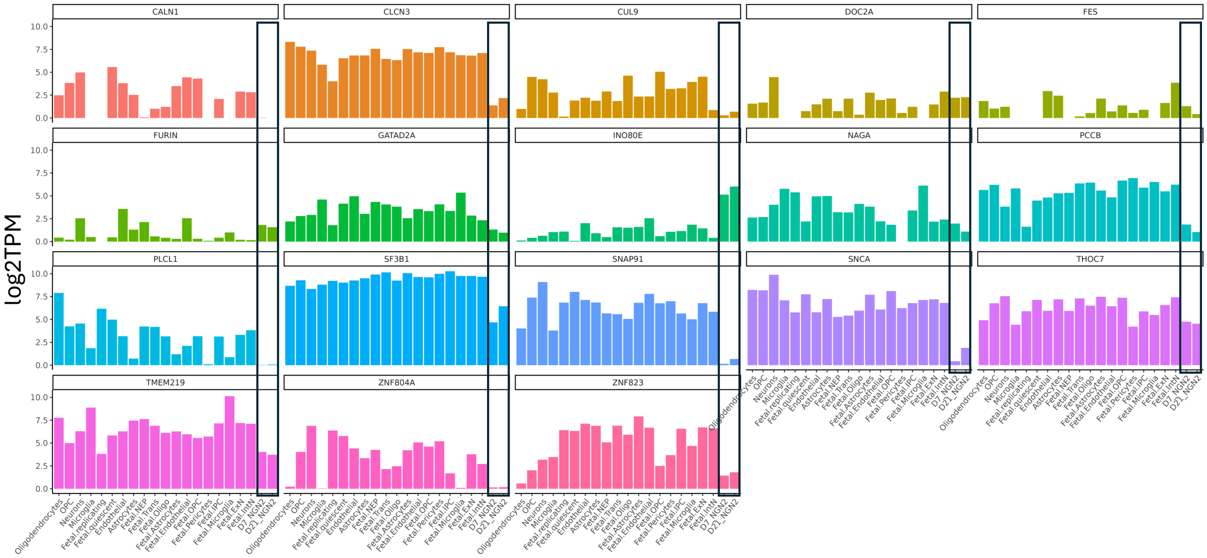


**Supplementary Figure 14**. ***Basal gene expression of SCZ eGenes perturbed in this study in D7 and D21 NGN2-neurons and fetal and adult cell-types in the prefrontal cortex***. **A.** Average expression (log2TPM) of CRIPSR-targeted SCZ eGenes in adult and fetal single-cell RNAseq from the frontal cortex and NGN2 single-cell RNA-seq in scramble controls.

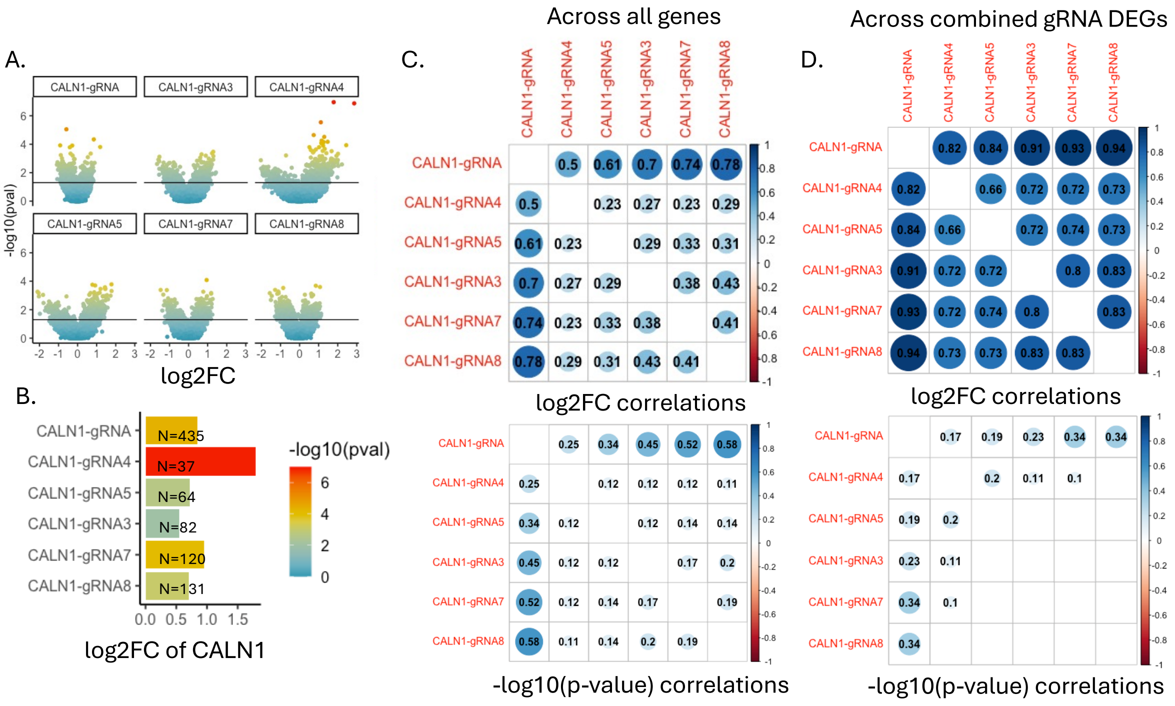


***Supplementary Figure 15. Concordance of differential gene expression patterns across individual gRNAs targeting CALN1.*** **A.** Volcano plots of differentially expressed genes when comparing all CALN1-gRNAs to Scramble-gRNA (CALN1-gRNA) and comparing each gRNA to scramble individually. **B.** Degree of *CALN1* upregulation (log2FC) across individual gRNAs and with gRNAs combined (CALN1-gRNA). N=number of cells containing each guide (or all guides combined). **C.** Correlation of log2FC and -log10(p-value) across the entire transcriptome between gRNAs individually and combined. **D.** Correlation of log2FC and -log10(p-value) across significantly differentially expressed genes between gRNAs individually and combined.

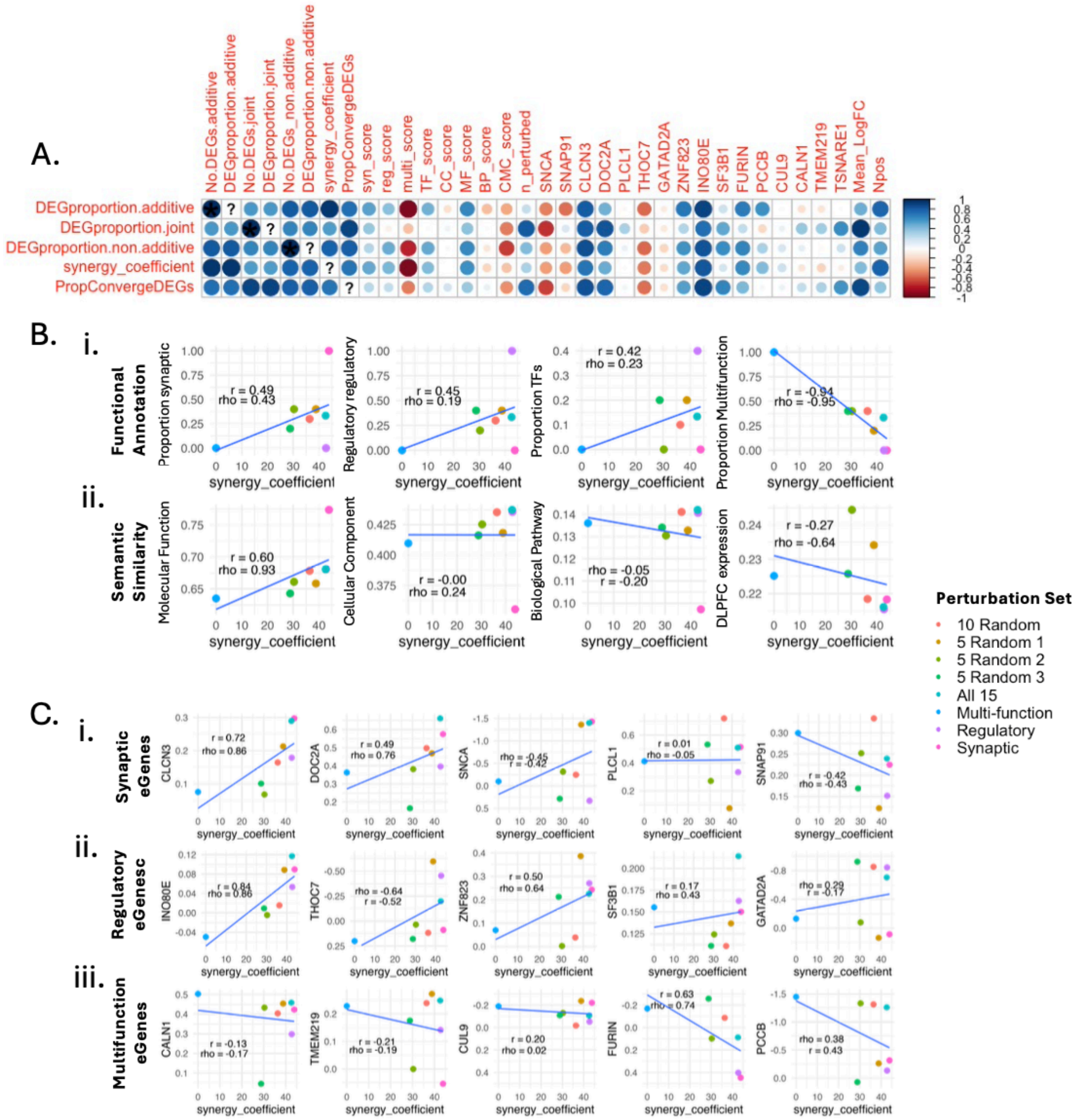


**Supplementary Figure 16**. **Similarity in molecular function and degree of perturbation of synaptic and regulatory genes drives non-additivity**. **A.** Pearson's correlation matrix of metrics of characterizing non-additivity in the join and combinatorial bulk-arrayed screens with functional similarity scores between eGenes and log2FC of the perturbed eGenes compared to scramble. **B.** Perturbation of eGenes with greater similarity in molecular function positively correlates with greater non-additivity **(i)** Scatter plots demonstrating the correlation between the proportion of synaptic, regulatory, or multifunctional genes in a combinatorial set (y-axis) and the synergy coefficient (x-axis). **(ii)** Scatter plots demonstrating the correlation between the semantic similarity scores by molecular function, cellular component, or biological function, or post-mortem brain expression correlations in a combinatorial set (y-axis) and the synergy coefficient (x-axis). **C.** The degree of perturbation of synaptic and regulatory eGenes, but not multifunction eGenes, correlates with greater non-additivity Scatter plots demonstrating the correlation between the degree of logFC of each eGene and synergy coefficients across all combinatorial sets. Pearson's rho and Spearman's r correlation coefficients annotated on each plot.

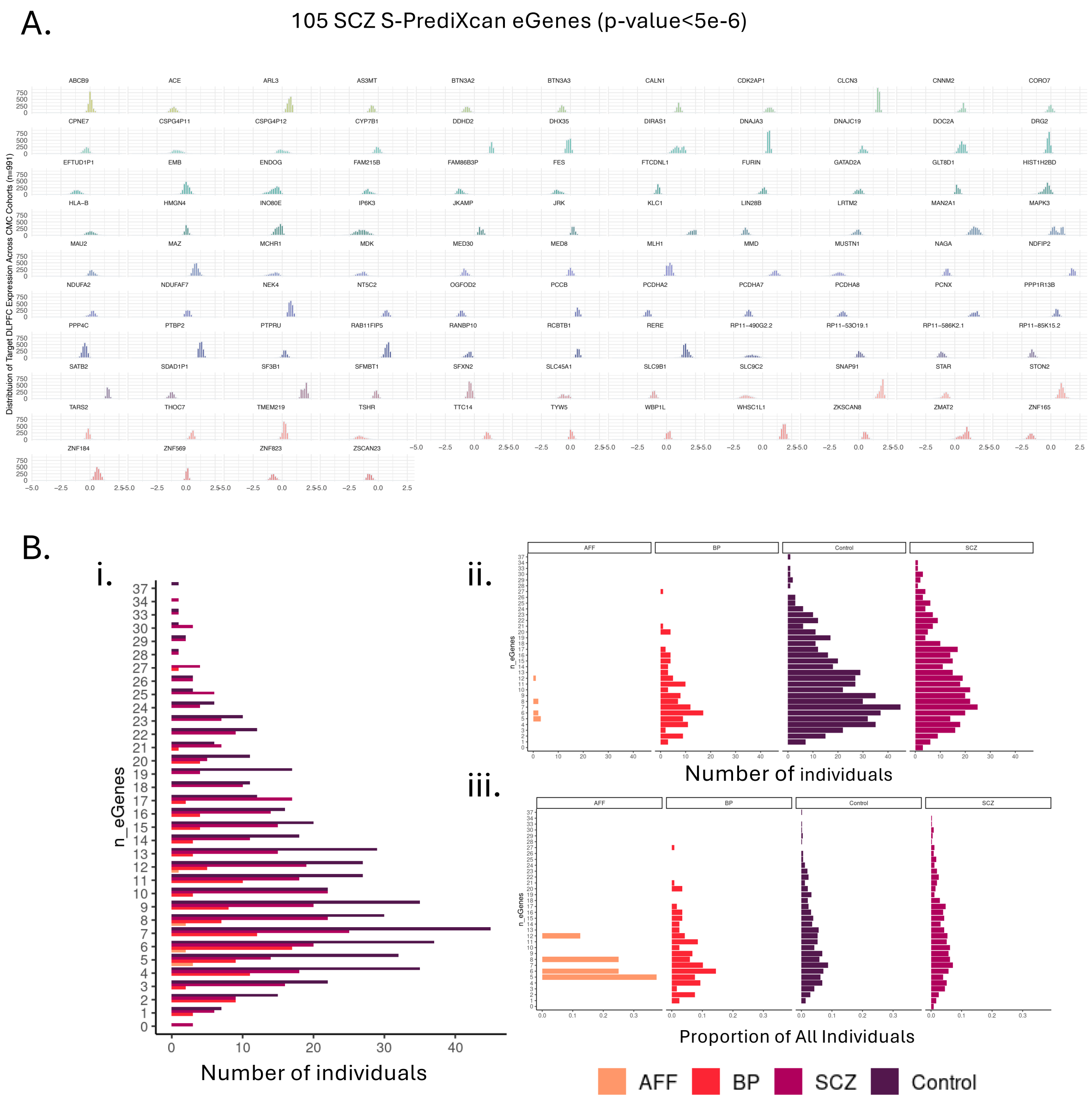


**Supplementary Figure 17*. Across ~100 SCZ risk genes, a median of 10 were classified as perturbed per individual.*** **A**. Z-scored expression levels of 105 SCZ S-PrediXcan eGenes in the Common Mind Consortiums adult post-mortem DLPFC. **B**. (i) Number of individuals by diagnosis and number of estimated eGene perturbations (eGene perturbation assigned based on expression levels in the top or bottom 10%). (ii) frequency separated by diagnosis (iii) absolute proportion separated by diagnosis.

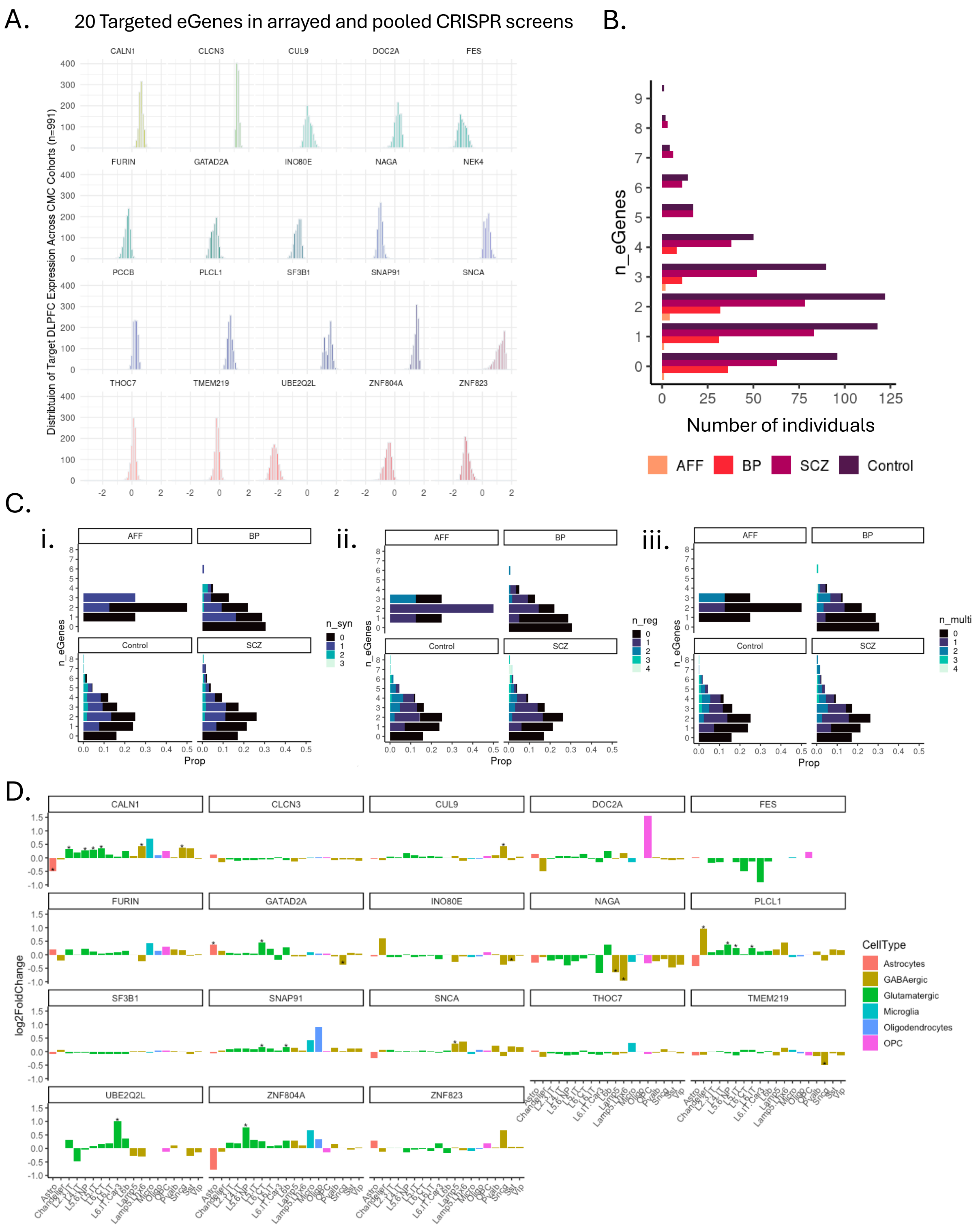


**Supplementary Figure 18. Across 20 SCZ risk genes targeted in the arrayed and pooled screens, an average of 2 and a maximum of 8 were classified as perturbed per individual**. **A.** Z-scored expression levels of the 20 SCZ eGenes perturbed in CRISPR screens in the Common Mind Consortiums adult post-mortem DLPFC. **B.**

Number of individuals by diagnosis and number of estimated eGene perturbations (eGene perturbation assigned based on expression levels in the top or bottom 10%). **C.** Absolute proportion of individuals classified by number of eGenes perturbed colored by the number of synaptic (i), regulatory (ii), and multifunctional (iii) genes perturbed within a given individual. **D.** Differential gene expression of the 20 SCZ eGenes between schizophrenia cases (n=47) and (n=53) controls from pseudo-bulked cortical single-nuclei screen in the CMC DLPFC pulled from the PsychEncode Consortium's PsychScreen.

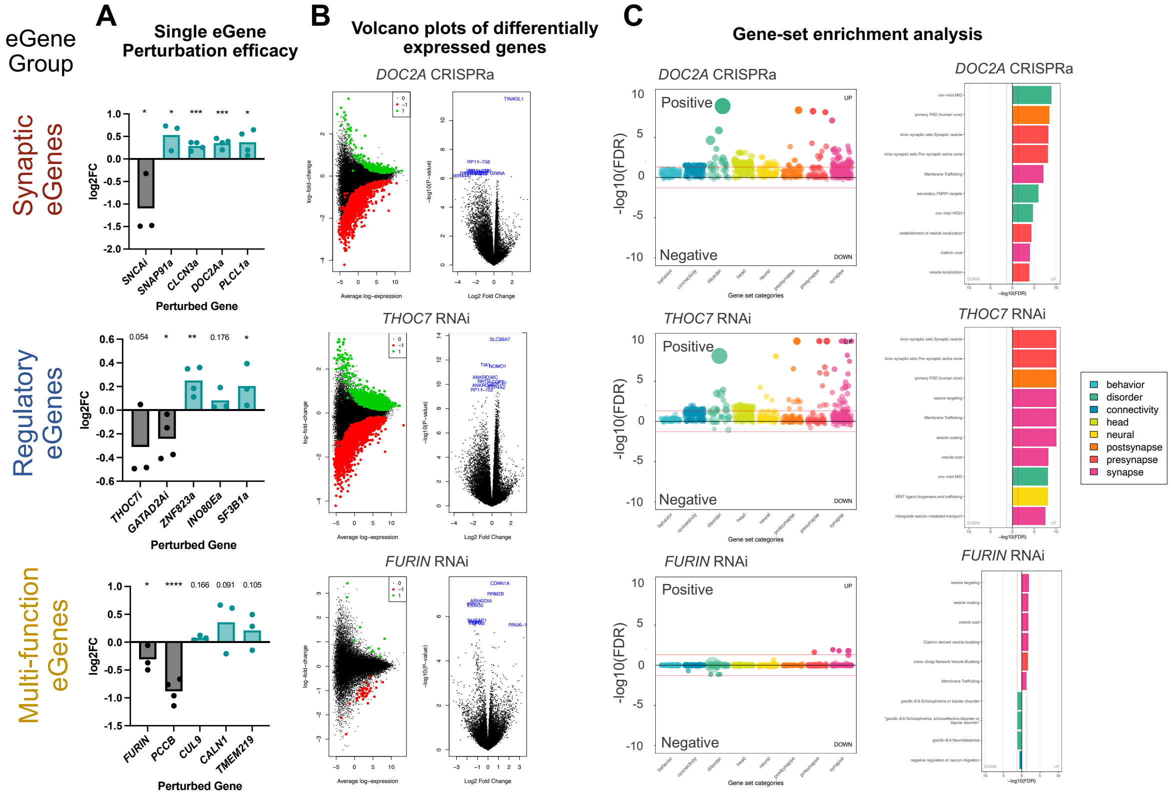


***Supplemental Figure 19. Perturbation of SCZ eGenes results in differential expression of genes relating to brain disorders and synaptic function (representative genes), related to Figure 1.***

**A.** Log2(fold change) of normalized counts data for target eGenes following single perturbations across all three eGene groups in D21 hiPSC-NPC derived iGLUTs. One tailed t-test, * = $p<0.05$; ** = $p<0.01$; *** = $p<0.001$; **** = $p<0.0001$. N = minimum of two donor lines with two replicates per condition, one differentiation. **B.** Volcano plots of differential gene expression in perturbation conditions relative to scramble control vectors. One representative eGene per group shown: *DOC2A* (synaptic), *THOC7* (regulatory) and *FURIN* (multi-function). Green dots designate significantly upregulated genes; red dots designate significantly downregulated genes (FDR $p < 0.05$). **C.** Gene set enrichment analysis (GSEA) across a collection of 698 manually curated neural themed gene-sets for the same three representative genes: *DOC2A* (synaptic), *THOC7* (regulatory) and *FURIN* (multi-function). Enrichments related to brain disorders (*DOC2A*, *THOC7*) and synaptic functions (*DOC2A*, *THOC7, FURIN*) were observed.

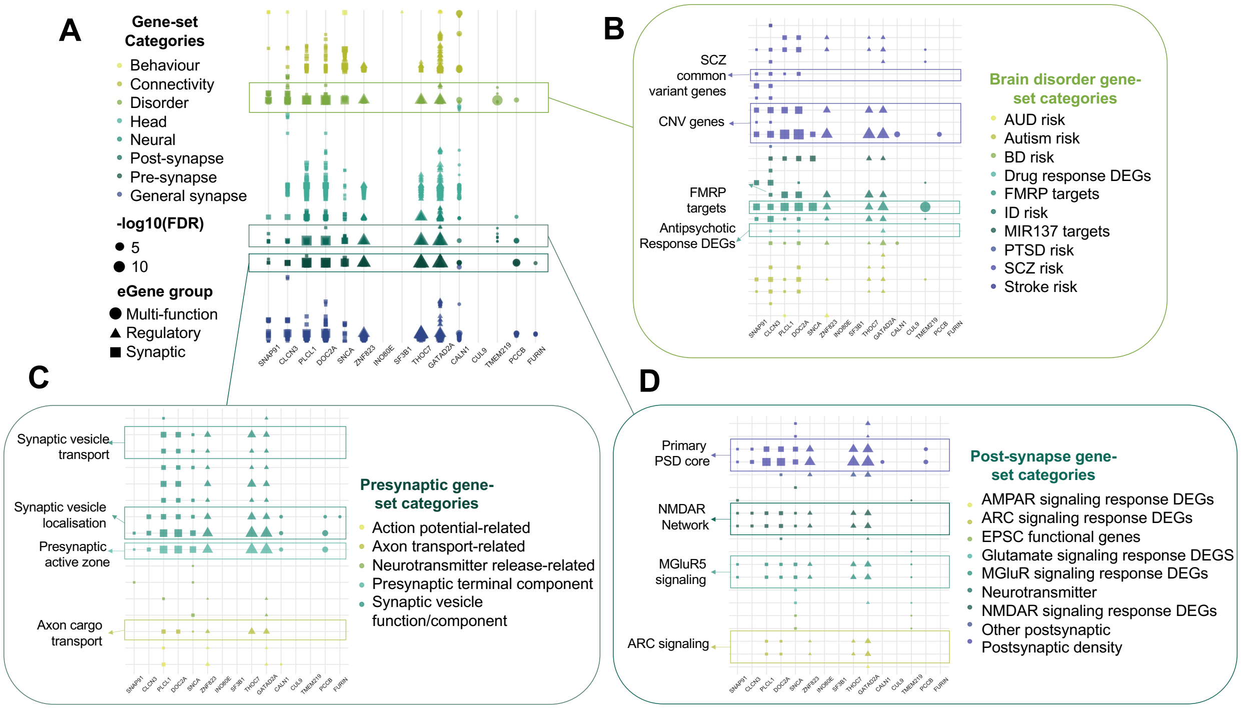


***Supplemental Figure 20. Perturbation of fifteen SCZ eGenes results in differential expression of genes relating to brain disorders and synaptic function (comprehensive analysis), related to Figure 1, SI figure 19.***

**A.** Gene set enrichment analysis (GSEA) across a collection of 698 manually curated gene-sets with a neural theme revealed enrichments of gene-sets related to brain disorders and synaptic functions across 12 out of 15 eGene perturbations. **B.** Across brain disorder-related gene-sets, 5 out of 5 synaptic eGene perturbations showed strong enrichment of other genes relating to SCZ risk, including both common variant-linked genes and copy number variant (CNV) genes associated with SCZ. **C.** Across presynaptic function gene-sets, 10 out of 15 eGene perturbations showed enrichment of genes relating to synaptic vesicle localization and transport. **D.** Across postsynaptic function gene-sets, 8 out of 15 eGene perturbations showed enrichment of genes relating to components of glutamatergic neurotransmission. Additional GSEA data for the arrayed screen using curated inflammatory and cell-death-related genesets can be found in **SI Data 1**.

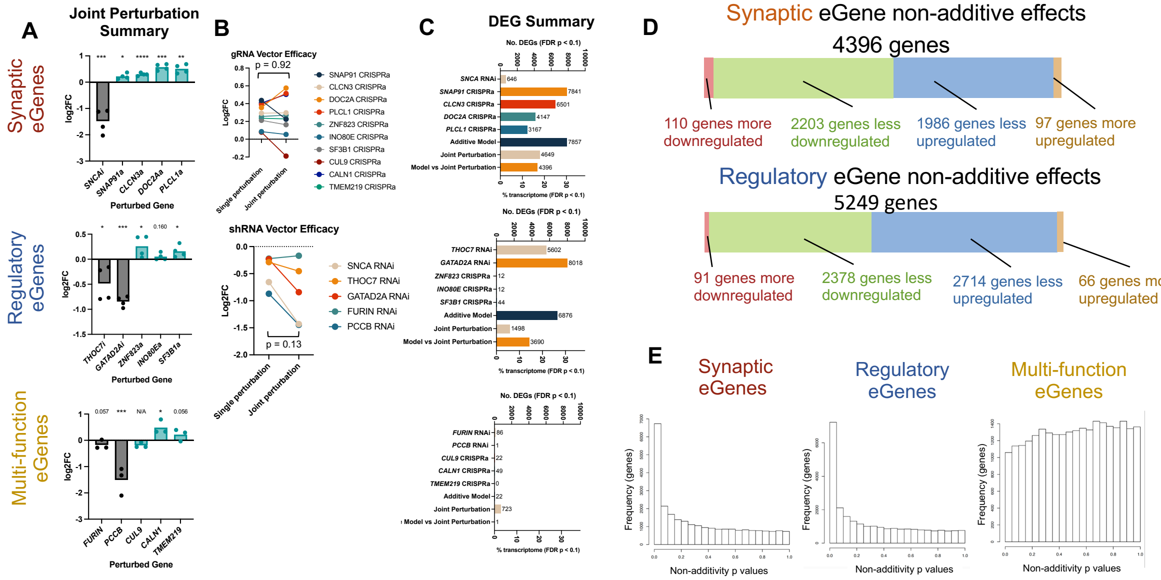


***Supplemental Figure 21. Combinatorial perturbation of regulatory eGenes results in non-additive effects on transcription, related to Figure 4.***

**A.** Log2(fold change) of normalized counts data for target eGenes following joint perturbations across eGene groups in D21 hiPSC-NPC derived iGLUTs. One tailed t-test, * = p<0.05; ** = p<0.01; *** = p<0.001; **** = p<0.0001. N = minimum of two donor lines with two replicates per condition, one differentiation. **B.** Comparison of gRNA and shRNA vector efficacy in single and joint eGene perturbations. The use of gRNA and shRNA vectors in joint perturbation conditions did not significantly reduce the magnitude of perturbation of target eGenes. Wilcoxon matched-pairs signed rank test, p = 0.125 (shRNA vectors), p = 0.922 (gRNA vectors). **C.** Summary of differentially expressed genes (DEGs) across the individual perturbations, additive model, combinatorial perturbation, and the additive-combinatorial comparison for each functional eGene group (FDR p<0.1). **D.** The majority (>95%) of non-additive genes in the Regulatory eGene group demonstrated significantly less differential expression in the measured combinatorial perturbation relative to the expected additive model (see Box 1 for an explanation of different categories of non-additive effects on transcription). **E.** Histograms of uncorrected non-additivity p values for each functional eGene set. Synaptic and regulatory eGene experiments resulted in a non-additivity p value distribution skewed to lower values, indicating significant non-additivity was present in these experiments. The same pattern was not observed for the multi-function eGene group, which showed an even distribution of non-additivity p values indicating a lack of non-additivity*.*

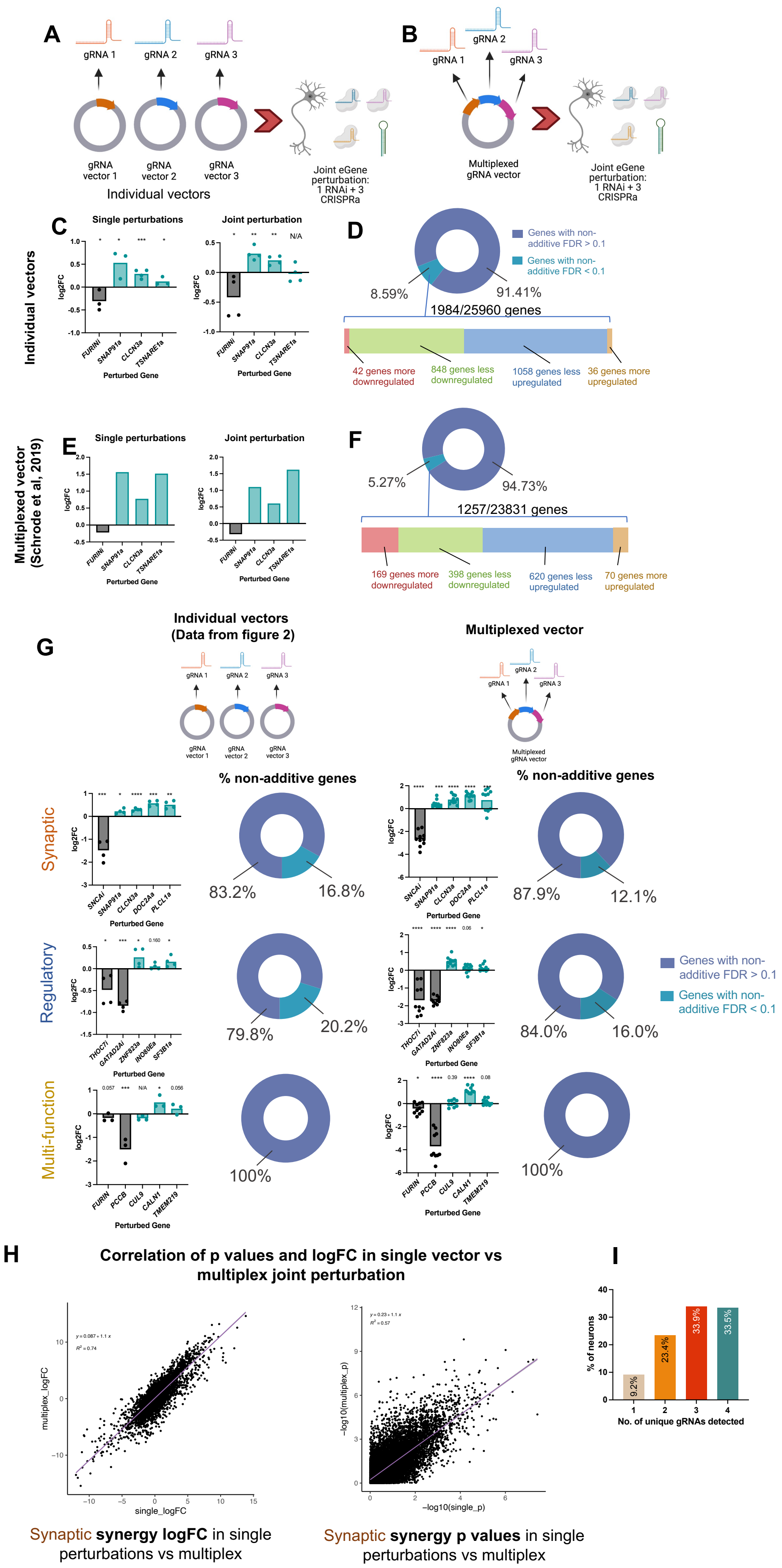

A
gRNA 1
gRNA 2
gRNA 3
gRNA vector 1
gRNA vector 2
gRNA vector 3
Individual vectors
Joint eGene perturbation: 1 RNAi + 3 CRISPRa
B
Multiplexed gRNA vector
Individual vectors
C
Single perturbations
Joint perturbation
log2FC
Perturbed Gene
FURINi
SNAP91a
CLCN3a
TSNARE1a
D
Genes with non-additive FDR > 0.1
Genes with non-additive FDR < 0.1
8.59%
91.41%
1984/25960 genes
42 genes more downregulated
848 genes less downregulated
1058 genes less upregulated
36 genes more upregulated
Multiplexed vector (Schrode et al, 2019)
E
F
5.27%
94.73%
1257/23831 genes
169 genes more downregulated
398 genes less downregulated
620 genes less upregulated
70 genes more upregulated
G
Individual vectors (Data from figure 2)
Multiplexed vector
% non-additive genes
Synaptic
83.2%
16.8%
87.9%
12.1%
SNCAi
DOC2Aa
PLCL1a
Regulatory
79.8%
20.2%
84.0%
16.0%
THOC7i
GATAD2Ai
ZNF823a
INO80Ea
SF3B1a
Multi-function
100%
FURIN
PCCB
CUL9
CALN1
TMEM219
H
Correlation of p values and logFC in single vector vs multiplex joint perturbation
multiplex_logFC
single_logFC
y = 0.087 + 1.1 x
$R^2$ = 0.74
−log10(multiplex_p)
−log10(single_p)
y = 0.23 + 1.1 x
$R^2$ = 0.57
Synaptic synergy logFC in single perturbations vs multiplex
Synaptic synergy p values in single perturbations vs multiplex
I
% of neurons
No. of unique gRNAs detected
9.2%
23.4%
33.9%
33.5%

***Supplemental Figure 22. Comparison of non-additive effects of joint perturbations using combined single vectors vs multiplexed vector, related to Figure 4.***

**A & B.** Four SCZ eGenes (*FURIN*, *SNAP91*, *CLCN3* and *TSNARE1*) were individually and jointly perturbed using gRNA and shRNA vectors, either using separate (**A**, here) or multiplexed (**B**, previous) vectors in an independent replication of the combinatorial eGene perturbation experiment conducted by Schrode *et al* (2019) using a multiplexed gRNA vector. **C.** Log2(fold change) of normalized counts data for four target eGenes following single (left) and joint (right) perturbations across in D21 hiPSC-NPC derived iGLUTs, using individual vectors. One-tailed t-test, * = p<0.05; ** = p<0.01; *** = p<0.001; **** = p<0.0001. **D.** Summary of non-additive effects of joint eGene perturbation using individual vectors across the transcriptome. 8.59% of genes demonstrated significant non-additive effects; most non-additive effects showed less differential expression following joint perturbation than predicted by the additive model (see **Box 1** for an explanation of different categories of non-additive effects on transcription). **E.** Log2(fold change) of eGenes following single and joint perturbations across in D21 hiPSC-NPC derived iGLUTs. RNA-seq data taken from Schrode *et al* (2019), using multiplexed vectors. **F.** Summary of non-additive effects of joint eGene perturbation using multiplexed vectors across the transcriptome. 5.27% of genes demonstrated significant non-additive effects. Again, the majority of non-additive effects showed less differential expression following joint perturbation than predicted by the additive model. RNA-seq data taken from Schrode *et al* (2019). **G.** Additional multiplexed, polycistronic gRNA vectors were generated expressing the same gRNA sequences targeting each of the three functional eGene sets. Each eGene set was jointly perturbed using the multiplexed vectors and differential gene expression was compared to the previously-generated additive models seen in **Figure 4**. Left: Log2(fold change) of normalized counts data for target eGenes following joint perturbations using individual vectors in D21 hiPSC-NPC derived iGLUTs (taken from **SI Fig 21**). Right: Log2(fold change) of normalized counts data for target eGenes following joint perturbations using multiplexed vectors in D21 hiPSC-NPC derived iGLUTs. One-tailed t-test, * = p<0.05; ** = p<0.01; *** = p<0.001; **** = p<0.0001. Right, pie charts: the proportion of the transcriptome showing significant non-additive effects (non-additive FDR p<0.1) following joint eGene perturbation using the multiplexed vectors was comparable to the non-additive effects seen after joint perturbation with individual vectors (left pie charts, taken from **Fig 4**). Multiplexed perturbation of synaptic and regulatory eGenes resulted in 12.1% and 16% of the transcriptome showing non-additive effects respectively, while multiplexed perturbation of the multi-function eGene group resulted in no detectable non-additive effects, similar to results using individual vectors. **H.** Comparison of non-additive logFC expression values and non-additive FDR p values following joint perturbation of 5 synaptic eGenes using multiplexed and individual vectors. Results generated using a multiplexed gRNA vector were highly correlated with those generated using individual vectors ($R^2$ = 0.77 for logFC values, $R^2$ = 0.57 for FDR p values across the transcriptome). **I.** Number of unique gRNAs detected in each cell following single cell sequencing and CRISPR detection of D21 hiPSC-NPC derived iGLUTs transduced with four gRNA vectors targeting synaptic eGenes as in **Figure 4**. A majority of unique gRNAs were detected in the majority of cells.

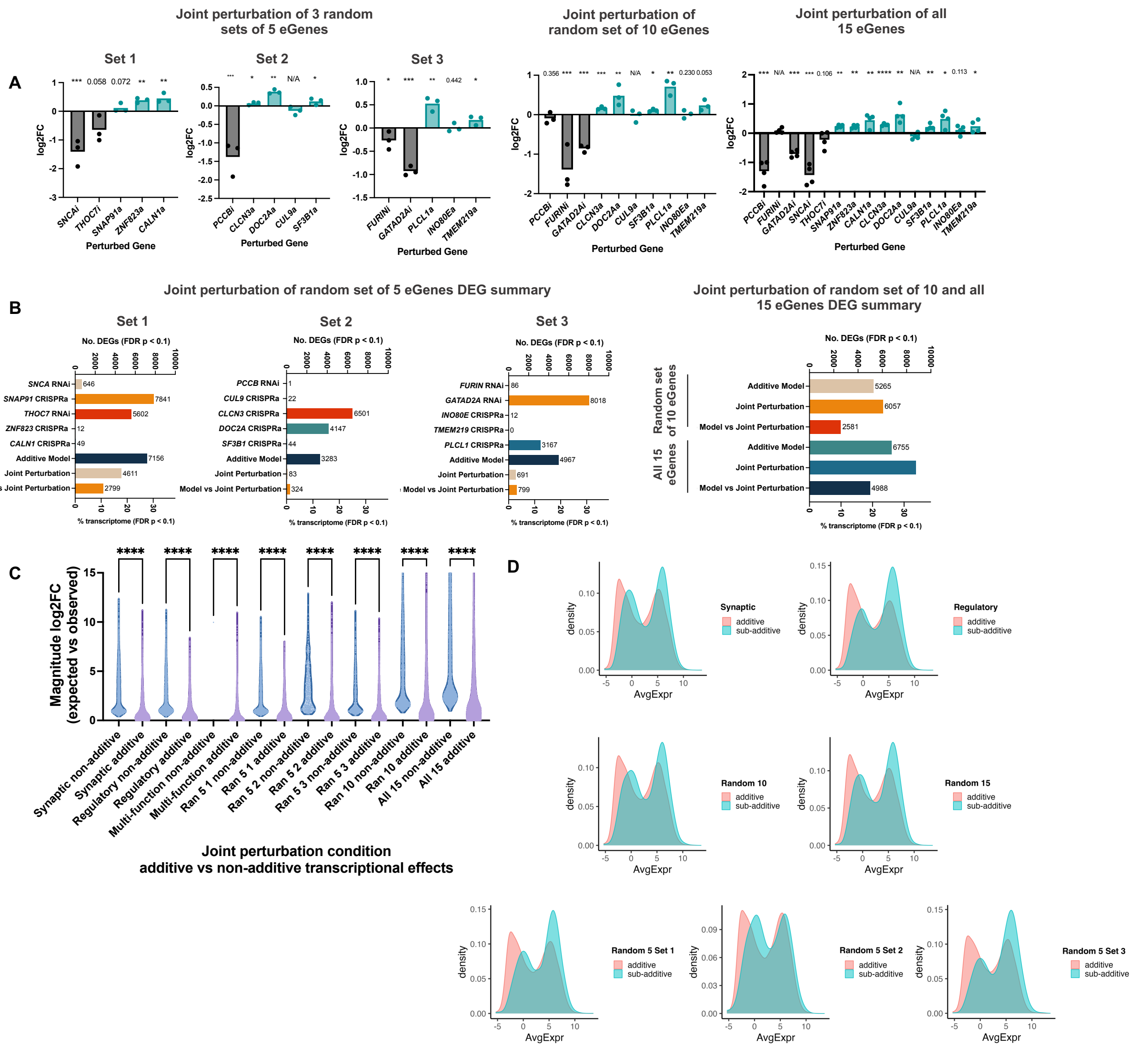


***Supplemental Figure 23. Combinatorial perturbation of random eGenes results in non-additive impacts on transcription which scale with the number of perturbed eGenes, related to Figure 4.***

Three groups of five eGenes and one group of ten eGenes were randomly selected from the full fifteen eGene list, as well as all fifteen eGenes. We label these sets as “5 random 1”, “5 random 2”, “5 random 3”, “10 random”, and “15 random”. These gene groups were manipulated alone and in combination, in parallel with the synaptic, regulatory, and multi-function groups. **A.** Log2(fold change) of normalized counts data for target eGenes following combinatorial perturbation of sets of five, ten, and fifteen eGenes randomly assigned from the synaptic, regulatory, and multi-function eGene groups. One-tailed t-test, * = p<0.05; ** = p<0.01; *** = p<0.001; **** = p<0.0001. **B.** Summary of differentially expressed genes (DEGs) across the additive model, combinatorial perturbation, and the additive-combinatorial comparison for each random eGene group (FDR p<0.1). **C.** Magnitude of log2(fold change) for additive and non-additive genes across all functional and random sets of eGenes. Genes designated as "non-additive" by our non-additivity analysis exhibited log2(fold change) values that were more than double those of "additive" genes (mean log2FC in synaptic set: 2.86 for non-additive genes, 1.16 for additive genes; regulatory set: 2.40 vs 1.09; multi-function set: 9.97 vs 0.84; random sets of 5 eGenes: 2.31 vs 1.07, 3.47 vs 1.17 and 2.47 vs 1.07; random set of 10 eGenes: 4.75 vs 1.96; all 15 eGenes: 6.42 vs 2.79). One way ANOVA with post-hoc Bonferonni correction for multiple comparisons, **** = p<0.0001. **D.** Expression histograms of genes classed as "additive" or "sub-additive" in different joint eGene perturbations: synaptic, regulatory, random combinations of 5, 10 and 15 eGenes. Expression distributions of additive and sub-additive genes largely overlapped in each comparison. AvgExp = Log2(counts per million) across all scramble control conditions in arrayed experiments.

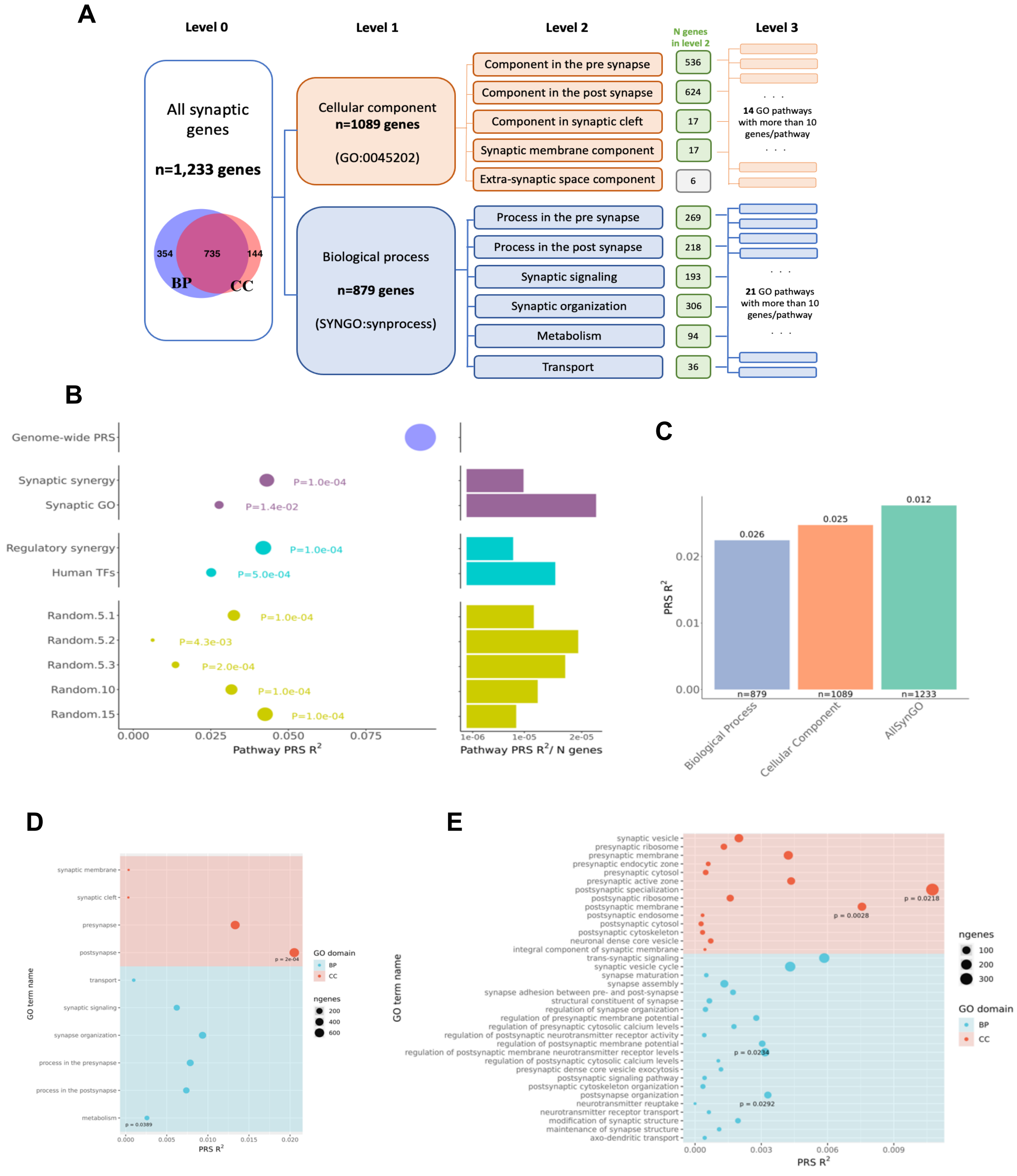


***Supplemental Figure 24. Within-pathway non-additive genes are associated with SCZ risk.***

**A.** Diagram depicting the structure of the Synaptic Gene Ontology and number of genes included in each gene set. Each box represents one gene-set used as pathway. **B.** Within-pathway non-additive genes are associated with SCZ risk**.** Seven groups of genes were defined to generate pathway PRS. Of those, five sets were obtained from our joint perturbation experiments and two sets from curated databases. For each pathway, genomic coordinates were extracted from GRCh37.75 GTF file to annotate SCZ risk variants to genes and pathways. Then, pathway specific PRS were calculated for each participant in a population-based cohort with genotype and SCZ case/control data, using summary statistics from the SCZ PGC-2 GWAS. Genome-wide PRS were calculated for comparison. Left: phenotypic variance explained (R2) by genome-wide (purple), synaptic (magenta), regulatory (blue) and random (green) pathway PRS. The size of the dot represents number of genes included in each pathway/gene set. Right: phenotypic variance explained (R2) normalized by the number of genes within each pathway. **C.** Results for pathway PRS of SYNGO gene-sets (Level 0 and Level 1). Number on top of the bar represents competitive P-value calculated using 10,000 permutations (See Methods). Number on bottom of the bar represents the number of genes included in the pathway. **D**. Results for pathway PRS of SYNGO gene-sets (Level 2). **E.** Results for pathway PRS of SYNGO gene-sets (Level 3). For panels c) and d), p indicated next to the dots correspond to competitive P-value calculated using 10,000 permutations.

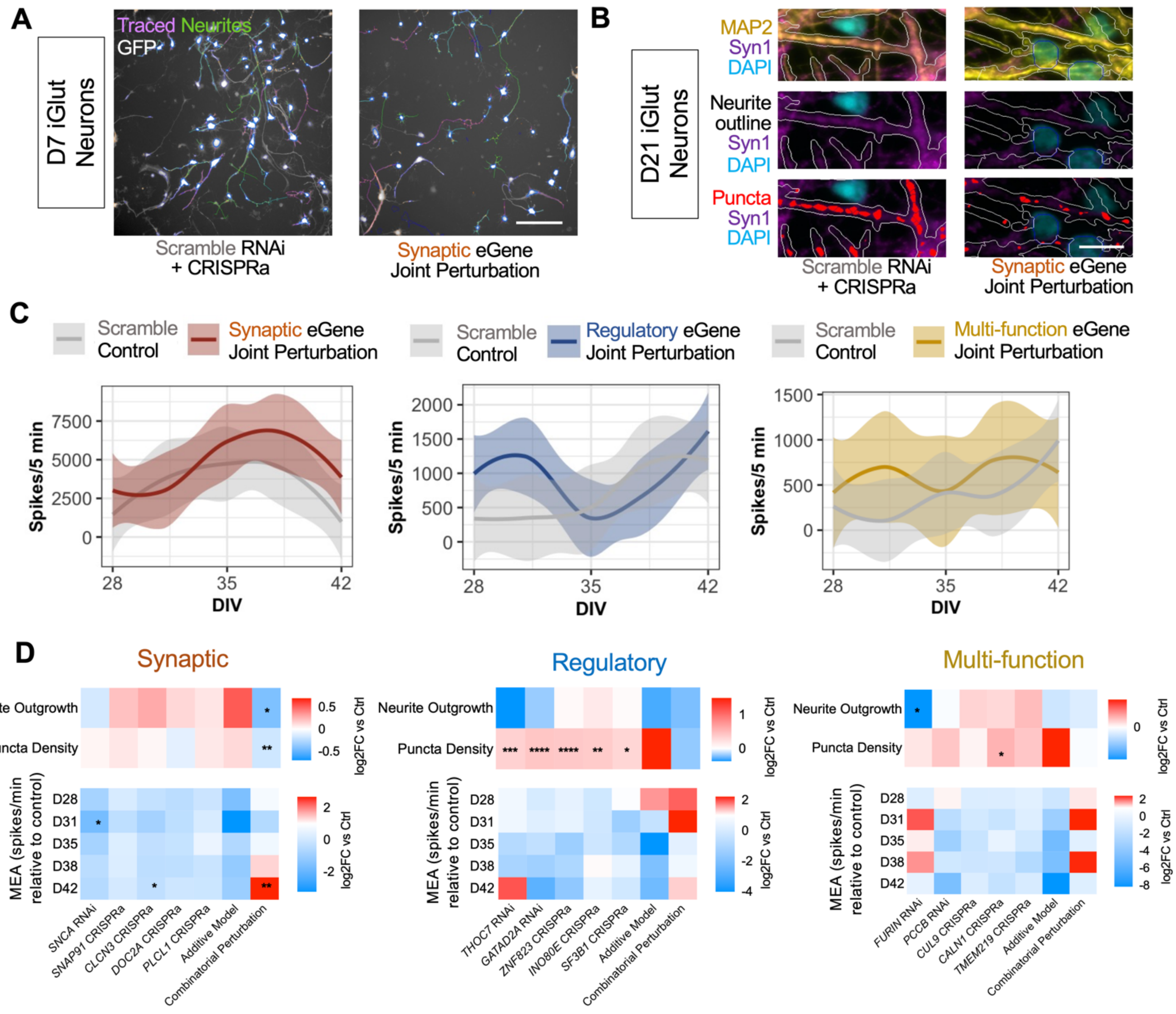


***Supplemental Figure 25. Combinatorial perturbation of SCZ eGenes with synaptic functions results in impaired neurite outgrowth, synaptic expression, and neuronal hyperactivity.***

**A.** Combinatorial perturbation of all five synaptic eGenes in D7 iGLUTs resulted in significant reduction of neurite outgrowth relative to a combinatorial scramble gRNA + shRNA control. **B.** Combinatorial perturbation of all five synaptic eGenes in D21 iGLUTs resulted in significant reduction of Syn1+ puncta density relative to a combinatorial scramble gRNA + shRNA control. **C.** LOESS plots; combinatorial perturbation of synaptic eGenes within results in transient neuronal hyperactivity in MEA recordings of D28-42 iGLUTs. **D.** Summary heatmap of neurite outgrowth, puncta density and neuronal activity data for within and across function eGene perturbations. N = minimum of 2 independent experiments across 2 donor lines with 10-12 technical replicates per condition. Neurite outgrowth, synaptic puncta density and MEA experiments were carried out separately using different batches of differentiated neurons from the same donor cell lines. Imaging data: one way ANOVA with post-hoc Bonferroni multiple comparisons test. MEA data: mixed effects model with post-hoc Bonferroni multiple comparisons tests across each perturbation-control comparison and time point. * = p<0.05; ** = p<0.01; *** = p<0.001; **** = p<0.0001.

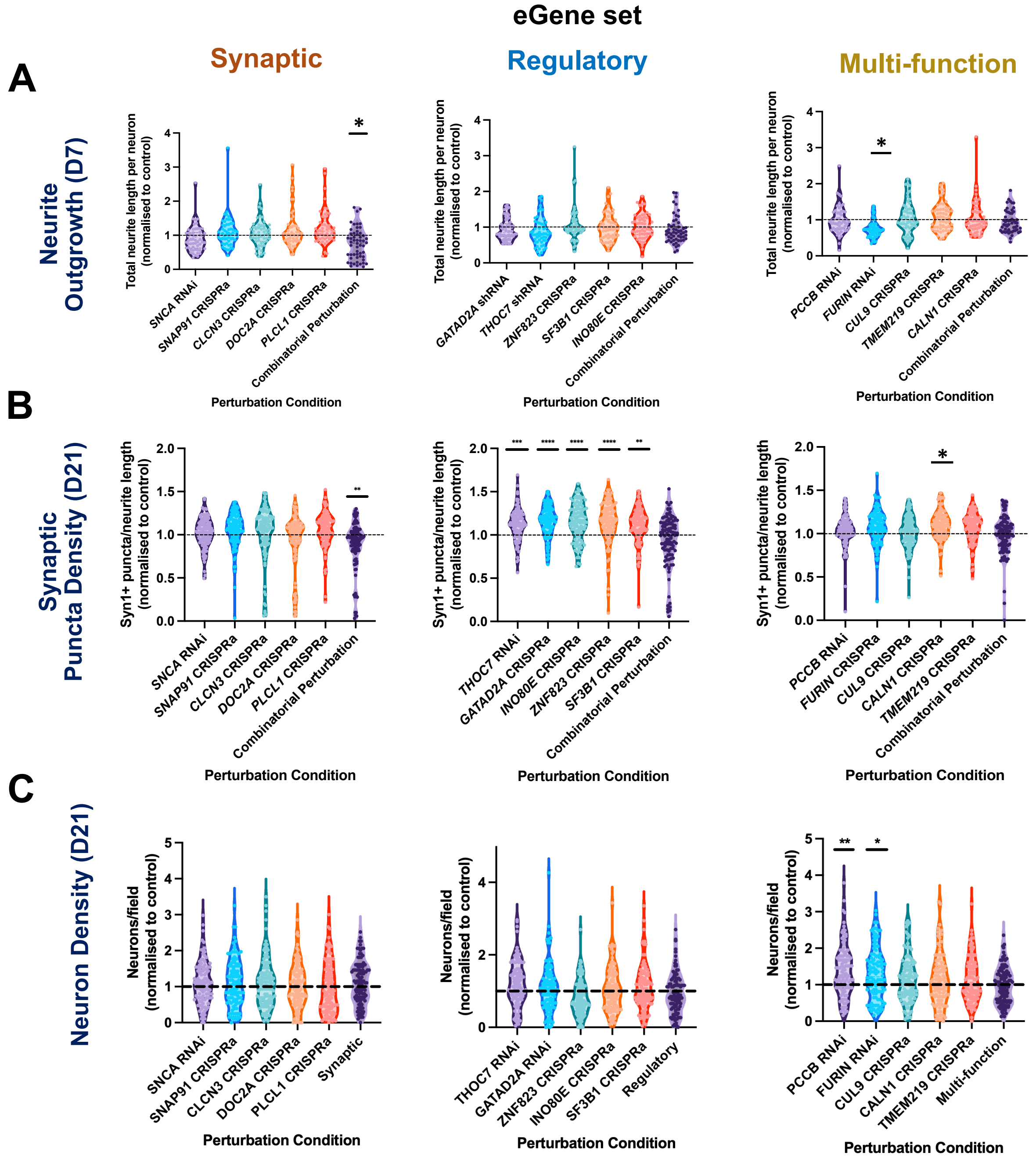


***Supplemental Figure 26. Combinatorial perturbation of SCZ eGenes with synaptic functions results in impaired neurite outgrowth and synaptic expression, related to SI Figure 25, Figure 4.***

**A**. Impact of single and joint eGene perturbations on early (D7) neurite outgrowth in hiPSC-derived iGLUTs across all three eGene groups. Combinatorial perturbation of all five synaptic eGenes and individual perturbation of FURIN (Multi-function eGene set) resulted in significant decreases in early neurite outgrowth relative to scramble control conditions. **B.** Impact of single and joint eGene perturbations on Synapsin1 (Syn1) +ve puncta expression in D21 hiPSC-derived iGLUTs across all three eGene groups. Syn1+ve puncta values are expressed relative to MAP2 +ve neurite length in each image. Combinatorial perturbation of all five synaptic eGenes resulted in significant decreases in synaptic puncta expression relative to scramble control conditions. Individual perturbation of regulatory eGenes and CALN1 (Multi-function eGene set) resulted in significant increases of Syn1+ puncta density relative to scramble controls. N = minimum of 2 independent experiments across 2 donor lines with 12 technical replicates per condition and 9 images analysed per replicate. One way ANOVA with post-hoc Bonferonni multiple comparisons test. * = $p<0.05$; ** = $p<0.01$; *** = $p<0.001$; **** = $p<0.0001$. **C.** Impact of single and joint eGene perturbations on neuron density (no. MAP2 +ve cell bodies per field, normalized relative to relevant control) in D21 hiPSC-derived iGLUTs across all three eGene groups. No significant reductions in neuronal density were seen in any eGene perturbation condition relative to control, indicating that eGene perturbations were not causing significant cell death. However, individual perturbations of the multi-function eGenes *FURIN* and *PCCB* were associated with modest but significant increases in neuronal density relative to scramble controls. N = minimum of 2 independent experiments across 2 donor lines with 12 technical replicates per condition and 9 images analysed per replicate. One way ANOVA with post-hoc Bonferonni multiple comparisons test. * = $p<0.05$; ** = $p<0.01$; *** = $p<0.001$; **** = $p<0.0001$.

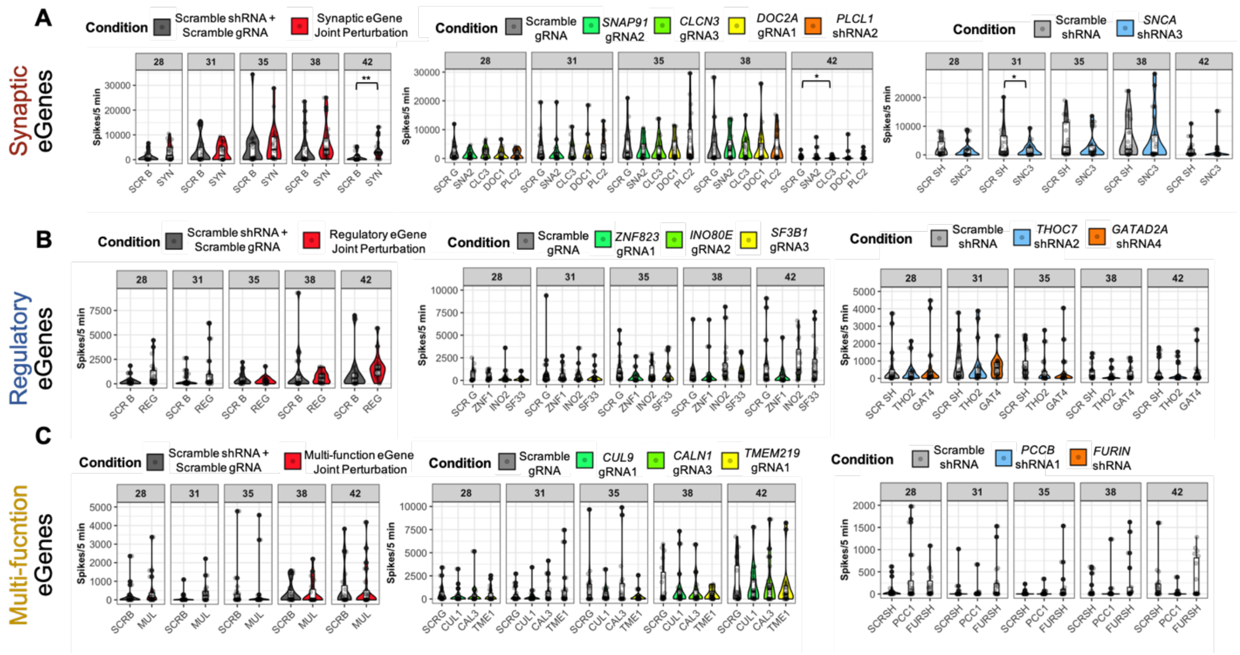

***Supplemental Figure 27. Combinatorial perturbation of SCZ eGenes with synaptic functions results in neuronal hyperactivity, related to SI fig 25, Figure 4.***

**A-C.** Impact of single and joint eGene perturbations on MEA-recorded spike traces in mature (4-6 week old) hiPSC-derived iGLUT/astrocyte co-cultures across all three eGene groups. **A.** Combinatorial perturbation of all five synaptic eGenes resulted in transient neuronal hyperactivity relative to a scramble control condition. **B.** Combinatorial perturbation of all five regulatory eGenes did not result in the same pattern of transient neuronal hyperactivity seen in the synaptic combinatorial eGene perturbation. **C.** Combinatorial perturbation of all five multi-function eGenes did not result in the same pattern of transient neuronal hyperactivity seen in the synaptic combinatorial eGene perturbation. Individual eGene perturbations did not significantly impact neuronal firing frequency. N = minimum of 2 independent experiments across 2 donor lines with 20 technical replicates per condition. Separate experiments were performed for each eGene set with NGN2 neurons from the same donor background. Separate mixed effects models were performed for each experiment, with post-hoc Bonferroni multiple comparisons tests across each perturbation-control comparison and time point. * = $p<0.05$; ** = $p<0.01$; *** = $p<0.001$; **** = $p<0.0001$.

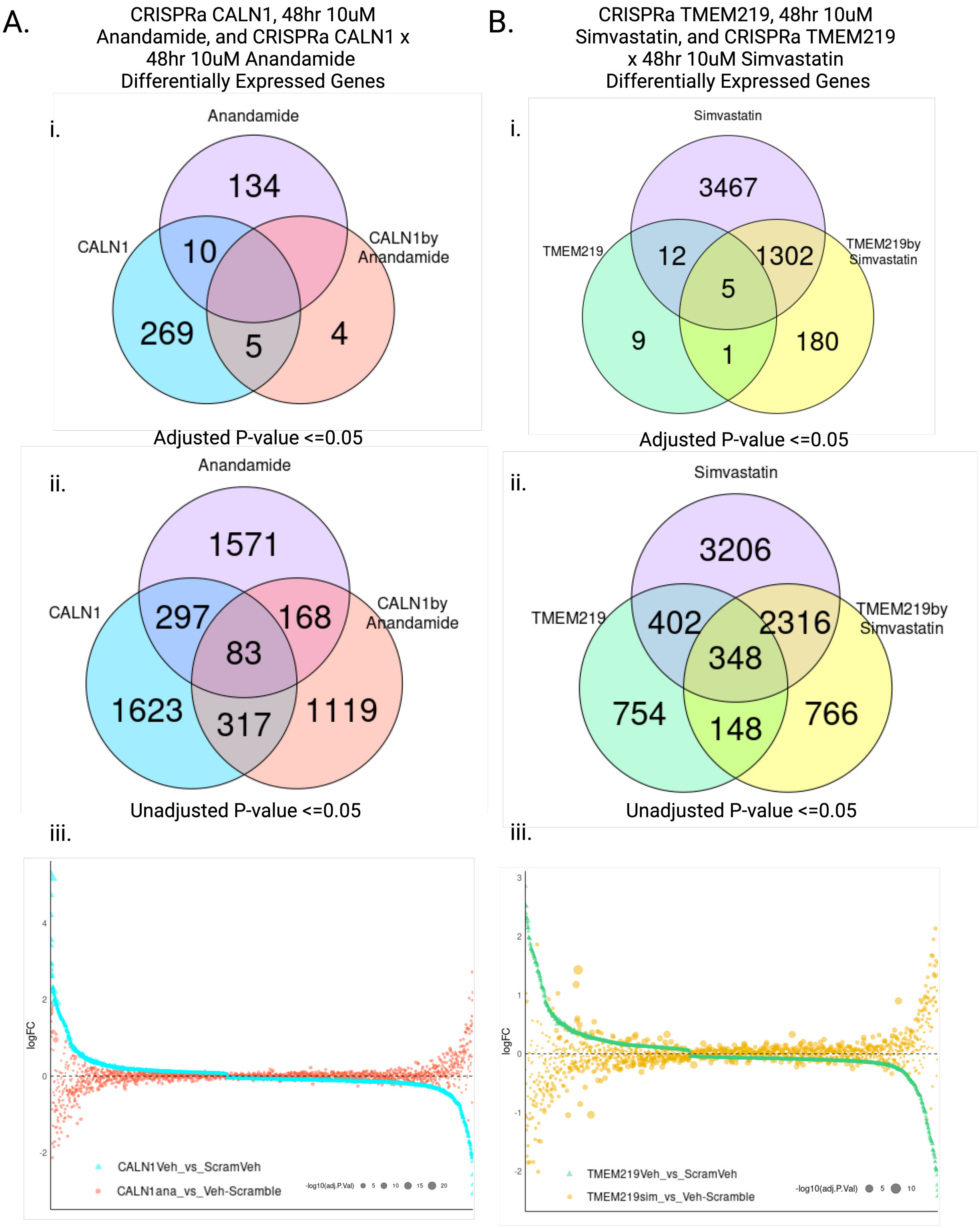
A.
CRISPRa CALN1, 48hr 10uM Anandamide, and CRISPRa CALN1 x 48hr 10uM Anandamide Differentially Expressed Genes
i.
Anandamide
CALN1
CALN1by Anandamide
134
10
269
5
4
Adjusted P-value <=0.05
ii.
Anandamide
CALN1
CALN1by Anandamide
1571
297
168
83
1623
317
1119
Unadjusted P-value <=0.05
iii.
logFC
CALN1Veh_vs_ScramVeh
CALN1ana_vs_Veh-Scramble
-log10(adj.P.Val) 5 10 15 20
B.
CRISPRa TMEM219, 48hr 10uM Simvastatin, and CRISPRa TMEM219 x 48hr 10uM Simvastatin Differentially Expressed Genes
i.
Simvastatin
TMEM219
TMEM219by Simvastatin
3467
12
1302
5
9
1
180
Adjusted P-value <=0.05
ii.
Simvastatin
TMEM219
TMEM219by Simvastatin
3206
402
2316
348
754
148
766
Unadjusted P-value <=0.05
iii.
logFC
TMEM219Veh_vs_ScramVeh
TMEM219sim_vs_Veh-Scramble
-log10(adj.P.Val) 5 10

***Supplementary Figure 28. In vitro validation of drug-eGene interactions at the transcriptomic level demonstrate an opposing effect of the predicted drugs on eGene activation, related to Figure 6.***

**A.** Treatment of cells with CRISPRa *CALN1*-gRNA and 10uM Anandamide over 48hrs opposes or suppresses the transcriptomic impact observed in CRISPRa *CALN1*-gRNA + Vehicle treated cells. Venn diagrams of significant DEGs at an **(i)** adjusted p-vale <= 0.05 and at an **(ii)** unadjusted p-value of <= 0.05. **(iii)** Dot plot demonstrating the logFC of each gene in either the *CALN1* + Vehicle (blue) or *CALN1* + 10uM Anandamide (orange) condition, order by degree of logFC in the *CALN1* + Vehicle treated cells. Size of the points corresponds to the -log10 (adjusted p-value) **B.** Treatment of cells with CRISPRa *TMEM219*-gRNA and 10uM Simvastatin over 48hrs opposes the transcriptomic impact observed in CRISPRa *CALN1*-gRNA + Vehicle treated cells. Venn diagram of significant DEGs at an **(i)** adjusted p-vale <= 0.05 and at an **(ii)** unadjusted p-value of <= 0.05. **(iii)** Dot plot demonstrating the logFC of each gene in either the *TMEM219* + Vehicle (green) or *TMEM219* + 10uM Simvastatin (yellow) condition, order by degree of logFC in the *TMEM219* + Vehicle treated cells. Size of the points corresponds to the -log10 (adjusted p-value)

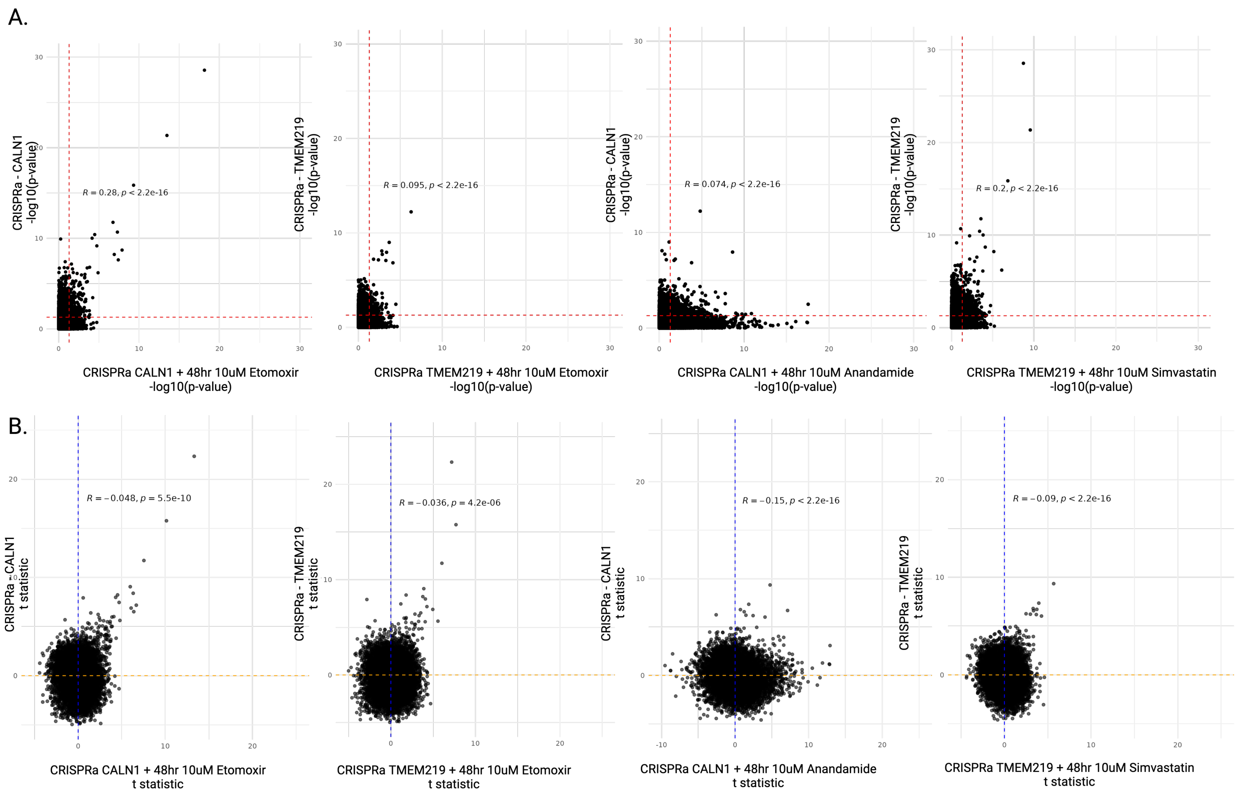


***Supplemental Figure 29. Comparison of the transcriptomic effect of eGene perturbations alone and the combined drug-eGene perturbations**, related to Figure 6.*

**A.** P-P plots comparing the -log10(p-value) of the target gRNA perturbation of *CALN1* CRISPRa or *TMEM219* CRISPRa only (y axis) compared to the interaction of the target gRNA perturbation and treatment with the predicted drug opposer (x-axis): *CALN1* CRISPRa + 48hr treatment with 10uM Etomoxir, *TMEM219* CRISPRa + 48hr treatment with 10uM Etomoxir, *CALN1* CRISPRa + 48hr treatment with 10uM Anadamide, *TMEM219* CRISPRa + 48hr treatment with 10uM Simvastatin. **B.** T-T plots comparing t-statistic of the target gRNA perturbation only (y axis) compared to the interaction of the target gRNA perturbation and treatment with the predicted drug opposer (x-axis).

A.

48hr 10uM Etomoxir

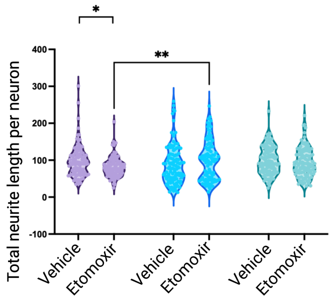


| Source of Variation | % of total variation | P value | P value summary | Significant? |
|---|---|---|---|---|
| Interaction | 1.476 | 0.0087 | ** | Yes |
| DRUG | 0.1279 | 0.3628 | ns | No |
| CRISPR | 0.4624 | 0.2242 | ns | No |

48hr 10uM Anandamide

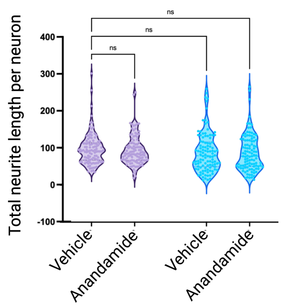


| Source of Variation | % of total variation | P value | P value summary | Significant? |
|---|---|---|---|---|
| Interaction | 1.476 | 0.0087 | ** | Yes |
| DRUG | 0.1279 | 0.3628 | ns | No |
| CRISPR | 0.4624 | 0.2242 | ns | No |

48hr 10uM Simvastatin

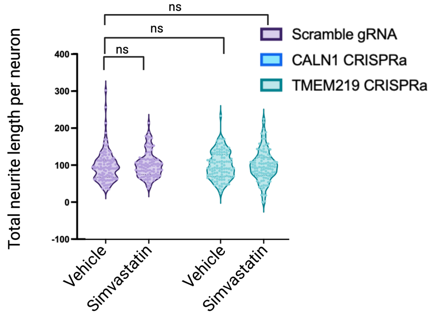


| Source of Variation | % of total variation | P value | P value summary | Significant? |
|---|---|---|---|---|
| Interaction | 0.09655 | 0.5273 | ns | No |
| DRUG | 0.02737 | 0.7364 | ns | No |
| CRISPR | 0.0005781 | 0.9610 | ns | No |

B.

48hr 10uM Etomoxir

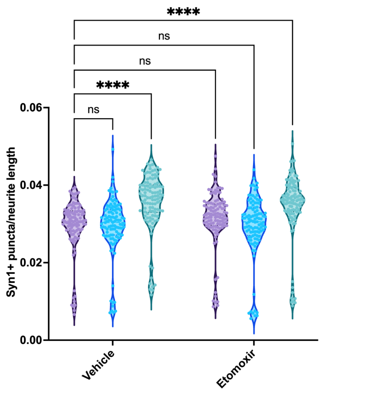


| ANOVA table | SS (Type III) | DF | MS | F (DFn, DFd) | P value |
|---|---|---|---|---|---|
| Interaction | 0.0001781 | 2 | 8.905e-005 | F (2, 641) = 1.545 | P=0.2140 |
| DRUG | 2.068e-005 | 1 | 2.068e-005 | F (1, 641) = 0.3589 | P=0.5493 |
| CRISPR | 0.004015 | 2 | 0.002007 | F (2, 641) = 34.84 | P<0.0001 |
| Residual | 0.03694 | 641 | 5.762e-005 | | |

48hr 10uM Anandamide

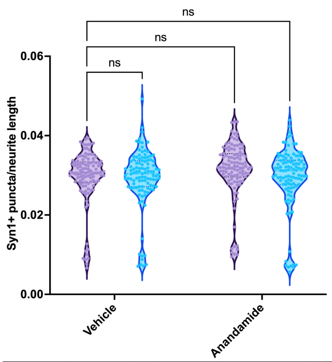


| ANOVA table | SS | DF | MS | F (DFn, DFd) | P value |
|---|---|---|---|---|---|
| Interaction | 8.578e-005 | 1 | 8.578e-005 | F (1, 428) = 1.549 | P=0.2140 |
| DRUG | 7.190e-005 | 1 | 7.190e-005 | F (1, 428) = 1.298 | P=0.2552 |
| CRISPR | 0.0001300 | 1 | 0.0001300 | F (1, 428) = 2.348 | P=0.1262 |
| Residual | 0.02370 | 428 | 5.538e-005 | | |

48hr 10uM Simvastatin

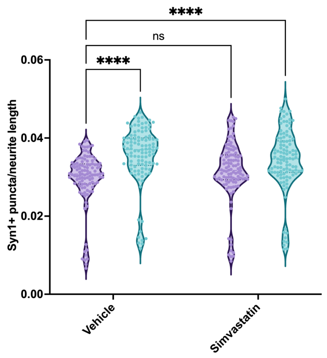


| ANOVA table | SS (Type III) | DF | MS | F (DFn, DFd) | P value |
|---|---|---|---|---|---|
| Interaction | 0.0002728 | 1 | 0.0002728 | F (1, 427) = 4.675 | P=0.0312 |
| DRUG | 8.758e-006 | 1 | 8.758e-006 | F (1, 427) = 0.1501 | P=0.6986 |
| CRISPR | 0.002020 | 1 | 0.002020 | F (1, 427) = 34.61 | P<0.0001 |
| Residual | 0.02492 | 427 | 5.835e-005 | | |

C.

48hr 10uM Etomoxir

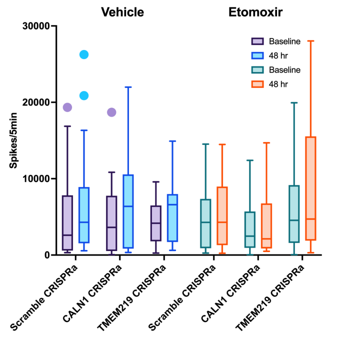


| Source of Variation | % of total variation | P value | P value summary |
|---|---|---|---|
| CRISPR | 0.8553 | 0.5851 | ns |
| DRUG | 0.1183 | 0.7002 | ns |
| treated vs untreated | 1.735 | <0.0001 | **** |
| CRISPR x DRUG | 2.534 | 0.2076 | ns |
| CRISPR x treated vs untreated | 0.09301 | 0.5954 | ns |
| DRUG x treated vs untreated | 0.1928 | 0.1446 | ns |
| CRISPR x DRUG x treated vs untreated | 0.1859 | 0.3565 | ns |

48hr 10uM Anandamide

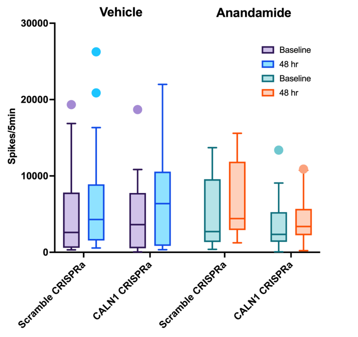


| Source of Variation | % of total variation | P value | P value summary |
|---|---|---|---|
| CRISPR | 0.7939 | 0.4621 | ns |
| DRUG | 1.181 | 0.3703 | ns |
| treated vs untreated | 2.276 | <0.0001 | **** |
| CRISPR x DRUG | 0.4832 | 0.5658 | ns |
| CRISPR x treated vs untreated | 0.0007388 | 0.9228 | ns |
| DRUG x treated vs untreated | 0.2335 | 0.0887 | ns |
| CRISPR x DRUG x treated vs untreated | 0.2354 | 0.0875 | ns |

48hr 10uM Simvastatin

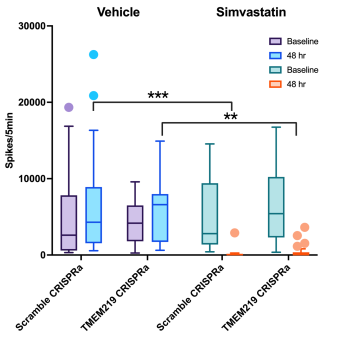


| Source of Variation | % of total variation | P value | P value summary |
|---|---|---|---|
| CRISPR | 0.004981 | 0.9396 | ns |
| DRUG | 6.952 | 0.0059 | ** |
| treated vs untreated | 3.271 | 0.0004 | *** |
| CRISPR x DRUG | 0.8854 | 0.3141 | ns |
| CRISPR x treated vs untreated | 0.09352 | 0.5277 | ns |
| DRUG x treated vs untreated | 11.93 | <0.0001 | **** |
| CRISPR x DRUG x treated vs untreated | 0.1302 | 0.4565 | ns |

***Supplementary Figure 30. In vitro validation of drug-eGene interactions on neuronal morphology, related to Figure 6.***

**A.** Effects of 48-hour treatment with etomoxir, anandamide, and simvastatin on neurite length in CRISPRa Scramble/*CALN1*/*TMEM219* gRNA transduced iGLUTs. **B.** Effects of 48-hour treatment with etomoxir, anandamide, and simvastatin on synaptic puncta density. **C.** Effects of 48-hour treatment with etomoxir, anandamide, and simvastatin on neuronal activity (MEA) in CRISPRa Scramble/*CALN1*/*TMEM219* gRNA transduced iGLUTs. A grouped three-way ANOVA found no significant differences in neuronal activity recorded as spikes per 5 minutes by Microelectrode Array (MEA) following 48hr treatment with etomoxir, anandamide, and simvastatin in CRISPRa perturbed or non-perturbed cells compared to baseline.

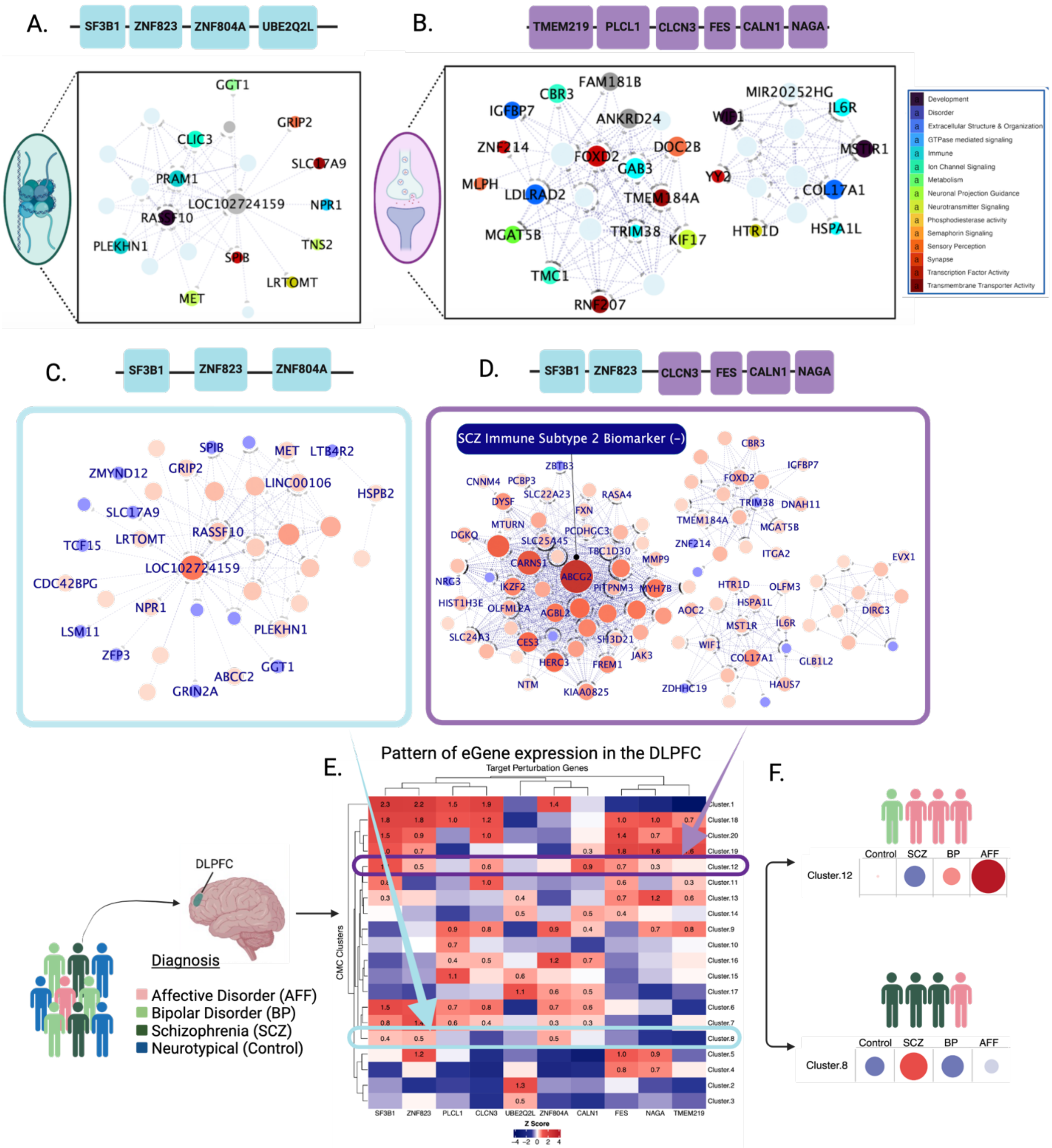


***Supplementary Figure 31.*** *Convergence across different eGenes resolve distinct networks that may inform molecular subtypes related to schizophrenia.* **A-B.** Unique convergent networks were resolved across **A.** regulatory (left) and **B.** signaling (right) SCZ eGenes, identifying node genes involved in synaptic signaling and immune response. Node color corresponds with general function with RNA genes colored in pale blue, while node size indicates node degree, and the line thickness represents weight of the connection (calculated as the percent duplication of the connection across bi-clustering runs). **C.** Convergent network resolved across three regulatory eGenes (*SF3B1*, *ZNF804A*, and *ZNF823*). **D.** Convergent network resolved across four signaling eGenes (*FES*, *NAGA*, *CALN1*, *CLCN3*) and two regulatory eGenes (*SF3B1* and *ZNF804A*). Nodes that are known targets of common or rare variants, drug targets, or known eGenes associated with neuropsychiatric disorders are labeled for both networks. Of note, the top node gene in the convergent network resolved across *FES*, *NAGA*, *CALN1*, *CLCN3*, *SF3B1* and *ZNF804A* is *ABCG2*, which is important in drug transport and blood-brain barrier permeability. Recently, relative under-expression of *ABCG2* in the blood of SCZ patients has been identified as a potential biomarker of SCZ immune type II. **(E)** Heatmap of average Z-scored gene expression of the 10 pooled CRISPRa target genes (columns) across 20 CMC clusters identified by K-means clustering (rows). **F.** Pearson's chi-squared test revealed significant differences in diagnosis status between clusters. SCZ diagnosis is significantly associated with increased expression of *SF3B1*, *ZNF804A*, and *ZNF823* (C) eGenes relative to the average Z-score in the DLPFC (cluster 8). AFF diagnoses is significantly associated with increased expression of these *FES*, *NAGA*, *CALN1*, *CLCN3*, *SF3B1* and *ZNF804A* in the DLPFC (cluster 12) (D). The size (larger = more positive) and color (blue = negative, red=positive) of circles in the matrix represent the chi-squared residuals for diagnoses of affective disorders (AFF), bipolar disorder (BP), and schizophrenia (SCZ) (rows) for each cluster (columns). Created with BioRender.com

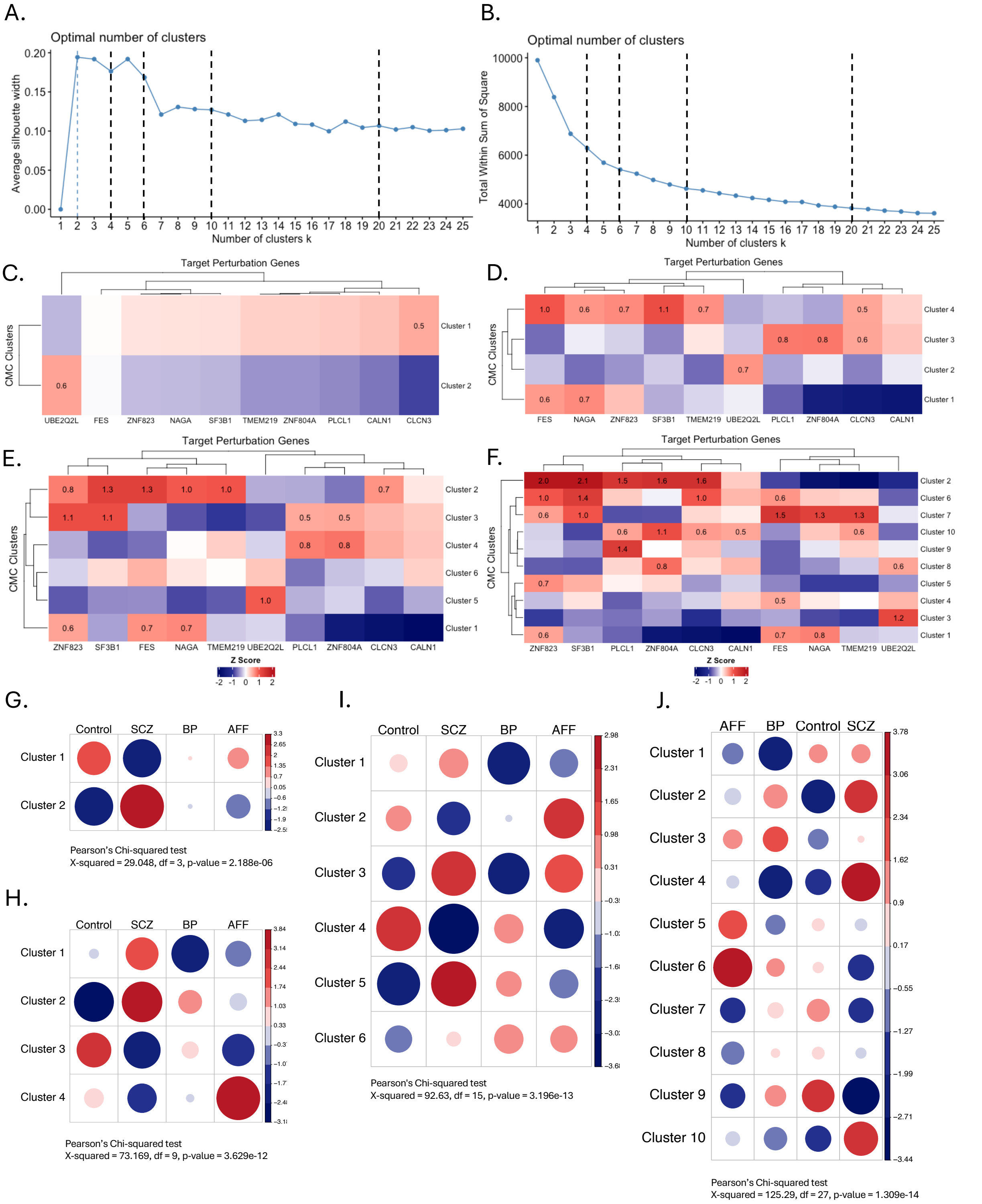


***Supplementary Figure 32.*** *K-means clustering resolution influences identification or diagnostic subgroups defined by eGene patterns of expression.* K means cluster optimization by silhouette analysis **(A)** and WSS optimization **(B)**. **(C-F)** Heatmaps of average Z-scored gene expression of the 10 pooled CRISPRa target genes (columns)

across 2, 4, 6, 10, and 20 CMC clusters identified by K-means clustering (rows). **(G-H)** Pearson's chi-squared test revealed significant differences in diagnosis status between clusters. For chi-squared plots, the size (larger = more positive) and color (blue = negative, red=positive) of circles in the matrix represent the chi-squared residuals for diagnoses of affective disorders (AFF), bipolar disorder (BP), and schizophrenia (SCZ) (columns) for each cluster (rows).
